# C-Reactive Protein Drives Potent Clearance of Blood Bacteria in the Liver by Activating the Complement System

**DOI:** 10.1101/2024.10.05.616753

**Authors:** Danyu Chen, Jiao Hu, Mengran Zhu, Yufeng Xie, Hantian Yao, Haoran An, Yumin Meng, Juanjuan Wang, Xueting Huang, Zhujun Shao, Ye Xiang, Jianxun Qi, George Fu Gao, Jing-Ren Zhang

**Affiliations:** Center for Infection Biology, School of Medicine, Tsinghua University, Beijing 100084, China; CAS Key Laboratory of Pathogen Microbiology and Immunology, Institute of Microbiology, Chinese Academy of Sciences, Beijing 100101, China; University of Chinese Academy of Sciences, Beijing 100049, China; Institute of Medical Technology, Peking University Health Science Center, Beijing 100191, China; Tsinghua-Peking Center for Life Sciences, Tsinghua University, Beijing 100084, China; State Key Laboratory for Infectious Disease Prevention and Control, National Institute for Communicable Disease Control and Prevention, Chinese Center for Disease Control and Prevention, Beijing, China

**Keywords:** C-reactive protein, capsule, capsular polysaccharide, encapsulated bacteria, septic infection, Kupffer cell, liver, complement system, C3, and complement receptor

## Abstract

Plasma C-reactive protein (CRP) is widely used as a biomarker for bacterial infections due to its massive induction during infections, however, the precise function of CRP in bacterial infections remains undefined. Here we show that CRP enables Kupffer cells (liver macrophages) to capture and eliminate a wide range of encapsulated bacteria from the bloodstream of mice and thereby provides rapid and effective immunity. Mechanistically, CRP binds to the structurally diverse capsular polysaccharides of major Gram-positive and -negative pathogens, and thereby activates complement C3 at the bacterial surface. The C3-opsonized microbes are in turn captured by C3 receptors on the surface of Kupffer cells, and eliminated in the liver sinusoids. Since CRP principally shares the functional features of antibodies in pathogen recognition/execution, CRP-based defense combines the broad spectrum of the innate immunity with the swiftness, potency and specificity of the adaptive immunity, which helps explain massive rise of CRP during systemic bacterial infections.

## INTRODUCTION

C-reactive protein (CRP) is an acute phase plasma protein named from its binding to the cell wall C-polysaccharide or phosphocholine (PC) of *Streptococcus pneumoniae* (pneumococcus).^1^ PC was later found to be a constituent of other bacterial polysaccharides.^2–5^ Human CRP is present at a trace level in healthy individuals, but rapidly rises by as much as 1,000 folds during severe bacterial infections and other inflammatory conditions, making it a common biomarker for clinical diagnosis of systemic infections and inflammation.^6^ Surprisingly, the biological function of CRP has remained largely speculative for nearly a century since its first identification in 1930.^7^ A polymorphism in the promoter region of the CRP gene is associated with the increased mortality in pneumococcal bacteremia patients.^8^ Passive administration or transgenic expression of human CRP in mice has been reported to protect mice from septic infection of *S. pneumoniae*,^9–13^ but the combined data demonstrate a modest immunity. Consistently, CRP binding to pneumococcal cell wall does not stimulate effective phagocytic killing *in vitro*.^14^

Plasma CRP is a disc-shaped homo pentameric protein, in which each protomer contains two calciums.^15^ While PC is the best-defined ligand of CRP, many autologous PC-free targets of CRP have been described, such as complement component 1q (C1q) and Fcγ receptors.^16–19^ Structural studies have revealed that the CRP has two functional faces: the activating face (A face) and ligand binding face (B face).^20,21^ Under the *in vitro* conditions, CRP interaction with PC at the B face leads to the subsequent C1q binding at the A face and activation of the complement system through the classical pathway.^19,22^ Mouse CRP shares 71% amino acid homology with the human ortholog, but is stably expressed at low levels even under infection conditions.^23^ Human CRP has been shown to activate mouse complement system, which is attributed to human CRP-mediated protection in mouse models of pneumococcal bacteremia.^24,25^ In contrast, CRP-binding to Fcγ receptors does not contribute to protective effect of CRP against pneumococcal infection.^24^

Encapsulated bacteria cause a series of devastating invasive diseases in humans, such as pneumonia, sepsis and meningitis.^26,27^ Capsules are known for their special physical and antiphagocytic properties.^28–30^ Nearly all of the capsules consist of capsular polysaccharides (CPS).^27,31^ Many pathogens produce large numbers of capsule types that differ in structure and antigenicity.^27^ More than 100 capsule serotypes have been documented for human pathogens *S. pneumoniae* and *Klebsiella pneumoniae.*^32,33^ Capsule types are clinically associated with disease potentials of encapsulated bacteria. The low-numbered pneumococcal serotypes/serogroups were dominantly identified in fatal pneumonia cases in pre-antibiotic era;^34^ *Haemophilus influenzae* type b is responsible for the vast majority of the childhood infections among the six serotypes.^35^

Kupffer cells (KCs), the liver resident macrophages embedded in the liver sinusoids, represent approximately 90% of total tissue macrophages in the body.^36^ KCs are highly capable of clearing blood-borne bacteria.^37–44^ Our recent studies have uncovered that the polysaccharide capsules, the outmost bacterial structure, shield encapsulated bacteria from KC capture in the liver sinusoids.^45,46^ Consistent with clinical association between capsule serotype and disease potential, certain capsule types are more capable of fending of KC recognition than the others in mouse sepsis model.^45,46^ The high-virulence (HV) types escape KC capture and thereby replicate in the blood, resulting in severe bacteremia and septic death. To less extents, the low-virulence (LV) types partially escape the hepatic interception and only cause fatal infection when KC-mediated anti-bacterial “firewall” is compromised or overwhelmed.^45,46^ The capture of the LV capsule types by KCs indicates the existence of host receptors for these capsules. In sharp contrast to massive numbers of capsule variants associated with human diseases, the asialoglycoprotein receptor (ASGR) is the only known capsule receptor on KCs (recognizing seroyte-7F and -14 capsules of *S. pneumoniae*).^45^ It remains completely unclear how KCs recognize many other LV capsules.

This study sought to identify host receptors that enable KCs to capture the LV bacteria in the liver. human and mouse plasma CRPs were found to broadly recognize a wide range of capsular serotypes of Gram-positive and -negative bacteria. The receptor-ligand interactions enable liver KCs to capture and kill blood-borne bacteria, providing potent immunity against otherwise fatal infections.

## RESULTS

### CRP binds to capsular polysaccharide of serotype-23F *S. pneumoniae*

To understand how the liver executes the clearance of blood-borne bacteria, we sought to identify the receptor(s) recognizing the capsule of serotype-23F *S. pneumoniae* (*Sp*23F), one of the most predominant serotypes causing the childhood invasive infections.^47^ Our previous study has shown that *Sp*23F is rapidly captured by KCs into the liver of mice.^45^ Our initial experiment showed that the hepatic immunity is blocked by free capsular polysaccharide of *Sp*23F (CPS23F) (**Figure 1A**). Consistently, the freshly isolated KCs from mouse liver showed significant adherence to LV *Sp*23F with normal mouse serum, but were poorly adherent to HV *Sp*8 (**Figure S1B**). We enriched CPS23F-binding protein(s) by affinity pulldown after incubation of CPS23F-coated beads with membrane protein-enriched fraction of mouse liver nonparenchymal cells (NPCs) in the presence of 10% mouse serum (**Figure 1B**). CPS of the HV serotype-8 *S. pneumoniae* (CPS8) was used as a negative control due to its poor binding to KCs.^45^ Serum was included since circulating bacteria are soaked in the blood before being captured by KCs. Proteomic analysis identified a total of 725 significantly enriched proteins by CPS23F, with CRP being the top hit (**Figure 1C** and **Table S2**). We verified the interaction between CPS23F and CRP using the recombinant mouse CRP (r-mCRP) (**Figures 1D** and **S1C**). r-mCRP showed a dose-dependent binding to immobilized CPS23F, but not CPS8. The interaction was competitively blocked by free CPS23F, validating the binding specificity (**Figure 1E**). Furthermore, fluorescence microscopy revealed r-mCRP binding to the live serotype-23F but not serotype-8 pneumococci (**Figure 1F**). These results demonstrated that CRP specifically binds to the capsule of serotype-23F *S. pneumoniae*.

**Figure 1.**
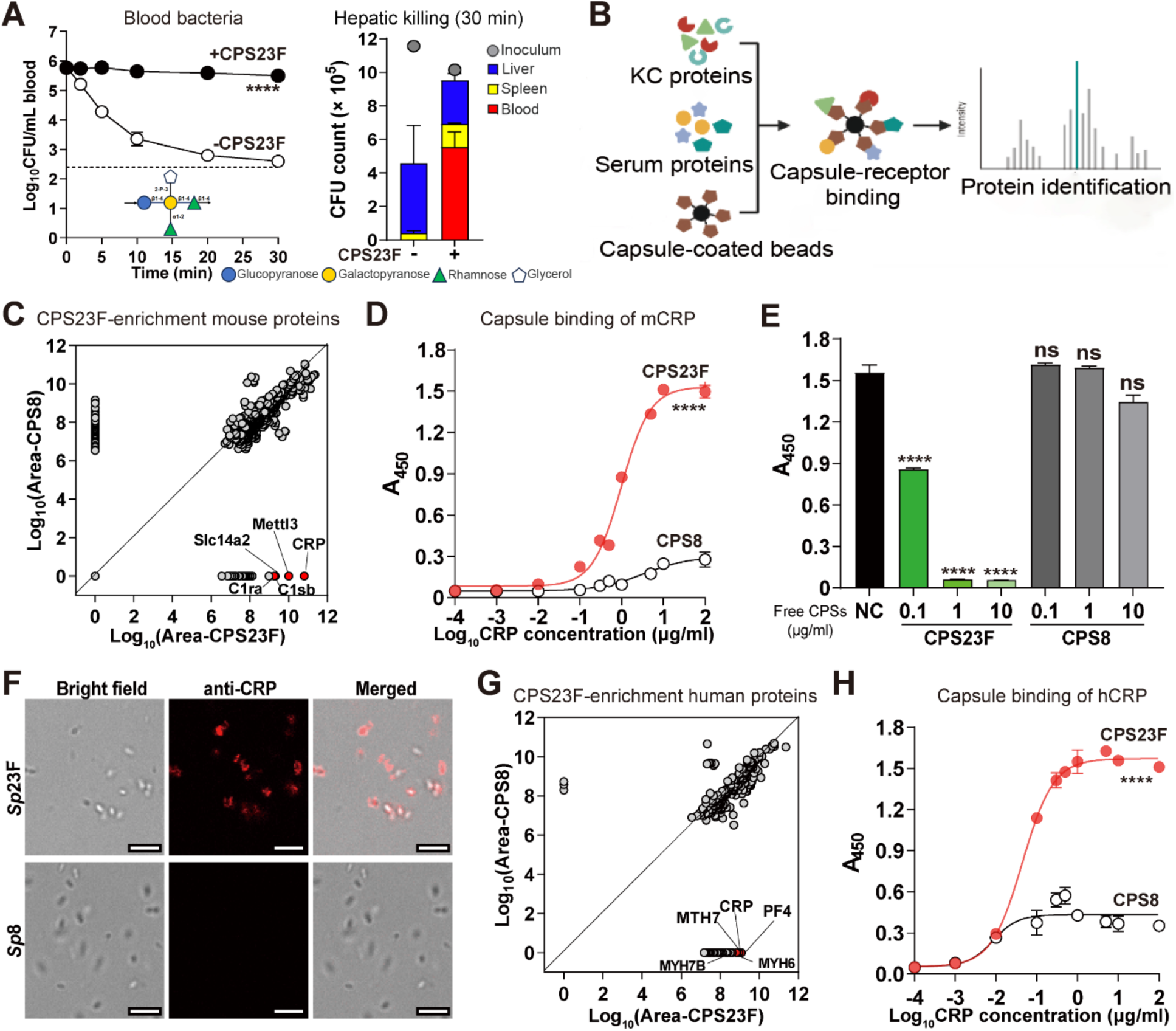
Identification of CRP as a CPS23F-binding protein. **A.** Inhibition against hepatic clearance of serotype-23F *S. pneumoniae* by free CPS23F. Mice were i.v. inoculated with 400 μg free CPS23F prior to i.v. infection and used to assess bacterial clearance left panel) and hepatic killing (right panel). n = 3; dotted line, detection limit. The inoculum of each group is indicated with a filled circle. **B.** Illustration of the experimental procedure. Capsule of pneumococcal serotype 23F (CPS23F) was coated to latex beads to co-incubate with membrane proteins of mouse liver NPCs in the presence of 10% mouse serum. **C.** CPS23F-enriched mouse proteins identified by mass spectrometry are plotted against the levels of the same proteins in the CPS8 group; CRP and other four top hits specified. **D.** CPS23F binding of mouse CRP (mCRP) was detected by ELISA with CPS23-coated wells and recombinant mCRP, and is presented as A_450 nm_ values. n = 3. **E.** Competitive inhibition of CRP-CPS23F binding by free CPS23F was determined by co-incubating free CSP23F with r-mCRP before being added to the CPS23F-coated wells; CPS-bound CRP detected as in (D). n = 3. **F.** mCRP binding to *Sp*23F pneumococci was detected by immunofluorescence microscopy with r-mCRP and anti-mCRP antibody. Bacterium-bound CRP was visualized with AF647-conjugated secondary antibody (red). Scale bar, 5 μm. **G.** CPS23F-enriched human proteins were enriched and identified as in (C). **H.** CPS23F binding of human CRP (hCRP) was detected as in (D). n = 3.

We performed a similar affinity pulldown with normal human serum and CPS23F-conjugated beads, and identified human plasma CRP (hCRP) as one of the top hits (**Figure 1G** and **Table S5**). The ELISA result verified that recombinant hCRP (r-hCRP) specifically bind to CPS23F in a dose-dependent manner (**Figure 1H**). The common capsule-binding activity of human and mouse CRPs is consistent with the high level of sequence conservation between the two proteins. Human and mouse CRPs have a 70.9% amino acid sequence identity (**Figure S7A**).

### CRP enables Kupffer cells to capture blood-borne *Sp*23F in the liver

We next determined the functional contribution of CRP against *Sp*23F infection by comparing the early clearance of *Sp*23F between wildtype (WT) and CRP-deficient (*Crp*^-/-^) mice in the first 30 min post i.v. inoculation of 10^6^ CFU bacteria. While wild type (WT) mice rapidly cleared *Sp*23F with a 50% clearance time (CT_50_) of 1.3 min, *Crp*^-/-^ mice fully retained the inoculum in the circulation (**Figure 2A**). By comparison, *Crp*^-/-^ and WT mice were equally able to clear serotype-14 pneumococci (*Sp*14), which is known to be recognized by ASGR on KCs.^45^ Since the liver is the major organ to trap blood-borne *Sp*23F and other LV pneumococci,^45^ we tested the impact of CRP deficiency on hepatic capture of *Sp*23F. The livers of WT mice contained 72.9% of the inoculum at 5 min post infection and the infected WT mice have only a residual level of bacteremia, whereas only 3.0% of the inoculated bacteria accumulated in the livers of *Crp*^-/-^ mice, with the vast majority of the bacteria in the blood (**Figure 2B**). By comparison, both *Crp*^-/-^ and WT mice effectively trapped *Sp*14 bacteria in the liver at this time point. Further analysis revealed dramatic reduction of bacteria in the blood, spleen, and liver of WT mice at 30 min post infection, but *Crp*^-/-^ mice failed to clear *Sp*23F, resulting in substantial increase in the total pneumococci (134.9% of the inoculum). The non-CRP binding *Sp*14 bacteria were similarly eliminated in WT and *Crp*^-/-^ mice. Furthermore, passive i.v. administration of 1 µg r-mCRP fully enabled *Crp*^-/-^ mice to rapidly capture *Sp*23F by the liver and clear bacteria from the circulation (**Figure 2C**). These data demonstrated that CRP is necessary and sufficient for serotype-specific hepatic capture and killing of serotype-23F pneumococci.

**Figure 2.**
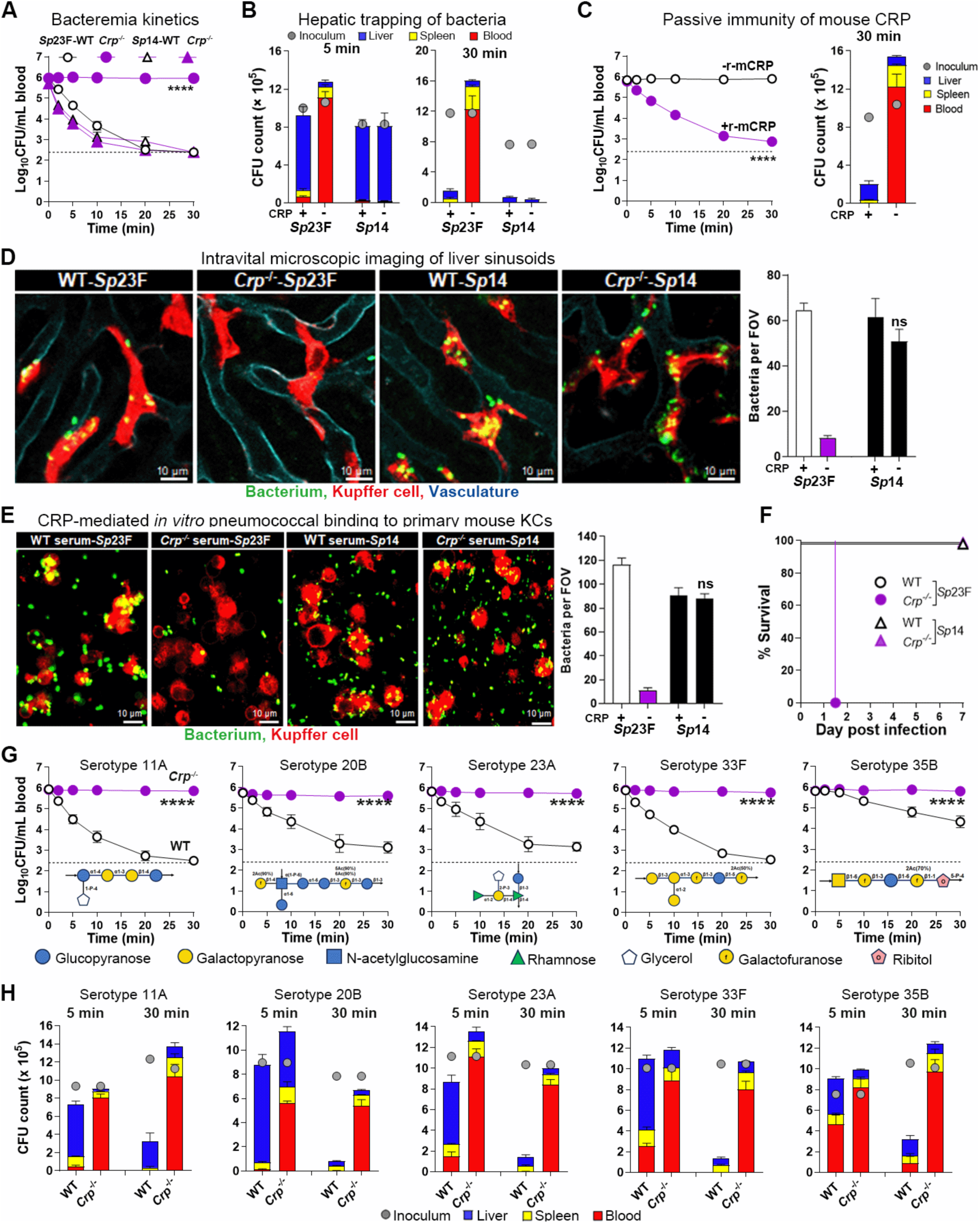
CRP-activated hepatic capture and killing of diverse *S. pneumoniae* serotypes. **A.** The importance of CRP in clearing serotype-23F *S. pneumoniae* (*Sp*23F) was determined by blood CFU plating of *Crp^-/-^* and WT mice at various time points post i.v. infection. n = 5; dotted line, detection limit. **B.** The contribution of CRP to hepatic trapping of *Sp*23F was characterized by CFU plating of the blood, liver and spleen of mice at 5 and 30 minutes post infection as in (A). The inoculum of each group is indicated with a filled circle. n = 5. **C.** The complementation of CRP deficiency was accomplished in *Crp^-/-^* mice by i.v. inoculation of 1 μg r-mCPR prior to infection as in (A) to test bacterial kinetics (left) and organ burden (right). n = 6. **D.** Capture of blood bacteria (green) by KCs (red) was visualized by IVM in the context of liver vasculatures (cyan) of *Crp^-/-^* and WT mice post i.v. infection. KC-associated bacteria are presented as immobilized bacteria per field of view (FOV). n = 2. Scale bar, 10 μm. **E.** CRP-mediated KC binding to *Sp*23F was measured *in vitro* by infecting primary mouse KCs (red) with *Sp*23F or *Sp*14 (green) in the presence of 10% mouse serum from *Crp*^-/-^or WT mice, and visualized by immunofluorescence microscopy 30 min later. n = 2. Scale bar, 10 μm **F.** Protective immunity of CRP against septic infection of *Sp*23F were monitored for 7 days post i.v. infection ed with 10^6^ CFU. n = 5-8. **G, H.** CRP-based immunity against other *S. pneumoniae* serotypes was assessed by CFU plating of the blood (G) and liver (H) at various time points post i.v. infection with serotype-representative strains as in (A) and (B), respectively. Presented are the data and capsule repeat units of serotypes 11A, 20B, 23A, 33F and 35B. The data for additional serotypes are available in Figure S2. n = 3-6.

Based on the role of KCs in hepatic capture of the low-virulence pneumococci,^45^ we tested the impact of CRP on hepatic capture of *Sp*23F using intravital microscopy (IVM) imaging. As expected, *Sp*23F pneumococci were rapidly tethered to KCs upon entering the vasculatures of the liver sinusoids of WT mice, but the circulating bacteria smoothly passed the KCs in *Crp*^-/-^ mice (**Figure 2D** and **Video S1**). The deficiency of *Crp*^-/-^ mice in pneumococcal capture is serotype-specific because *Sp*14 bacteria were similarly captured by KCs of WT and *Crp*^-/-^ mice (**Figure 2D** and **Videos S2**). The CRP-mediated binding to *Sp*23F was also tested *in vitro*. Primary mouse KCs abundantly bound to *Sp*23F in the presence of normal mouse serum; however, marginal bacterial binding was detected with serum from *Crp^-/-^* mice (**Figure 2E**). In contrast, CRP-deficiency showed no impact on KC binding to *Sp*14. These lines of evidence demonstrated that CRP is essential for specific recognition and capture of *Sp*23F by KCs.

To ascertain the contribution of CRP to host defense against *Sp*23F, we assessed the survival of *Crp^-/-^* mice after i.v. inoculation with 10^6^ CFU of *Sp*23F. As compared with the full survival of WT mice, all *Crp^-/-^* mice succumbed to the infection within 36 hours (hr) (**Figure 2F**). By comparison, *Crp*^-/-^ mice were fully competent in the defense against *Sp*14. The hyper-susceptibility of *Crp^-/-^* mice to *Sp*23F was also reflected by significantly higher bacteremia levels in *Crp^-/-^* mice in the first 24 hr post infection (**Figure S2F**). This result demonstrated that CRP confers a potent immunity against systemic infection of *Sp*23F.

### CRP broadly recognizes many capsule types of *S. pneumoniae*

We further tested CRP binding to CPSs of 24 additional pneumococcal serotypes in our collection, which showed the LV phenotype in our previous work.^45^ ELISA revealed broad binding interactions of mouse CRP with the capsular polysaccharides of many serotypes. r-mCRP bound to free CPSs of 16 additional serotypes: 11A, 15B, 15C, 16F, 17F, 20B, 21, 23A, 23B, 27, 33F, 35A, 35B, 35C, 37 and 41A, but not those of the other eight serotypes (9N, 9V, 10A, 14, 19A, 19F, 34 and 48) (**Figure S2D**). Except for 23B, the early clearance of all the other 15 serotypes was significantly impaired in *Crp^-/-^* mice although the degrees of the immune deficiency varied among these serotypes, in terms of the CT_50_ value (**Figures 2G** and **S2A**). For the sake of simplicity, we will refer the serotypes with significant clearance retardation in *Crp^-/-^* mice to as “CRP-sensitive” bacteria hereafter. Likewise, enumeration of organ bacteria at 5 min post i.v. inoculation revealed significantly decreased capture of the CRP-sensitive serotypes in the livers of *Crp^-/-^* mice as compared with WT mice (**Figure 2H**). Consistently, the vast majority of viable bacteria for the 15 CRP-sensitive serotypes were eliminated to residual levels in WT mice but still retained in *Crp^-/-^* mice at 30 min post infection (**Figures 2H** and **S2B**). This result showed that CRP-mediated hepatic capture of these serotypes leads to rapid bacterial killing. IVM examination of the liver sinusoids revealed potent capture of the CRP-sensitive serotypes by KCs of WT mice, but not *Crp^-/-^* mice (**Figure S2C** and **Videos S3-S7**). Notably, *Crp^-/-^* mice were equally able to capture and kill serotype-23B pneumococci as WT mice, indicating that this serotype is also recognized by another uncharacterized receptor(s).

Finally, the CRP-sensitive serotypes showed significantly enhanced virulence in *Crp^-/-^* mice, which is reflected by the reduced survival (**Figure S2E)** and persistent bacteremia (**Figure S2F**) of *Crp^-/-^* mice post i.v. inoculation with 10^6^ CFU of the bacteria. Together, these findings unequivocally demonstrated the great importance of CRP in broad but serotype-specific protection against septic infection of *S. pneumoniae* by enabling KCs to rapidly capture of these bacteria in the liver.

### CRP also recognizes the capsules of major Gram-negative pathogens

To understand if CRP recognizes any capsules of Gram-negative bacteria, we first tested the impact of CRP deficiency on the early clearance of *H. influenzae*, an important etiological agent of childhood pneumonia, septicemia and meningitis.^35^ Five among the six *H. influenza* capsule serotypes excepting serotype e were rapidly eliminated from the bloodstream post septic infection in WT mice (**Figure S3A**). The CT_50_ of serotype-a strain (*Hia*) was moderately elongated from 0.5 min in WT mice to 1.7 min in *Crp^-/-^*mice (**Figures 3A** and **S3A**), which was accompanied by substantial reduction in hepatic trapping of *Hia* at 5 min post infection (**Figure 3A**, right panel). *Hia* bacteria were significantly cleared in *Crp^-/-^* at 30 min, indicating the redundant clearance mechanisms against the bacterium in mouse. In contrast, infection of *Crp^-/-^* mice with *Hib* yielded a much more dramatic phenotype in early clearance. The bacteria were fully maintained in the circulation of *Crp^-/-^* mice in the first 30 min post infection (**Figure 3B**, left panel), and marginally sequestered in the livers at 5 and 30 min (**Figure 3B**, right panel). These results demonstrated CRP as an essential host factor in hepatic trapping and clearance at the early infection stage of blood-borne *Hib*, the most dominant serotype of *H. influenzae* in causing invasive infections in young children.^35^

**Figure 3.**
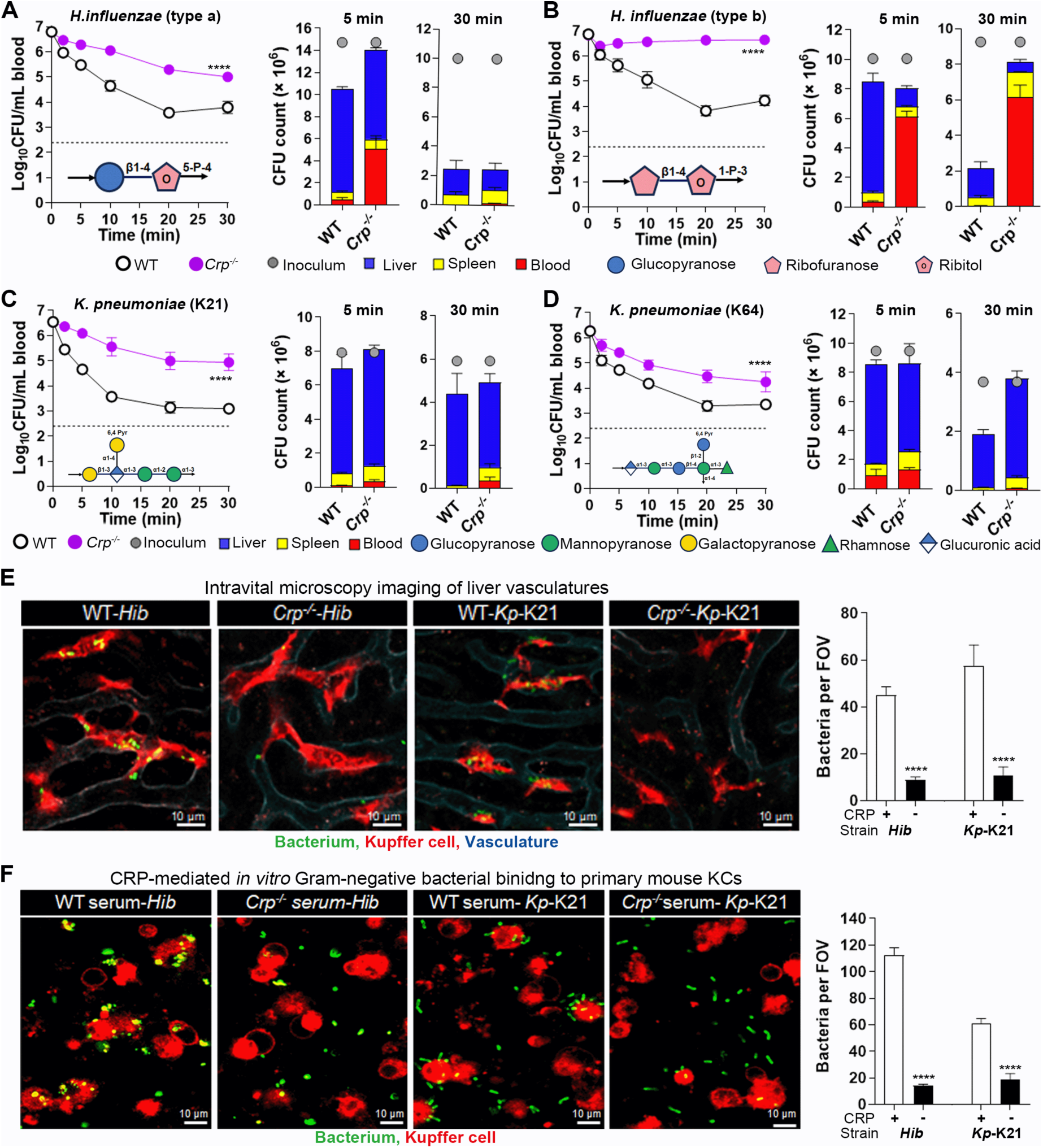
CRP-activated hepatic clearance of *H. influenzae* and *K. pneumoniae*. **A.** CRP-mediated clearance and hepatic trapping (5 min)/killing (30 min) of serotype-a *H. influenzae* were assessed by i.v. infection with 10^7^ CFU as in Figure 2A and 2B, respectively. The capsule repeat unit is diagrammatically presented below the dash line. n = 3. **B.** CRP-mediated clearance and hepatic trapping (5 min)/killing (30 min) of serotype-b *H. influenzae.* The data were obtained and presented as in (A). n = 3. **C.** CRP-mediated clearance and hepatic trapping (5 min)/killing (30 min) of serotype-K21 *K. pneumoniae.* The data were obtained and presented as in (A). n = 3. **D.** CRP-mediated clearance and hepatic trapping (5 min)/killing (30 min) of serotype-K64 *K. pneumoniae.* The data were obtained and presented as in (A). n = 3. **E.** IVM imaging of CRP-mediated capture of Gram-negative pathogens by KCs. The capture of serotype-b *H. influenzae* (left panel) and serotype-K21 *K. pneumoniae* (right panel) by KCs in the liver sinusoids of *Crp^-/-^* and WT mice were visualized and quantified by IVM as in Figure 2D. n = 2. **F.** CRP-mediated *in vitro* KC binding to *H. influenzae* and *K. pneumoniae* was assessed as in Figure 2E. n = 2.

We further tested the impact of CRP deficiency on the clearance of *K. pneumoniae,* an important Gram-negative pathogen of hospital-acquired pneumonia and sepsis with over 100 capsule types.^48^ Our recent study showed that the LV rather than the HV *K. pneumoniae* strains can be quickly cleared by liver KC, but no capsule receptor is known for any *K. pneumoniae* serotypes.^46^ The 11 serotypes tested in this trial were rapidly cleared from the bloodstream of WT mice. Besides, the clearance of serotypes K21 (**Figure 3C**) and K64 (**Figure 3D**) was significantly retarded in *Crp^-/-^* mice (**Figure S3B**). This finding revealed that CRP recognizes multiple capsule variants of *K. pneumoniae*.

Gram-negative bacteria in the liver sinusoids were further confirmed by IVM imaging. The KC-immobilized *Hib* bacteria were significantly lower in *Crp^-/-^* mice than in WT mice (**Figure 3E** and **Video S8**). In a similar fashion, *Crp^-/-^* mice showed a significantly reduced level of KC-associated *K. pneumoniae* K21 (**Figure 3E** and **Video S9**). Specific interactions of CRP with Gram-negative bacteria were confirmed with primary mouse KCs (**Figure 3F**). The bacteria of *H. influenzae* type b and *K. pneumoniae* K21 abundantly attached to KCs in the presence of normal serum, but the binding interactions were barely detectable with *Crp^-/-^* serum (**Figure 3F**). These imaging analyses showed that CRP promotes serotype-specific KC capture of *H. influenzae* and *K. pneumoniae*.

We further verified the direct binding of CRP to *H. influenzae* and *K. pneumoniae* capsules under the *in vitro* conditions. ELISA test revealed r-mCRP binding to the purified free CPSs of *H. influenzae* serotypes a and b (**Figure S3C**). In a similar manner, there was serotype-specific binding of r-mCRP to purified CPSs of serotypes-K21 and -K64 *K. pneumoniae* (**Figure S3D**). Taken together, these experiments uncovered that CRP could act as a plasma receptor for the capsules of multiple Gram-negative pathogens.

### The complement system relays the CRP-initiated anti-bacterial immune reaction

Since CRP is a plasma protein that is not physically associated with KCs, we reasoned that it must indirectly engages KCs for bacterial capture. Previous *in vitro* studies have shown that human CRP binding to PC promotes phagocytosis by binding to complement protein C1q and activating complement C3 via the classic complement pathway.^19,49^ Consistently, we detected C3 deposition on CPSs after incubation with WT serum, but not *Crp^-/-^* serum (**Figures S4A** and **S4B**). We further determined bacterial clearance in mice lacking C3 (*C3^-/-^*), a core protein in the complement system.^50^ The *C3^-/-^* mice showed significant retardation in clearing *Sp*23F (**Figure 4A**, left panel). In a similar pattern, *C3^-/-^* mice trapped relatively fewer bacteria in the liver than WT controls at 5 min post infection (**Figure 4A**, right panel). However, the phenotype of *C3^-/-^* mice was much milder than *Crp^-/-^* mice (**Figure 2**), indicating that C3-independent mechanism functionally bridges CRP and liver macrophage. To assess the relative contributions of C3-dependent and -independent mechanisms to the CRP-driven immunity, we tested bacterial clearance of *C3^-/-^* mice with an elevated infection dose of 5 × 10^7^ CFU, based on our preliminary trial (**Figure S1A**). *C3^-/-^* mice completely failed to clear *Sp*23F from the bloodstream (**Figure 4B**). Consistently, the absence of C3 led to the loss of hepatic capture of the bacteria. These data revealed that the C3-dependent and -independent mechanisms of CRP function are both capable of eliminating relatively milder form of blood infection, but CRP relies on the complement system to combat more severe infection.

**Figure 4.**
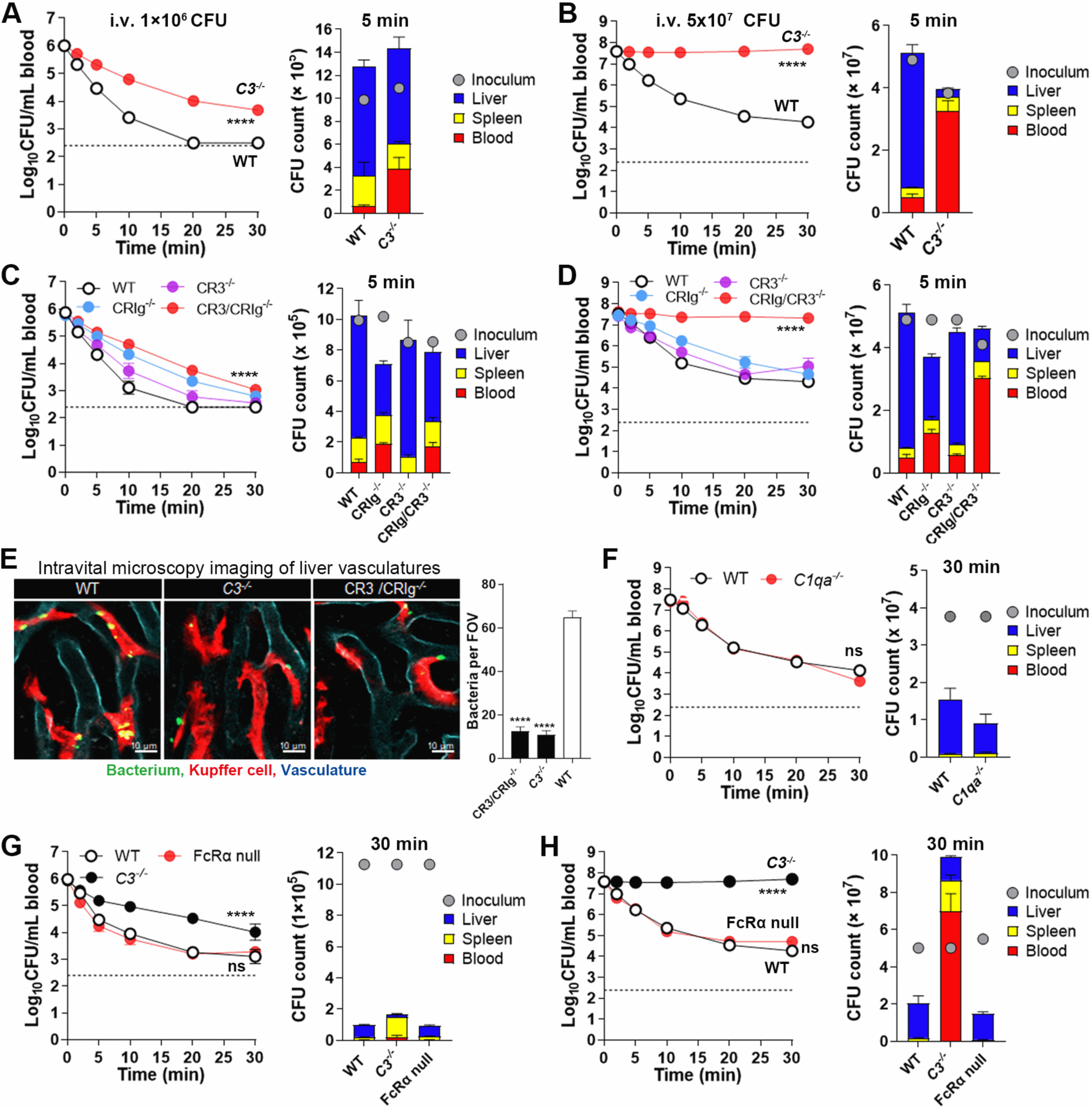
The role of C3 in the CRP-based immunity. **A**, **B**. The contribution of C3 to CRP-mediated immunity was evaluated by CFU plating of the blood (left) and liver (right) in *C3^-/-^* mice infected with a low (A) or high (B) dose of *Sp*23F as in Figure 2. n = 6. **C, D.** The role of C3 receptors CRs and CRIg in CRP-mediated immunity was assessed by CFU plating of the blood (left) and liver (right) in CR3^-/-^ and/or CRIg^-/-^ mice infected as in (A) and (B), respectively. n = 6. **E**. Visualization of CRP-mediated pathogen capture by KCs was performed in mice lacking C3 or C3 receptors as in Figure 2D. n = 2. **F**. The role of C1q in CRP-mediated immunity were tested in *C1qa^-/-^* mice as in (B). n = 3. **G, H.** The roles of Fcγreceptors in CRP-mediated immunity were tested in FcRα null mice with the low-(G) or high (H) inoculum of *Sp*23F as in (A) and (B), respectively. n = 3.

C3 is known to promote microbial phagocytosis by engaging complement receptors.^50^ Among the four major C3 receptors, mouse KCs express CR3 and CRIg,^45^ both of which have been shown to promote macrophage uptake of C3-opsonized microbes.^37,51^ In agreement with the infection dose-dependent phenotype of *C3^-/-^* mice, CR3- and CRIg-deficient mice showed marginally impaired clearance of blood-borne bacteria as compared with WT when infected with 10^6^ CFU bacteria, although CR3 and CRIg double KO mice showed a greater immune deficiency (**Figure 4C**). However, the double KO mice completely were completely unable to clear the at the high infection inoculum (**Figure 4D**), which was comparable to *C3^-/-^*mice. This observation indicated that KCs employ both CR3 and CRIg to capture of the C3-opsonized *Sp*23F pneumococci.

At the cellular level, IVM imaging also confirmed the extreme importance of C3 and the C3 receptors in CRP-based KC capture of *Sp*23F at the high infection dose. As compared with WT mice, the liver sinusoids C3*^-/-^*and CR3/CRIg*^-/-^* mice showed dramatic reduction in the KC-immobilized bacteria (**Figure 4E** and **Video S10**).

The previous studies have shown that human CRP-PC complex activates the complement classical pathway via binding to C1q, the first protein in the progressive line of the complement classical pathway ^19,52^. We thus tested the phenotype of *C1q^-/-^*mice post i.v. infection with 5 × 10^7^ CFU of *Sp*23F. In contrast to the dramatic loss of CRP-mediated bacterial clearance and hepatic capture of *Sp*23F in *C3^-/-^* mice (**Figure 4B**), *C1q^-/-^* mice showed a similar pattern of bacterial clearance and hepatic capture as WT mice (**Figure 4F**). This result demonstrated that CRP activates C3 in a C1q-independent manner.

To understand the C3-independent mode of the CRP immunity, we tested potential involvement of Fcγ receptors, based on the existing evidence that human CRP binds to FcγRI and FcγRIIA on phagocytes.^16–18^ However, the FcRα null mice lacking all of the four phagocytosis-associated FcγRs (FcγRI, FcγRIIB, FcγRIII and FcγRIV) showed comparable bacterial clearance and hepatic capture of *Sp*23F as WT mice when infected with 1 × 10^6^ CFU (**Figure 4G**). In a similar manner, the FcRα null mice did not show obvious defect in the early clearance and hepatic capture of *Sp*23F when the infection dose was raised to 5 × 10^7^ CFU (**Figure 4H**). These data showed that mouse Fcγ receptors are not involved in the C3-independent mechanism of CRP-activated bacterial clearance.

### CRP recognizes diverse polysaccharide structures via distinct binding modes

The CRP-binding capsules greatly differ in sugar composition, length and branch of repeating unit (**Figures 5A** and **S5**). Previous studies have revealed that CRP bind to phosphocholine (PC) and phosphoglycerol (PG) groups in a pocket around the Ca^2+^ binding sites.^15,53,54^ We determined the binding affinity between CRP and capsules with phosphocholine (PC) (CPS27), phosphoglycerol (PG) (CPS23F) or none of the either group (CPS33F) by isothermal titration calorimetry (ITC). The results showed that mCRP exhibited the highest affinity to CPS23F with a dissociation constant *K_D_* of 2.46 × 10^-7^ M (**Figures 5B** and **S6**). Besides, the binding stoichiometry of N = 4.91 suggested a molecular ration of 1:5 in the complex between mCRP and CPS23F ligands. Relatively lower binding affinity was obtained with CPS27 (*K_D_* = 1.1× 10^-5^ M) and CPS33F (*K_D_* = 3.58 × 10^-7^ M). These experiments showed that the binding pocket of CRP for PC, the best-known natural substrate of CRP, differs from those for the non-PC-containing capsular polysaccharides, suggesting that CRP recognizes bacterial capsules via distinct binding modes.

**Figure 5.**
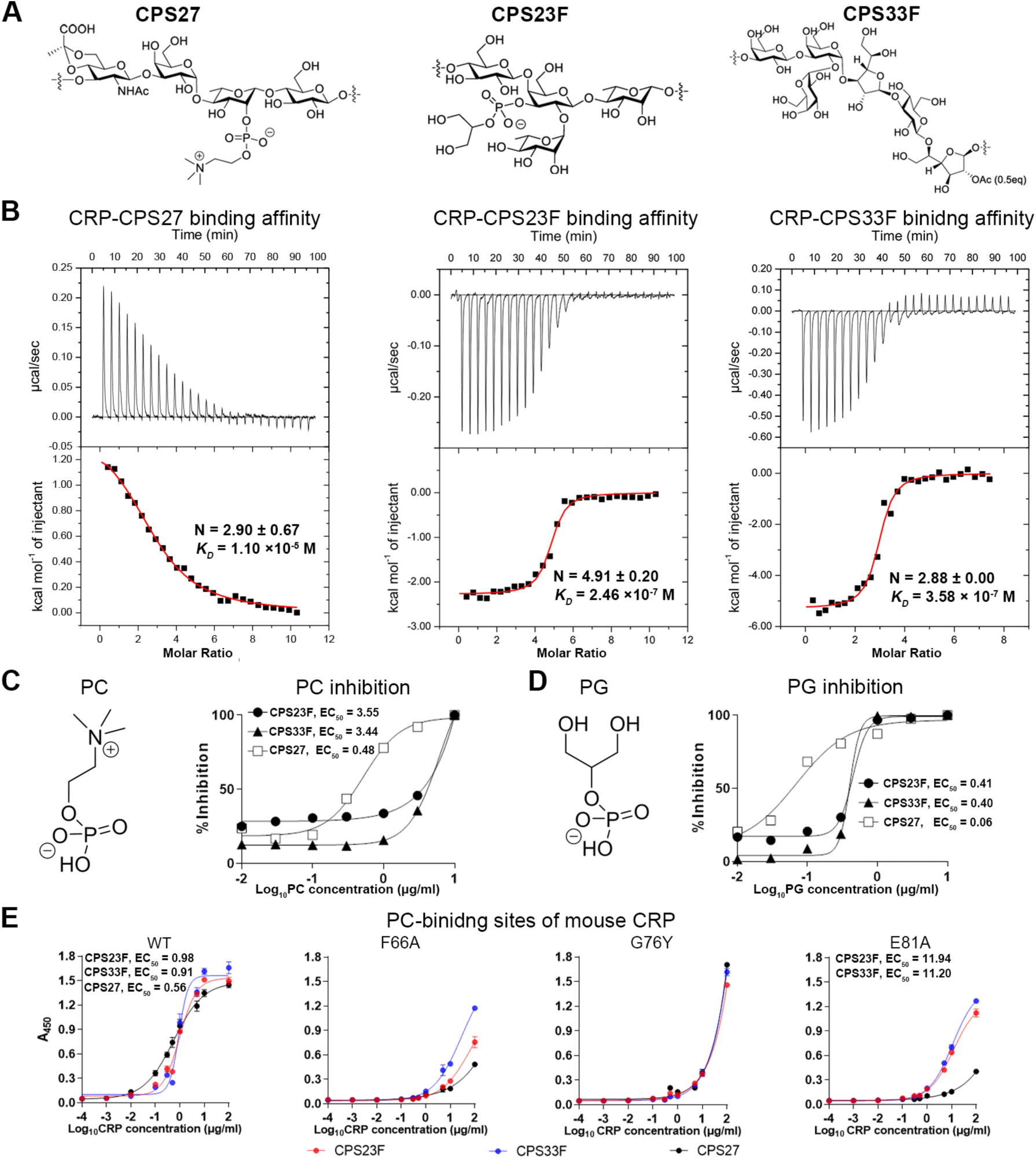
Molecular basis of capsule recognition by mouse CRP. **A.** The repeat unit structures of the PC-containing (CPS27), PG-containing (CPS23F) and PC/PG-free (CPS33F) capsules of *S. pneumoniae*. **B.** The binding affinity of CRP to CPS23F, CSP27 and CP33F was measure by ITC. The dissociate constant (*K_D_*), binding enthalpy (ΔH) and stoichiometry (N) are shown after base-line integration and concentration normalization. n = 2. **C.** Competitive inhibition of mCRP-capsule interactions by PC was tested by preincubating r-mCRP with various concentrations of PC (structure shown at the left panel) before being added to CPS-coated wells and detecting CPS-bound CRP as in Figure 1D (right panel). n = 3. **D.** Competitive inhibition of mouse CRP-capsule interactions by free PG as in (C). n = 3. **E.** Capsule binding of mCRP mutants was tested by ELISA using various concentrations of WT and mutant mCRP and plate wells coated with CPS23F, CPS33F, and CPS27 as in Figure 1D. n = 3.

To further define the capsule-binding modes of CRP, we tested the capacity of free PC and PG in competitively inhibiting the binding interaction between CRP and capsules with PC (CPS27), PG (CPS23F) or neither group (CPS33F). Competitive ELISA showed that free PC prevented mCRP from binding to CPS27 in a dose-dependent manner, suggesting that CRP recognizes PC and the PC-containing capsule in a similar mechanism (**Figure 5C**). In contrast, free PC had much weaker inhibition against mCRP binding to the PC-free CPS23F and CPS33F. The values of the media effective concentration of inhibition, EC_50_) for CPS27 was 7.4 and 7.2 folds lower than that for CSP23F and CPS33F, respectively. Surprisingly, free PG was more potent in inhibiting mCRP binding to CPS27 (EC_50_ = 0.06 μg/ml) than PC (EC_50_ = 0.48 μg/ml) (**Figure 5D**). Likewise, PG also showed more effectively blockage against PC-free CPS23F and CPS33F than PC. In particular, PG prevented mCRP from binding to the PG-containing (CPS23F) and PG-free (CPS33F) capsules in a similar manner. These data indicated that mouse CRP accommodates various structures of bacterial capsules with a highly dynamic platform.

To characterize the molecular basis of the differences in mCRP binding to capsular polysaccharides, we constructed mCRP point mutants in the 66^th^ phenylalanine, 76^th^ glycine and 81^th^ glutamic acid residues based on their location at the PC-binding surface of human CRP.^53,55^ The F66 and E81 are conserved in both mouse and human CRPs, whereas there is a cross-species polymorphism at the 76^th^ position between human (threonine) and mouse (glycine) (**Figure S7A**). The ELISA result showed the G76Y mutant showed a similar binding to CPS23F, CPS27 and CPS33F as the WT form (**Figure 5E**). The dispensability of this position in capsule binding is consistent with its cross-species polymorphism. The F66A and E81A mutation had significant impairment in mCRP binding to all the three capsules, but its impact on the binding activity of the PC-containing CPS27 was far more severe than on that of the PC-free capsules (CPS23F and CPS33F). Together, these data have revealed that mCRP recognizes the diverse structures of bacterial capsules by related but distinct binding modes.

### Human CRP mediates broad serotype-specific recognition of capsules

To understand the functional significance of human CRP in enabling hepatic clearance of encapsulated bacteria, we determined if hCRP shares the capsule-binding features of mCRP. While the ELISA result revealed significant binding of hCRP to all of the 17 mCRP-binding capsules of *S. pneumoniae* (**Figure 6A**), and thus demonstrated the common characteristic of human and mouse CRPs in anti-pneumococcal immunity. Surprisingly, no significant binding was detected between hCRP and the four mCRP-binding capsules of Gram-negative bacteria. This result revealed species-specific features of CRP orthologs in the pathogen recognition. We further verified the ability of human CRP in promoting hepatic bacterial capture using primary human KCs. As compared with the poor binding of serotype-8 pneumococci to human KCs, *Sp*23F bacteria showed a low but significant level of adhesion to human primary KCs in the presence of normal human serum (**Figure 6B**). This activity was further augmented by r-hCRP. These results strongly suggested that human CRP also acts as a plasma receptor for structurally diverse capsules to promoter hepatic capture of blood-borne bacteria.

**Figure 6.**
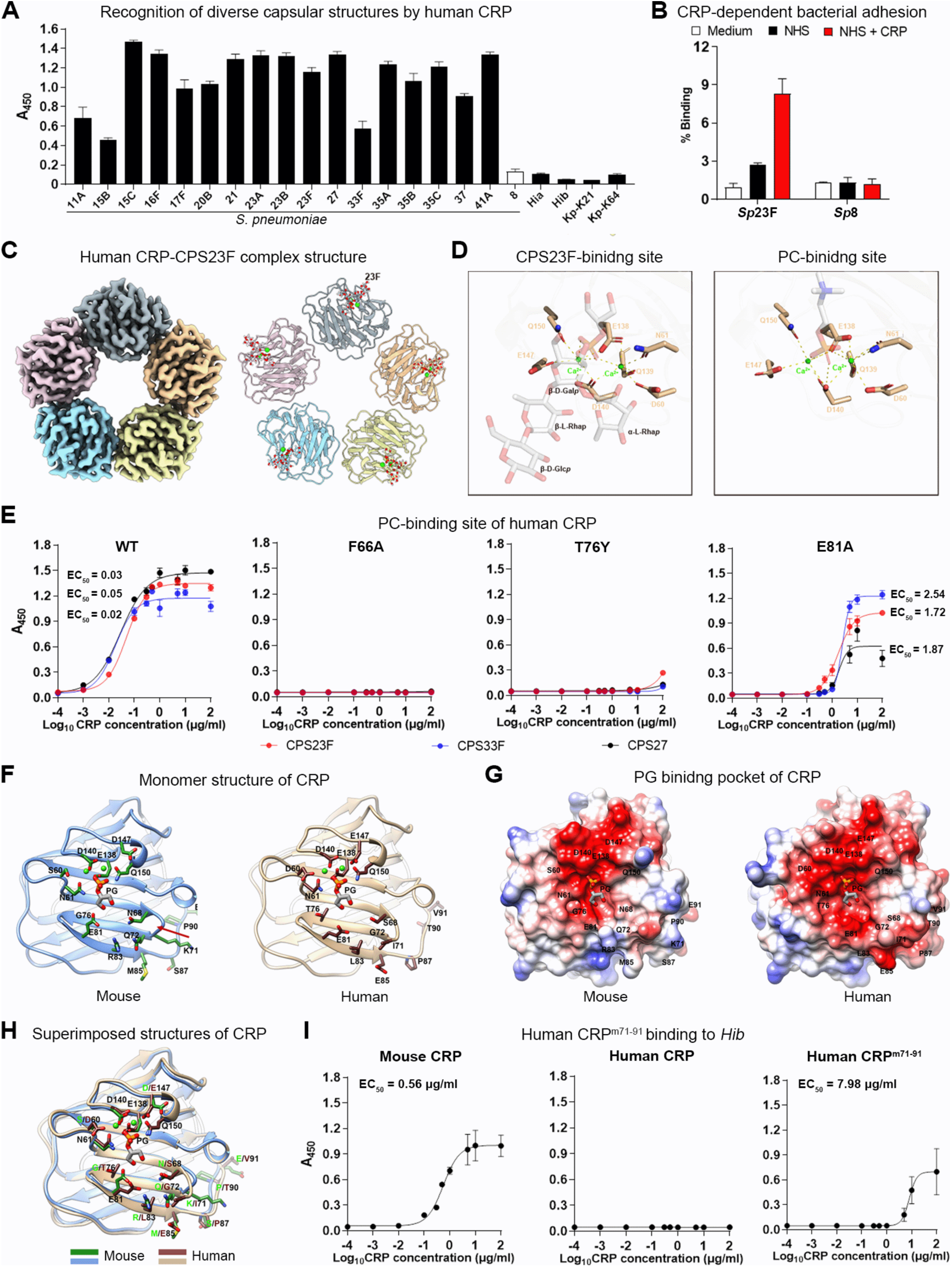
Immune recognition of structurally diverse capsules by human CRP. **A.** The *in vitro* hCRP binding to the capsules of *S. pneumoniae, H. influenzae* and *K. pneumoniae* was assessed by ELISA with 5 μg/ml r-hCRP as in Figure 1D. n = 3. **B.** CRP-mediated bacterial binding to primary human primary KCs was tested in the presence of 10% normal human serum (NHS) alone or in combination with r-hCRP (NHS + CRP, 10 μg/ml). No serum (medium) served as a negative control. KC-bound bacteria in each well are present as the ratio with free bacteria. n = 2. **C.** Cryo-EM structure of the hCRP-CPS23F complex. Five protomers are indicated with different colors (left panel); the two bound calciums shown as green spheres (right panel). **D.** Contact interface between human CRP and CPS23F. The left panel displays the molecular moieties of CPS23F (gray), key amino acids of hCRP (khaki) and two calciums (green) in the protein-polysaccharide complex. The previously published contact interface of the hCRP with PC (PDB: 1B09) is shown at the right panel as a reference. **E** Capsule binding of hCRP variants with mutations in amino acid residues structurally associated with CPS23F were tested with CPS23F, CPS27 and CPS33F as in A. n = 3. **F.** mCRP-PC complex structure predicted by Alphafold3 is presented along with the hCRP-PC structure. The bound PG and residues involved in calcium binding and around the PG binding pocket are shown in sticks with the O, P and N atoms colored red, orange and blue, respectively. The C atoms in the residues are colored dark green in mCRP and brown in hCRP. The red arrows indicate the loop 68-72, in which residue 71 flips in mCRP compared to that in hCRP. **G.** Surface rendered representation showing the binding pocket of PG. The surface is colored according to the surface electrostatic potential. The bound PG is shown in stick. **H.** The superimposed structures of mCRP and hCRP. The polymorphic residues in human and mouse CRPs are indicted by dark red and green letters, respectively. **I.** *Hib* capsule binding to the hybrid human-mouse CRP (hCRP^m71–91^) was determined and presented as in A, with normal hCRP and mCRP as controls. n = 3.

**Figure 7.**
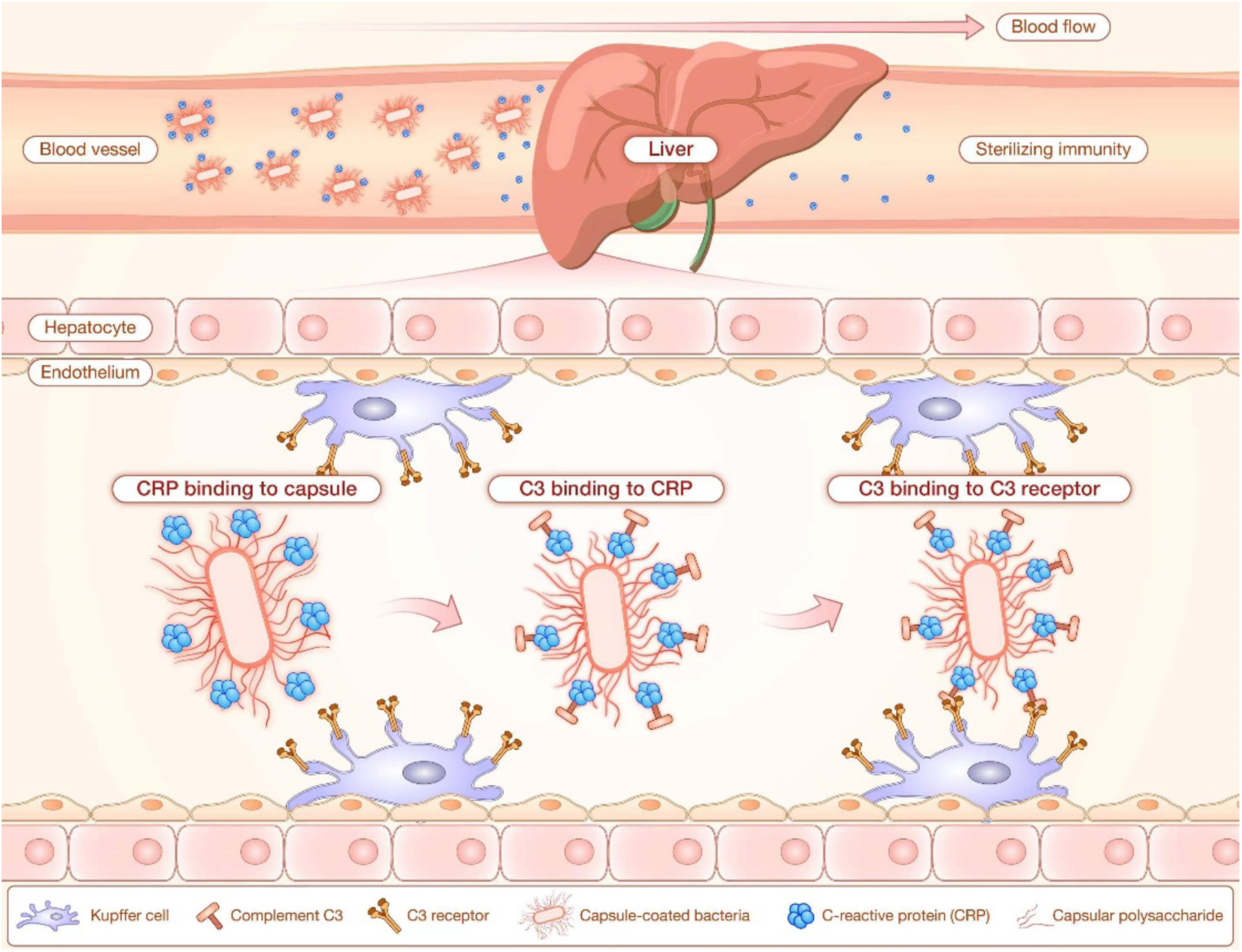
Diagrammatic model for the CRP-based blood sterilizing immunity in the liver. In the blood vessels, CRP binds to capsular polysaccharides of invading bacteria to achieve sterilizing immunity in the liver. The bacterium-bound CRP activates complement C3. The C3-opsonized bacteria are captured by the C3 receptors (CR3 and CRIg) on Kupffer cells that are embedded in the endothelium of the liver sinusoids.

We further determined the binding affinity of hCRP to five pneumococcal capsule types (**Figure S6**). hCRP showed the strongest binding to the PC-containing CPS27 with a dissociation constant of *K_D_*, 1.7 μM, which is comparable to the reported affinity between hCRP and the famous ligand PC (*K_D_*, 5 μM).^56^ However, except for CPS27, the binding affinities of hCRP to other four capsules are generally lower than those of mCRP. As an example, the CPS11A-binding affinity of mCRP (*K_D_*, 0.28 μM) is 100 folds of that of hCRP (*K_D_*, 27.90 μM), which may contribute to human tropism of *S. pneumoniae*. To define the molecular details of hCRP binding to capsular polysaccharides, we obtained the complex structure of hCRP and CSP23F by cryo-electron microscopy (EM) at a resolution of ∼2.78 Å (**Figure S8** and **Table S6**). The EM structure revealed the five identical non-covalently bound subunits of hCRP and Ca^2+^-dependent interactions with PG residues of CSP23F (**Figures 6C** and **6D**). In principle, the primary interactions between hCRP and CPS23F are similar to what is described for the hCRP-PC binding.^53,57^ The CPS binding site is formed by residues D60, N61, E138, Q139, D140, E147, and Q150 of hCRP, which are coordinated by two Ca^2+^ atoms (**Figure 6D**).

To define the precise roles of individual amino acids of hCRP in capsule recognition, we tested the impact of F66A, T76Y, and E81A mutation. While F66 and G76 were not essential for capsule recognition of mouse CRP (**Figure 5E**), the F66A and T76Y mutants completely lost the binding to CPS23F, CPS27 and CPS33F (**Figure 6E**), indicating that these residues are placed in more crucial positions in hCRP. hCRP E81A variant showed capsule type-dependent variations as observed with mCRP counterpart. The E81A mutants of both human and mouse CRPs showed much more severe impairment in binding to CPS27 than to CPS23F and CPS33F. In the context of undetectable binding of human CRP to the four Gram-negative capsules that are recognized by mCRP (**Figure 6A**), these mutagenesis results revealed functional commonality and differences between human and mouse CRPs in capsule recognition.

To determine the molecular basis of the selective capsule binding of human and mouse CRPs, we predicted the structure of mCRP by Alphafold3 and its complex structures with PC and PG (**Figures 6F-H**). Comparative analyses revealed major differences in the pocket that accommodates the capsular PC and PG groups. The pocket is overall based on residues 61-91, which form three antiparallel helices and two short loops (loops 69-72 and 76-81). The loop 68-72 has a flipped conformation in mCRP compared with that of hCRP. In addition, in the loop 76-81, a replacement of the threonine residue at position 76 with glycine in mCRP significantly alters the conformation of E81. G76 of mCRP would allow E81 to establish close contacts with the bound ligands. However, T76 of hCRP would push E81 away from the bound ligand. This result suggested that residues 71-91 mainly determine the functional difference between human and mouse CRPs. This notion was tested by constructing a human-mouse hybrid CRP - hCRP^m71–91^, in which amino acids71-91 of hCRP were replaced with those of mCRP. In contrast to the undetectable binding of hCRP to *Hib* capsule, hCRP^m71–91^ gained the capsule-binding capability of mouse CRP (**Figure 6I**). This result verified that the differences in amino acids 71-91 of human and mouse CRPs are primarily responsible for the mouse-specific binding to the *H. influenzae* capsule. The relatively lower affinity of hCRP^m71–91^ to CPS-Hib (EC_50_ = 7.98 μg/ml) than that of mouse CRP (EC_50_ = 0.56 μg/ml) suggested the contribution of additional amino acids of mCRP beyond the 71-91 region to capsule binding, which is supported by the reduced binding of hCRP^m71–91^ to CSP23F (**Figure S7D**). Together, these data have revealed that human and mouse CRPs share the common structural principles of capsule recognition, with certain species-specific differences that may explain host-specific tropism and virulence of encapsulated bacteria.

## DISCUSSION

CRP is a hall marker of severe bacterial infections in humans due to its dramatic rise in plasma during infection.^58^ Surprisingly, the precise function of CRP is yet a mystery nearly a century after its discovery.^1,7^ This study shows that CRP broadly recognizes a large number of capsules from three invasive bacteria tested thus far. These molecular interactions enable the liver macrophages to swiftly capture and kill blood-borne pathogens in the liver sinusoids, and confer remarkable levels of protection against septic infections. CRP-deficient mice were hyper-susceptible to septic infections of the CRP-recognizable capsule types. More importantly, CRP-mediated immunity appears to operate in humans because human and mouse CRPs share the major structural and functional features of capsule recognition. This study has thus discovered a potent and broad-spectrum anti-bacterial function for CRP.

Several proteins in human plasma have been shown to bind to pathogenic bacteria *in vitro*, including L-ficolin^59–63^ and the mannose-binding protein,^64^ but their precise roles in the immune clearance of blood-borne bacteria remain undefined. CRP is known for recognizing the cell wall PC moiety of *S. pneumoniae*,^1,65^ but this molecular interaction has not been linked to the *in vivo* bacterial clearance in the liver, the master machinery for clearing blood-borne microbes.^45,46^ Our data have unequivocally demonstrated that circulating CRP binds to a wide range of bacterial capsules and thereby enable KCs to capture encapsulated Gram-positive and -negative bacteria. To the best of our knowledge, CRP is a completely new plasma receptor that mediates broad but serotype-specific capture of encapsulated bacteria by KCs.

### CRP drives a potent innate immunity against encapsulated bacteria

Since the discovery of CRP,^1^ many studies have attempted to define the anti-infection function due to its massive induction upon septic bacterial infections.^58,66^ CRP function has been commonly studied in mice by passive administration or transgenic expression of human CRP, because it is believed that the low levels of endogenous CRP in mouse plasma (< 2 µg/ml) are insufficient to manifest the function of the copious human CRP during severe bacterial infections.^7^ Several studies show that human CRP is protective against *S. pneumoniae* by binding to the cell wall PC,^9–13^ but other lines of evidence have cast some doubt on the immunological significance of the CRP-PC interaction in host defense.^14,67^ CRP binding to pneumococcal cell wall does not stimulate effective phagocytic killing *in vitro,* as compared to the CRP binding to PC-containing serotype-27 capsule.^14^ This work has shown that the natural level of CRP in mice confers a robust immunity against septic infections of CRP-recognizable encapsulated bacteria. This CRP-based immunity resembles what is induced by pneumococcal capsular polysaccharide vaccine in mice.^68^ In this regard, CRP acts like an anti-capsule antibody.

The potency of the CRP-mediated immunity varies among the targetable capsule types. Among the 21 CRP-recognizable capsule types identified in this work, CRP is fully required for the early clearance of 13 types in the liver, and partially involved in the clearance of the other 7 types. This result indicates that CRP is the sole receptor for many capsule types. The partial clearance of certain CRP-binding capsule types in CRP-deficient mice indicates that these capsules are also recognized by additional receptor(s). Along this line, the full dispensability of CRP in the hepatic clearance of serotype-23B *S. pneumoniae* is likely due to the functional redundancy of the other receptor(s). Our discovery of CRP as a plasma receptor for many bacterial capsules also explains why these types of bacteria show the LV phenotype in mouse sepsis models in our previous studies.^45,46^

### CRP mediates a broad spectrum of anti-bacterial immunity

Our limited screening has identified 21 CRP-binding capsules from the 41 LV serotypes tested thus far. Except for serotype-23B *S. pneumoniae,* CRP-deficient mice showed significant impairment in the hepatic clearance of all the CRP-binding serotypes. Mice lacking CRP were much more susceptible to septic infections of all the CRP-recognizable capsule types. While the precise spectrum of the CRP-mediated immunity remains to be defined, it is reasonable to expect that CRP recognizes additional capsule types beyond those discovered in this work. Despite the broad coverage of encapsulated bacteria, the CRP-mediated immunity is highly specific only to the CRP-binding capsule structures/serotypes, CRP-deficient mice still retained normal hepatic immunity against the non-CRP-binding serotypes.

It is astonishingly that CRP binds to such a large list of structurally diverse capsules. Except for the PC moiety of serotype-27 capsule of *S. pneumoniae*, none of the remaining 20 CRP-binding capsules possess the PC group in their repeat units, the known substrate of CRP. Our cryo-EM structural analysis has provided a molecular basis for CRP recognition of the PG-containing capsules. CRP binds to the PG and PC groups in a similar manner via electric charge-based interaction between the positive-charged calciums in each CRP protomer and the negative-charged phosphate group in PG. While it is difficult to explain how CRP recognizes the vast majority of the PG-free capsules, the charge-based molecular attraction may operate for certain capsule types with the phosphate groups in their repeat units. The repeat units of CRP-binding serotypes a and b of *H. influenzae* each possess a phosphate group, which may interact with the CRP-based calciums. However, the phosphate group alone in capsular polysaccharide is not sufficient for CRP binding because certain capsules contain the chemical group does not bind to CRP (e.g. serotypes c and f of *H. influenzae*). Further studies are warranted to fully understand the structural basis of the broad CRP-capsule interactions. In this regard, diverse CRP interactions with bacterial capsules can be informatic models to decipher the generalizable molecular rules for protein-polysaccharide binding recognitions.

### CRP activates multiple hepatic immune pathways

The previous *in vitro* studies have shown that human CRP binding to PC and other related substrates activates the complement system.^19,22^ While this activity is attributed to CRP-mediated protection in mouse models of pneumococcal bacteremia,^24,25,69^ human CRP does not activate the mouse classic complement pathway.^70,71^ In this study, we have shown that the mouse complement system is required for CRP-mediated anti-bacterial immunity since C3-deficient mice were significantly impaired in clearing blood-borne *Sp*23F. Although human CRP-PC complex is known to activate the complement classical pathway via binding to C1q,^72^ C1q-deficient mice displayed normal level of CRP-mediated bacterial clearance. This finding indicates that CRP activates C3 via an C1q-independent mechanism.

The hepatic clearance of CRP-binding bacteria in C3-deficient mice at the low infection dose indicates that CRP also activates a C3-independent immune mechanism. Human CRP has been shown to bind to FcγRI- or FcγRIIA-transfected cells *in vitro*.^16,17^ However, none of the major Fcγ receptors are involved in the CRP-mediated immunity since mice lacking these Fcγ receptors kept the normal CRP-based immunity. It thus appears that an uncharacterized receptor(s) on KCs receives the CRP-opsonized bacteria. Alternatively, CRP may activate a C3-like adaptor molecule in the plasma, which in turn engages KCs

### This study demonstrates that CRP engages liver macrophages to remove blood-borne bacteria

Consistent with our previous report that LV types of encapsulated bacteria are trapped in the liver,^45,46^ IVM imaging showed that CRP-deficient mice lost the ability to capture the CRP-targetable capsule types by Kupffer cells in the liver sinusoids. The data obtained with mice lacking C3 revealed that the functional linkage between CRP and KCs can be fulfilled by activated by the C3-dependent pathway. The C3-opsonized bacteria are captured by CR3 and CRIg, the two dominant C3 receptors on KCs,^45^ because the KCs of CR3/CRIg-deficient mice lost the ability to capture circulating CRP-bound bacteria. This CRP-activated KC capture of blood-borne bacteria is reminiscent of hepatic bacterial capture that is driven by anti-capsule IgM antibodies. Our recent studies have shown that capsular polysaccharide vaccine-elicited IgM antibodies confers immune-protection by enabling KCs to capture encapsulated *S. pneumoniae* via the C3 and C3 receptor pathway.^68^ In this regard, CRP behaves like a natural antibody with the exceptionally broad spectrum of polysaccharide targets.

### Our data strongly suggest that CRP-mediated anti-bacterial immunity operates in humans

The current literature is overwhelming concentrated on human CRP. This is due to the low plasma level and irresponsiveness of mouse CRP to microbial infections and other stress conditions. Additionally, there is a high level of sequence homology between human and mouse CRPs. As a result, passive administration or transgenic expression of human CRP has been used to study CRP function in mice.^7,69^ In this study, we have demonstrated that the endogenous level of CRP in mice is sufficient to confer potent protection against septic infections by CRP-recognizable bacteria. Multiple lines of evidence strongly suggest that human CRP principally shares the anti-bacterial immunity. First, human CRP in normal human serum was identified as the specific receptor for the *Sp*23F capsule that is also recognized by mouse plasma CRP. Second, human CRP binds to all of the 17 pneumococcal capsule types that are recognized by mouse CRP. Lastly, the addition of recombinant hCRP in normal human serum significantly enhanced the adhesion of S*p*23F to primary human KCs. In this context, massive increase in plasma CRP in humans during severe bacterial infections appears to represent an important host defense response to combat high bacterial burden. This work has thus provided a functional explanation for dramatic increase of human plasma CRP in bacterial infections.

### The high potency and broad spectrum of CRP-mediated immunity may be exploited for therapeutic treatment of invasive bacterial infections

Encapsulated bacteria represent the major pathogens of invasive infections, and the leading microorganisms with extensive antimicrobial resistance.^26^ With the current shortage of effective antimicrobials, monoclonal antibodies have been considered as the therapeutic option.^73,74^ In this context, CRP may be an attractive choice for treating drug-resistant encapsulated bacteria based on its potency and broad spectrum. In this study, passive administration of 1 μg r-mCRP successfully restored hepatic capture of *Crp^-/-^* mice; moreover, additional r-hCRP dramatically improved the bacterial binding of *Sp*23F to primary human KCs, both supporting the therapeutic potential of CRP in treating septic infections.

### Limitations of the study

The molecular and cellular basis of anti-bacterial function of CRP is mostly characterized in mouse sepsis model. Because human and mouse CRPs showed a similar pattern of binding interactions with many capsule types, we expect that CRP serves an important immune function against invasive bacterial infections. This notion is supported by CRP-dependent binding of human primary KCs to mouse CRP-sensitive capsule types *in vitro*. However, it is known that human CRP binds to human C1q complement protein but not the mouse ortholog.^70^ The precise function of CRP in human innate immunity will need to be further characterized.

## STAR METHODS

### RESOUCE AVAILABILITY

#### Lead Contact

Additional information and requests including materials and resources are available by contacting the Lead Contact, Jing-Ren Zhang.

#### Materials Availability

All materials are available upon request, including bacterial and mouse strains generated in this work.

### EXPERIMENTAL MODEL AND SUBJECT DETAILS

#### Human subjects

Human liver sections were obtained from freshly disposed tissues of liver surgery or transplantation patients with the approval by the Tsinghua University Science and Technology Ethics Committee (Medicine) (THU01-20240036).

#### Mice

All infection experiments were conducted in C57BL/6 (6-8 weeks old) according to the animal protocols approved by the Institutional Animal Care and Use Committee in Tsinghua University. All of gene-deficient mice were maintained in the C57BL/6 background. *Crp*^-/-^ mice and *C1qa*^-/-^ mice were acquired from Gempharmatech (Nanjing, China). *C3*^-/-^ mice were purchased from the Jackson Laboratory (Bar Harbor, Maine, USA). CRIg^-/-^ mice^37^ were obtained from Genentech (CA, USA). Complement receptor 3 (CR3)-deficient mice were generated with CRISPR/Cas9 system as described.^68^ CR3/CRIg^-/-^ mice were generated by crossing CRIg^-/-^ mice with CR3^-/-^ mice. FcRα null mice were generously provided by Jeffery Ravetch.^75^

#### Bacteria

All of the *S. pneumoniae*, *H. influenzae*, and *K. pneumoniae* strains used in this study are described in Table S1.

#### Primary cells

Mouse and human Kupffer cells were isolated from fresh liver sections as described.^45^

#### Cell lines

HEK293F cells were cultured in SMM 293-TII Expression Medium and grown at 37 °C with 5% CO_2_.

### METHOD DETAILS

#### Bacterial cultivation

Pneumococci were cultured in Todd-Hewitt broth with 0.5% yeast extract (THY) or tryptic soy agar (TSA) plates with 3% defibrinated sheep blood at 37 °C and 5% CO_2_ as described.^76^ *H. influenzae* strains were propagated under the same conditions in brain-heart infusion (BHI) broth or on BHI agar supplemented with 10 µg/ml hemin (Sigma) and 10 µg/ml NAD (Sigma) as described.^77^ *K. pneumoniae* strains were grown in Luria-Bertani (LB) broth or on LB agar plates as described.^46^

#### Mouse infection

Infection experiments were conducted in C57BL/6 (Vital River, Beijing, China) according to the animal protocols approved by the Institutional Animal Care and Use Committee in Tsinghua University. All mice were kept under specific pathogen free (SPF) conditions with free access to food and water. Septic infections were carried out and analyzed as described.^45^ Briefly, bacteria in 100 μl of Ringer’s solution were i.v. injected into tail vein. Bacteria in the blood were assessed by retroorbital bleeding and CFU counting on blood agar plates. Bacterial 50% clearance time (CT_50_) was calculated by nonlinear regression analysis of bacteremia kinetics using the formula T = ln{(1-50/Plateau)/(- K)}. Organ bacteria were similarly quantified with homogenized tissues. Total viable bacteria in each mouse were estimated as the sum of CFU values from blood and organ samples. Competitive inhibition of bacterial clearance was performed by i.v. administration of CPS 2-5 min prior to bacterial inoculation. For passive protection, bacteria were incubated with mouse serum or r-mCRP in 100 μl phosphate-buffered saline (PBS) supplemented with 2 mM CaCl_2_ at room temperature for 5 min before being bacterial i.v. inoculation. Survival rate was determined by monitoring infected mice for 7 days post infection. LD_50_, lethal dose 50%, was calculated by infection dose and percent of survival using LD_50_ calculator (Quest GraphTM, AAT Bioquest, Inc.).

#### CPS purification

CPSs were purified from broth cultures of *S. pneumoniae*, *H. influenzae* and *K. pneumoniae* strains and quantified as described previously.^45,46^

#### Isolation of liver non-parenchymal cells (NPCs)

Human and mouse liver non-parenchymal cells (NPC) were isolated by a collagenase-DNase digestion procedure as described previously.^45^ The concentrations of membrane proteins were quantified with the BCA Assay kit (Solarbio).

#### Screening for CPS23F-binding proteins

Mouse and human capsule-binding proteins were screened by an affinity pulldown approach as described.^78^ In brief, membrane proteins of mouse liver NPCs were enriched by the Mem-PER Plus Membrane Protein Extraction Kit (Thermo Scientific) according to the manufacture’s instructions. Capsule binding-proteins were enriched by co-incubating CPS23-coated latex beads with mouse liver NPC membrane proteins in the presence of normal mouse serum 10% (v/v) at room temperature for 1 hr. Beads with CPS8 from the HV serotype 8 was similarly processed as a negative control. Proteins bound to the CPS-coated beads were identified by mass spectrometry. Protein abundance was compared between CPS23F- and CPS8-coated beads to identify CPS23-enriched proteins. Proteins with at least 2-fold enrichment by CPS23F were considered as CPS23-binding candidates. CPS23-binding proteins in human serum were identified in a similar manner by co-incubating CPS23- and CP8-coated beads with 10% normal human serum.

#### Production of recombinant CRPs

Mouse and human CRPs were expressed as Strep-tagged recombinant proteins in human HEK293F suspension culture cells as described.^79^ Briefly, the complementary DNAs (cDNAs) were generated with the total RNAs of mouse liver to amplify the coding sequence of mCRP with primers Pr19504 and Pr19505. The amplicon was cloned in the SalI/BamHI site of pCMV-chikv-strepII vector with a Srep-tag (WSHPQFEK)^79^ to generate pCMV-mCRP (pTH17157). The coding sequence of hCRP was chemically synthesized according to accession NM_000567.3, and cloned in the PstI/BamHI site of pCMV-chikv-strepII vector to generate pCMV-hCRP (pTH17158). The insert sequences were confirmed by DNA sequencing. Recombinant human CRP was expressed by transient transfection of HEK293F cells, isolated from culture supernatants by affinity chromatography with Strep-Tactin sepharose resin (IBA) and eluted with 2.5 mM desthiobiotin (IBA) in TBS-Ca^2+^ buffer (20 mM Tris, 150 mM NaCl, 5 mM CaCl_2_, pH 8.0). While for mCRP, except for the buffer used was high salt TBS-Ca^2+^ buffer (20 mM Tris, 500 mM NaCl, 5 mM CaCl_2_, pH 8.0), other procedures were exactly the same. The fractions containing target proteins were pooled and proceeded for desthiobiotin removal. Protein concentration was quantified using the BCA Assay Kit. pCMV-mCRP was used as template for the construction of F66A (TH17199), E81A (TH17198), and G76Y (TH17202) mutant mouse CRP. The upstream and downstream fragments of mutant mouse CRP were amplified and were fused to generate mutant mouse CRP cDNA, which was then subcloned into SalI/BamHI sites of pCMV-chikv-strepII vector. The F66A (TH17201), E81A (TH17200), and T76Y (TH17202) mutant human CRP was similarly constructed. The recombinant mutant fragment was subcloned into PstI/BamHI sites of pCMV-chikv-strepII vector. Mutant plasmids of pCMV-hCRP^m71–91^ (TH17388) were constructed by site-directed mutagenesis as described.^80^ Briefly, pCMV-hCRP served as the template, and the PCR products were circularized using ClonExpress Ultra One Step Cloning Kit (Vazyme). These mutant CRP were expressed and purified as describe above. The strains and primers used in this study were respectively listed in Table S3 and Table S4.

#### Enzyme-linked immunosorbent assay (ELISA)

CRP binding to CPS was assessed by ELISA principally as described.^68^ Briefly, 96-well plates were coated with purified CPSs and blocked with 5% non-fat milk before sequentially incubating with Strep-tagged recombinant CRP, Strep-Tag mouse monoclonal antibody (0.5 μg/ml) and HRP-conjugated goat anti-mouse IgG (H+L). All the reagents were diluted in TBST-Ca^2+^ (TBS-Ca^2+^ with 0.05% Tween-20). CRP-CPS binding was quantified by adding 100 μl TMB chromogenic substrate (TIANGEN) and measuring optical density (OD) at 450 nm. Competitive inhibition of the CRP-CPS binding was accomplished by adding CRP and free CPS, PC or PG to CPS-coated wells simultaneously. Titration curves were generated using sigmoid dose response of nonlinear fit from GraphPad, and effective concentration of inhibition (EC_50_) were determined.

To determine the complement activation levels by CPSs, 100 μl of 10% mouse serum in PBS supplemented with 2 mM CaCl_2_ and 2 mM MgCl_2_ was added to CPS-coated plates. The plates were then incubated at 37 ℃ for 5, 10, 20, and 30 min, respectively. And 5,000-fold diluted HRP-conjugated Goat anti-mouse C3 IgG antibody was used to detect the amount of C3 fragments deposited on CPSs.

#### *In vitro* CRP binding to encapsulated bacteria

CRP binding to live bacteria was detected by immunofluorescent microscopy essentially as described.^76^ Briefly, bacteria (10^5^CFU) were sequentially treated in 100 μl each of 5% non-fat milk (w/v), 10 μg/ml r-mCRP, rabbit anti-mCRP antibody (1:200 dilution), and AF647-conjugated goat anti-rabbit IgG (1:200 dilution). Each incubation step was followed by centrifugation and resuspension in 100 μl PBS. The bacteria were resuspended in 20 μl PBS and visualized under a Leica TCS SP8 confocal microscope.

#### Intravital microscopy (IVM)

IVM imaging of mouse liver sinusoids was conducted as described.^45^ LSECs and KCs were labeled by i.v. injection of AF594 anti-CD31 and AF647 anti-F4/80 antibodies, respectively before i.v. administration of FITC-labeled bacteria. Images of the liver vasculatures were acquired with Leica TCS-SP8 confocal microscope using 10×/0.45 NA and 20×/0.80 NA HC PL APO objectives 10-15 min post infection. Photomultiplier tubes (PMTs) and hybrid photo detectors (HyD) were used to detect fluorescence signals (600 × 600 pixels for time-lapse series and 1024 × 1024 pixels for photographs). At least 5-10 ten random fields of view (FOV) were captured to calculate bacterial number per FOV. Leica Biosystems software was used for image and movie processing (LAS X Life Science).

#### Bacterial binding to primary human and mouse Kupffer cells

Bacterial binding to primary KCs *in vitro* was evaluated as described.^45^ In brief, KCs from human and mouse liver NPCs were isolated as abovementioned, and immobilized to the bottom of 48-well plates for 30 min at 37°C before being infected with 1 × 10^5^ CFU of bacteria (MOI = 1) in the presence or absence of 10% serum (v/v) for 30 min at 37 °C with 5% CO_2_. The CFUs in culture supernatants and KC lysates were enumerated as free and KC-bound bacteria. The *in vitro* visualization of bacterial binding to KCs was carried out by immunofluorescence microscopy as recently documented.^81^ Mouse liver NPCs were seeded in 4-chamber plates with 35mm glass slips (Cellvis). After non-adherent cells were removed, KCs were stained with AF647 anti-F4/80 and infected with FITC-labeled bacteria (MOI = 10) in the presence of 20% mouse serum for 30 min at 37 °C with 5% CO_2_. KCs and bacteria were visualized using the Leica TCS SP8 confocal microscope. Photographs with more than 100 bacteria were chosen for quantification.

#### Isothermal titration calorimetry (ITC)

The binding characteristics of CRP to CPSs were measured at 25 ℃ using a VP-ITC Microcalorimeter (Malvern Instruments Ltd) as described.^76^ In principle, the titrant (CPS) and titrated molecules (CRP) were prepared in the TBS-Ca^2+^ buffer. Each CPS was diluted to a 50-fold final concentration of recombinant CRP, except hCRP-27 (1:35) and hCRP-33F (1:40). CPS solutions were injected in the CRP-containing calorimeter cell with a titration volume of 10 μl and a spacing time of 210s. Calorimetric data were recorded in real-time, and analyzed to acquire the stoichiometry (N, the molar ratio of binding), the association equilibrium constant (*Ka*), binding entropy (ΔS), and binding enthalpy (ΔH) with Origin software (Version 7, MicroCal). The binding affinity is presented as dissociation equilibrium constant (*Kd*) (1/*Ka*).

#### Western blotting

To detect complement activation levels (deposition) on *S. pneumoniae*, 10^7^ CFU of pneumococci were incubated with 10% mouse serum in 100 μl PBS supplemented with 2 mM CaCl_2_ and 2 Mm MgCl_2_ at 37 ℃ for 20 min. The reaction was stopped by adding 1 ml ice-cold PBS. Bacteria were collected by centrifugation at 10,000 g, 4 ℃ for 2 min, and washed three times with ice-cold PBS. Bacterial pellets were suspended in 10 μl 1 ×SDS-PAGE loading buffer, and boiled for 10 min. Supernatants were collected by centrifugation at 10,000 g, 4 ℃ for 2 min, and were then subjected for SDS-PAGE.

A volume of 10 μl sample was loaded to the well, and proteins were separated by 12.5% SDS-PAGE gel. Then proteins were wet-transferred to polyvinylidene difluoride (PVDF) membrane. C3 fragments were detected by 5,000-fold diluted HRP-conjugated Goat anti-mouse C3 IgG antibody.

#### Cryo-EM sample preparation and data collection

Human CRP was concentrated to 6 mg/ml, and incubated with CPS23F to a final concentration of 4.5 mM on ice for ∼ 1h. Tween-20 was added to the solution to a final concentration of 0.02% (w/v). Droplets (3.5 μl) of the complex were applied to glow-discharged carbon-coated copper grids (Quantifoils R1.2/1.3 300 mesh), which were blotted for 2 seconds before being plunge frozen in liquid ethane using Vitrobot Mark IV (Thermo Fisher Scientific). Gain normalized movies were collected on a Titan Krios microscope (ThermoFisher) operated at 300 keV and equipped with a Gatan K3 detector. Movies were acquired automatically using EPU software using a total dose of 50 e^-^/Å^2^ with 32 frames, at 105,000 × magnification with a calibrated pixel size of 0.85 Å and a defocus range of -1.0 to -2.0 µm.

The movie frames were motion-corrected using MotionCor2 (v1.2.4) ^82^ and the contrast transfer function (CTF) values of each micrograph were calculated using patch CTF estimation implemented in cryoSPARC (v.4.1.0).^83^ All imaging processes were performed using cryoSPARC unless mentioned elsewhere. The map was finally sharpened by DeepEMhancer.^84^ We first picked ∼600,000 particles from 1,000 micrographs, and these particles were subjected to perform 2D-classification and generated a particles-dataset for Topaz training.^85^ To strengthen the Topaz procedure’s effect in particle picking, we retrained Topaz model using newly 2D-classification-filtering particles or heterogeneous refinement-filtering particles. Then, we applied well-trained Topaz model to pick up particles from the entire dataset, and 854,515 particles were extracted and subjected to 2D-classification. After 2D-classification, a clean dataset containing 506,697 particles was selected to perform initial reconstruction and heterogeneous refinement. 207,101 particles were separated out, and a density map at 2.78 Å resolution estimated by the gold-standard FSC 0.143 criterion was obtained after CTF refinement and non-uniform refinement. The image-processing workflow is summarized in Table S6.

To build the pentameric model for hCRP and CPS23F complex, a monomeric subunit from a crystal structure of human CRP (PDB code 1B09) ^53^ was placed into the cryoEM map as a rigid body using UCSF ChimeraX.^86^ The model was then manually adjusted using Coot.^87^ The model of this monomer was duplicated into the remaining 4 sites as rigid bodies using ChimeraX. The resulting pentameric model was automatically refined using Phenix real-space refine with Ramachandran restraints, secondary structural restraints, and geometry restraints.^88^ The output from Phenix was manually inspected and refined for several rounds aided by stereochemical quality assessment using MolProbity.^89^ Besides, the structure of mouse CRP was predicted by AlphaFold (https://alphafold.ebi.ac.uk/).^90^ Structural figures were prepared using PyMOL (www.pymol.org/) and UCSF ChimeraX.

#### Sequence analysis

DNA and amino acid sequences were analyzed using the DNASTAR Lasergene version 15.0 for Macintosh (Madison, WI).

### QUANTIFICATION AND STATISTICAL ANALYSIS

Statistical analyses were performed using GraphPad Prism software version 8.0.0, and all data were expressed as Mean ± SEM unless otherwise stated. Two-sided unpaired student’s t-test was used to determine the statistical significance between two groups, and two-way ANOVA multiple comparisons test was used to analyze data between multiple groups. Survival curves were analyzed by log-rank (Mantel-Cox) test. *P* value < 0.05 was considered as significant (not significant, ns; \**P* < 0.05; \*\**P* < 0.01; \*\*\**P* < 0.001; \*\*\*\**P* < 0.0001).

## ACKNOWLEDGEMENTS

We thank the Tsinghua research platforms for assistance in animal experimentation (Laboratory Animal Research Center), flow cytometry and IVM imaging (Center for Cell Biology) and protein mass spectrometry (Center for Proteomics). We would be grateful to Yuanyuan Chen, Zhenwei Yang and Bingxue Zhou (Institute of Biophysics, Chinese Academy of Sciences) for technical help with ITC experiments. This work was supported by grants from National Natural Science Foundation of China to J.-R.Z. (31820103001 and 31530082), Tsinghua University Spring Breeze Fund to J.-R.Z. (20201080767) and the Tsinghua-Peking Joint Center for Life Sciences Postdoctoral Foundation and the China Postdoctoral Science Foundation to H.A. (2016M590085).

## DECLARRATION OF INTEREST

The authors declare no competing interests.

## AUTHOR CONTRIBUTIONS

Conceptualization, J.H., D.C., H.A. and J.-R.Z.; experimentation, D.C, J.H., M.Z., Y.X., H.Y., Y.M., J.W. and X.H.; methodology, Z.S. and Y.X.; data analysis, J.H., D.C., Y.H., J.Q, G.F.G and J.-R.Z.; composition, J.H., D.C., Y.H., J.Q, G.F.G and J.-R.Z.; funding, H.A. and J.-R.Z.

---

**Figure S1.**
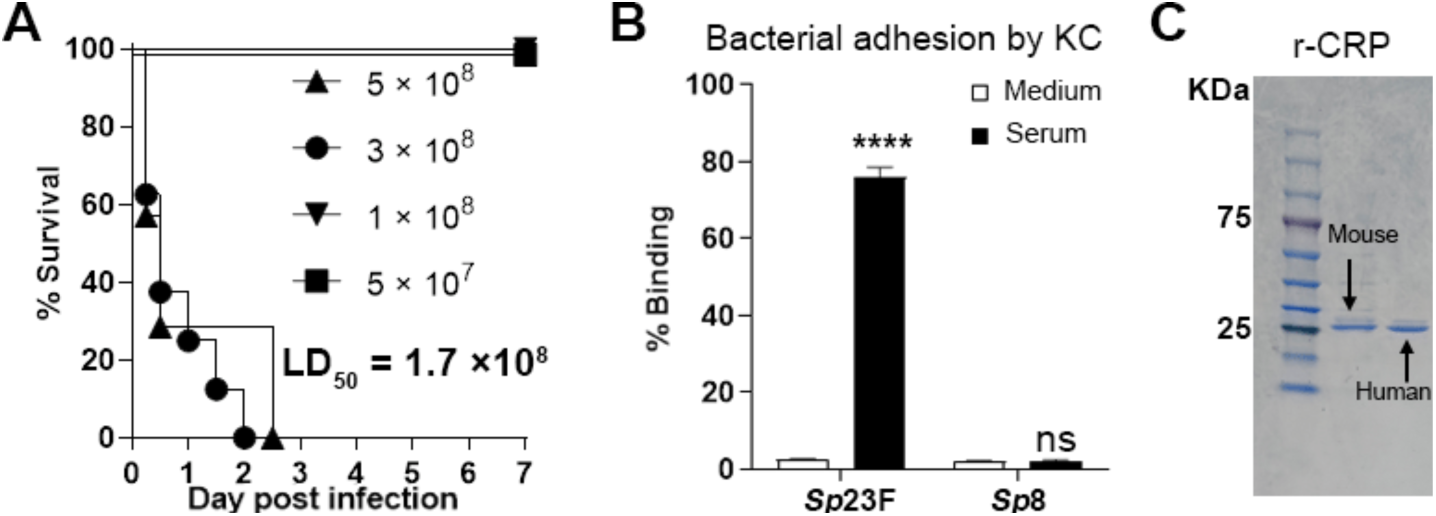
The virulence level of serotype-23F *S. pneumoniae*. **A.** Dose limit of WT mice against *Sp*23F. WT mice were i.v. infected with different doses of *Sp*23F to determine survival. n = 6-8. **B.** The requirement of serum for *in vitro* KC binding to *Sp*23F. Primary mouse KCs were infected with *Sp*23F at an MOI of 1:1 (KC vs. bacterium) in the presence or absence (medium) of mouse serum. The free and cell-associated bacteria were quantified by CFU plating 30 min post infection, and presented as their ratio values. n = 3. **C.** Construction of r-mCRP and r-hCRP. Purified r-mCRP and r-hCRP were detected by SDS-PAGE electrophoresis. n = 2.

**Figure S2.**
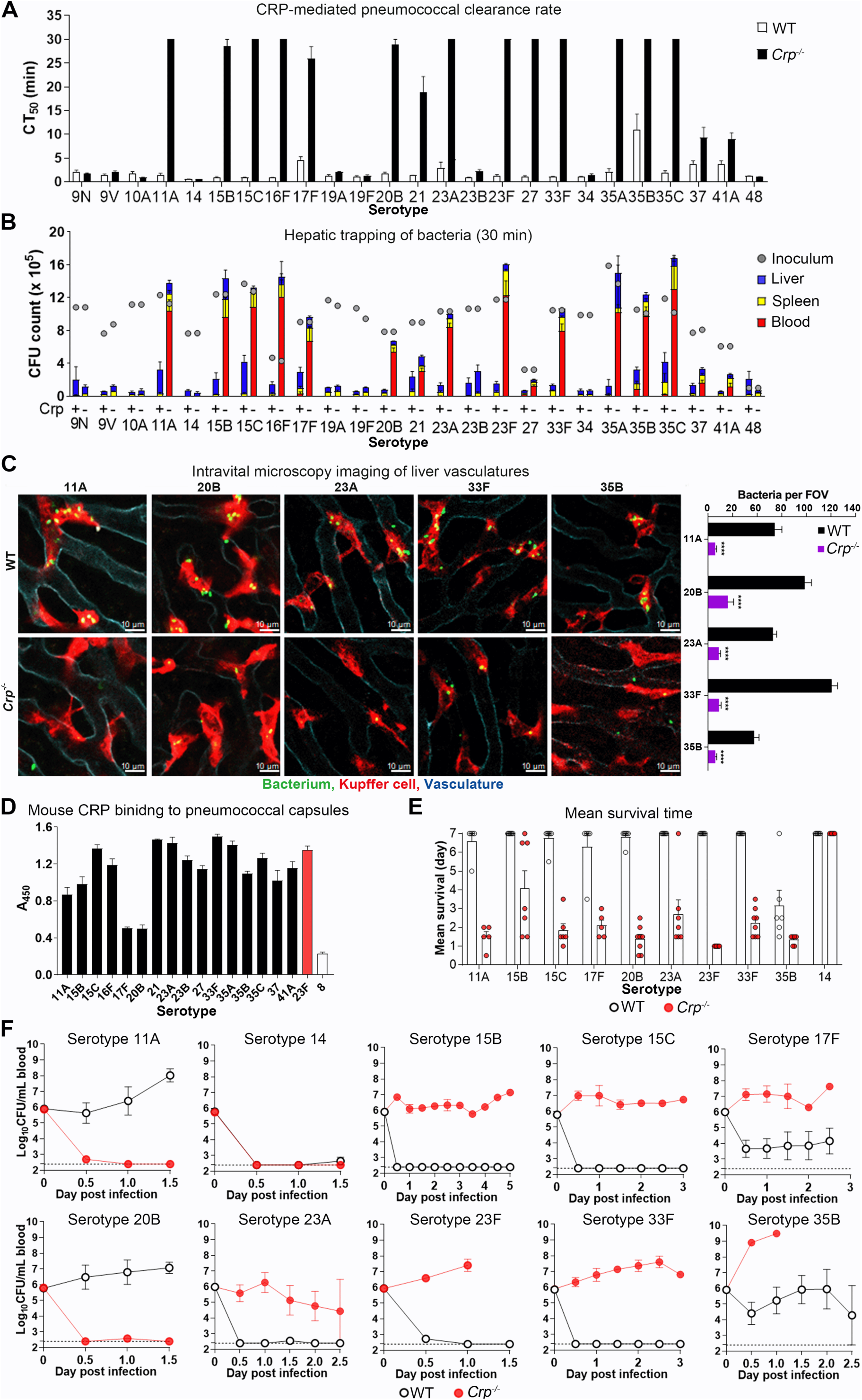
The requirement of CRP for broad serotype-specific shuffling of many pneumococcal serotypes from the blood circulation to the liver. **A.** The clearance rates of 25 low-virulence pneumococcal serotypes from the bloodstream in WT and *Crp^-/-^* mice infected i.v. with 10^6^ CFU of each serotype. CT_50_ showed that *Crp^-/-^* mice significant delayed in clearing 16 of the 25 serotypes. n = 3-5. **B.** Serotype-specific distribution of viable bacteria in the blood, liver and spleen of WT and *Crp^-/-^* mice at 30 min post i.v. infection. The mice used to determine bacteremia kinetics in (A) were sacrificed at 30 min to quantify viable bacteria in the blood, liver and spleen by CFU plating. The inoculum of each group is indicated with a filled circle. n =3-5. **C.** IVM imaging of CRP-mediated KC capture of 5 representative CRP-sensitive serotypes of *S. pneumoniae* in WT and *Crp^-/-^* mice as in Figure 2D. n = 2. **D.** mCRP binding to multiple pneumococcal capsules. The 96-well plates were individually pre-coated with purified pneumococcal capsules, and then incubated with 5 μg/ml r-mCRP. The CPS-bound CRP was then detected and represented by the absorbance at A_450 nm_ as in Figure 1D. n = 3. **E.** Broad protection of CRP against septic infection of CRP-sensitive serotypes. WT and *Crp^-/-^* mice were i.v. infected with 10^6^ CFU of *Sp*23F, *Sp*14, *Sp*11A, *Sp*15B, *Sp*15C, *Sp*17F, *Sp*20B, *Sp*23A, *Sp*33F, and *Sp*35B. The survival of WT and *Crp^-/-^* mice were assessed as in Figure 2F. n = 5-8. **F.** The bacteremia kinetics of WT and *Crp^-/-^* mice infected i.v. with 10^6^ CFU of 10 different serotypes as in Figure S2E. n = 5-8.

**Figure S3.**
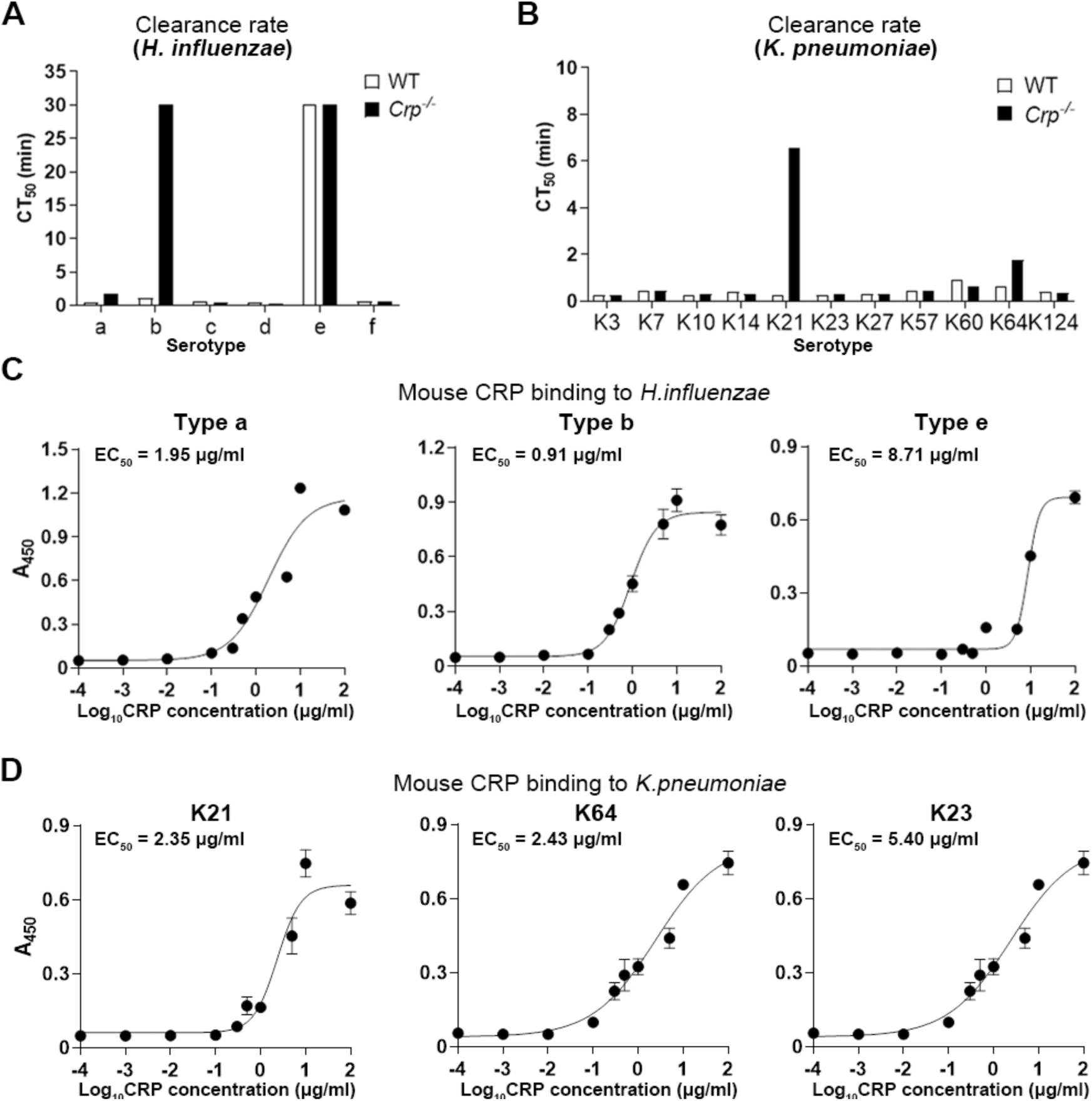
The contribution of CRP to the clearance of multiple Gram-negative pathogens. **A.** The role of CRP in the clearance of *H. influenzae* from bloodstream. WT and *Crp^-/-^* mice were i.v. infected with 10^7^ CFU. CT_50_ were calculated as in Figure S2A. n = 1-3. **B.** The role of CRP in the clearance of *K. pneumoniae* from bloodstream. WT and *Crp^-/-^* mice were i.v. infected with 5 × 10^6^ CFU, CT_50_ were assessed and presented as in (A). n = 1. **C.** r-mCRP binding to serotype a, b, and e of *H. influenzae* was assessed as in Figure 1D. n = 3. **D.** r-mCRP binding to K21, K64, and K23 of *K. pneumoniae* was assessed as in Figure 1D. n = 3.

**Figure S4.**
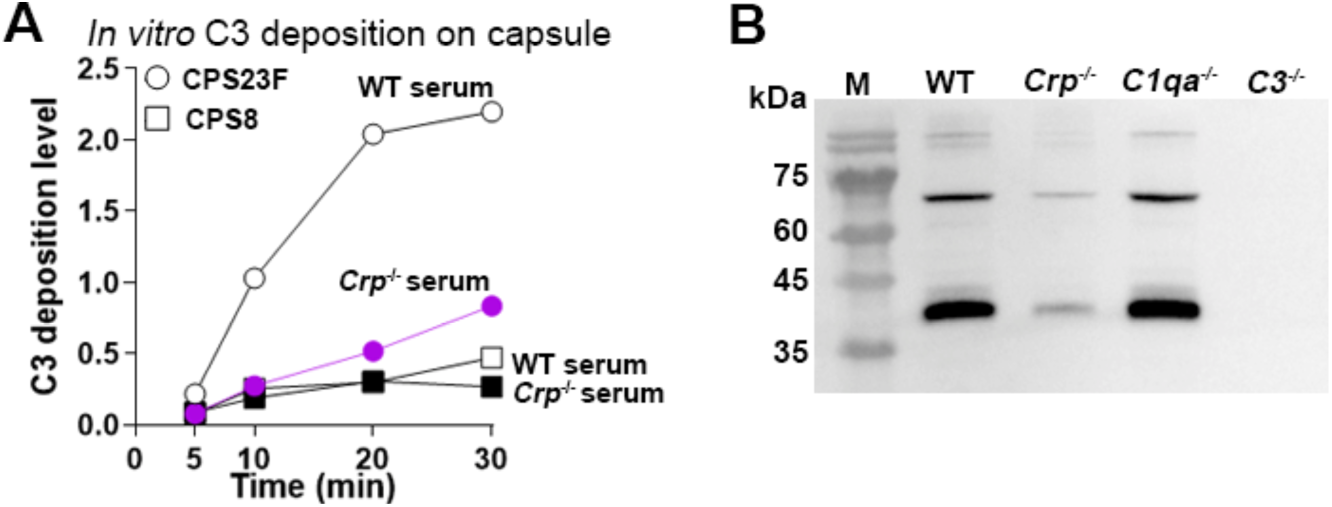
Evaluation of CRP-activated C3 activation on pneumococcal capsules. **A.** CRP-activated C3 deposition on free capsular polysaccharide of serotype-23F *S. pneumoniae* as detected by ELISA. The wells of 96-well plates were pre-coated with 10 μg/ml CPS23F or CPS8, and incubated with 100 μl of 10% *c* at 37 ℃ for various durations. The abundance of C3 bound to capsular polysaccharides was detected with anti-C3 antibody. n = 4. **B.** CRP-activated C3 deposition on the capsule of serotype-23F *S. pneumoniae* as detected by Western blotting. C3 activation levels on *Sp*23F after incubation with WT, *Crp*^-/-^, or *C1qa*^-/-^ serum. *C3*^-/-^ serum was used as a negative control. 10^7^ CFU of *Sp*23F were incubated with 100 μl of 10% *Crp^-/-^*, *C1qa*^-/-^ or WT mouse serum at 37 ℃ for 20 min. After extensive washing with PBS, pelleted bacteria were resuspended in the SDS-PAGE loading buffer; soluble proteins after centrifugation separated by protein electrophoresis; C3 detected with anti-C3 antibody.

**Figure S5.**
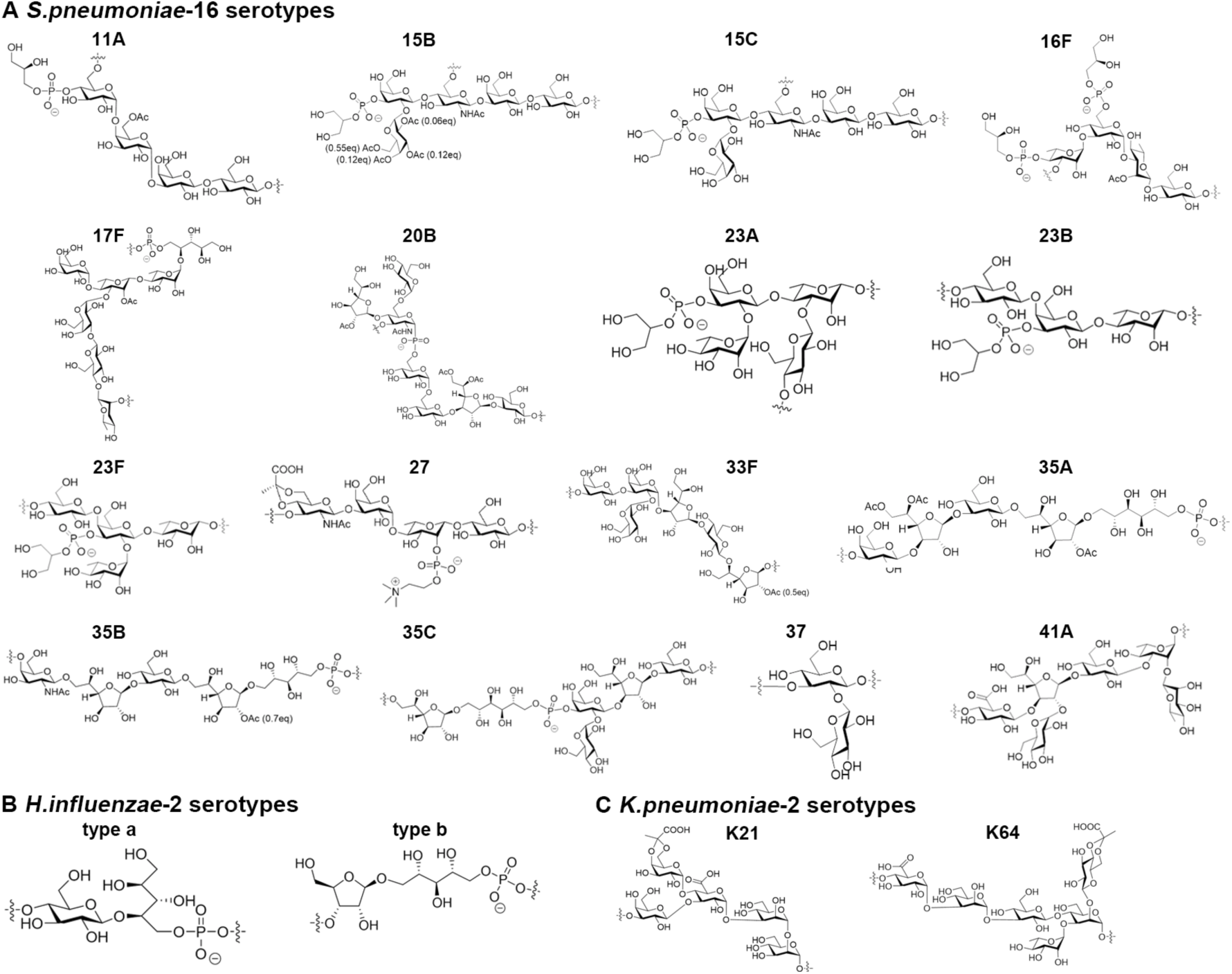
Chemical architectures of CRP-binding capsules. **A.** Chemical architectures of CRP-binding pneumococcal capsules (CPS11A, 15B, 15C, 16F, 17F, 20B, 23A, 23B, 23F, 27, 33F, 35A, 35B, 35C, 37 and 41A). **B.** Chemical architectures of *H. influenzae* types a and b capsules. **C.** Chemical architectures of *K. pneumoniae* type K21 and type K64 capsules.

**Figure S6.**
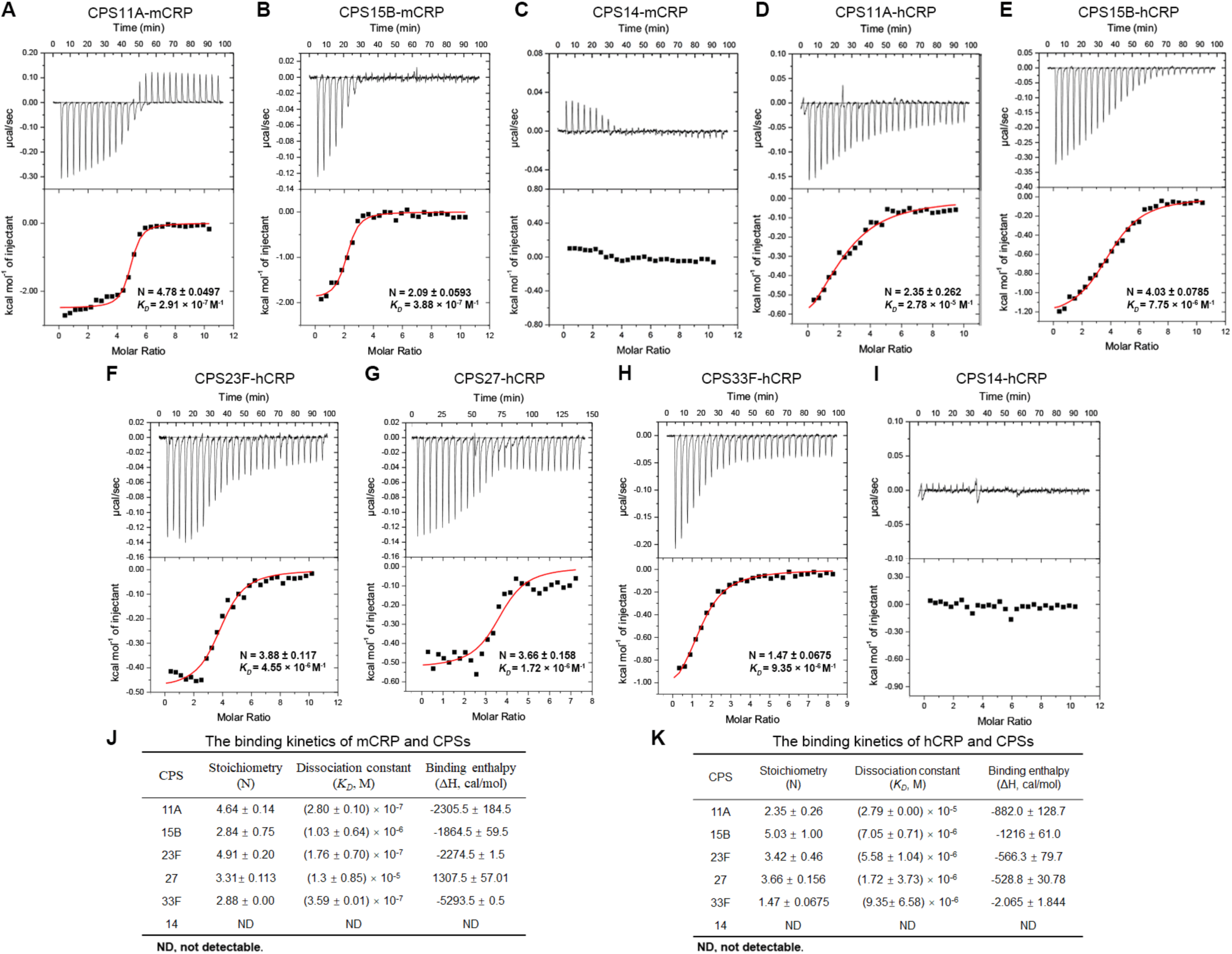
The binding affinity of CRP to capsules. **A-I.** Isothermal Titration Calorimetry (ITC) data for the interaction of CPS11A-mCRP (A), CPS15B-mCRP (B), CPS14-mCRP (C), CPS11A-hCRP (D), CPS15B-hCRP (E), CPS23F-hCRP (F), CPS27-hCRP (G), CPS33F-hCRP (H), and CPS14-hCRP (I). Data after base-line integration and concentration normalization were shown and the stoichiometry (N) and the dissociation constant (*K_D_*) were indicated in the plot. CPS14, negative control. n = 2. **J.** The summary of ITC data for the interaction of mCRP and capsules. **K.** The summary of ITC data for the interaction of hCRP and capsules.

**Figure S7.**
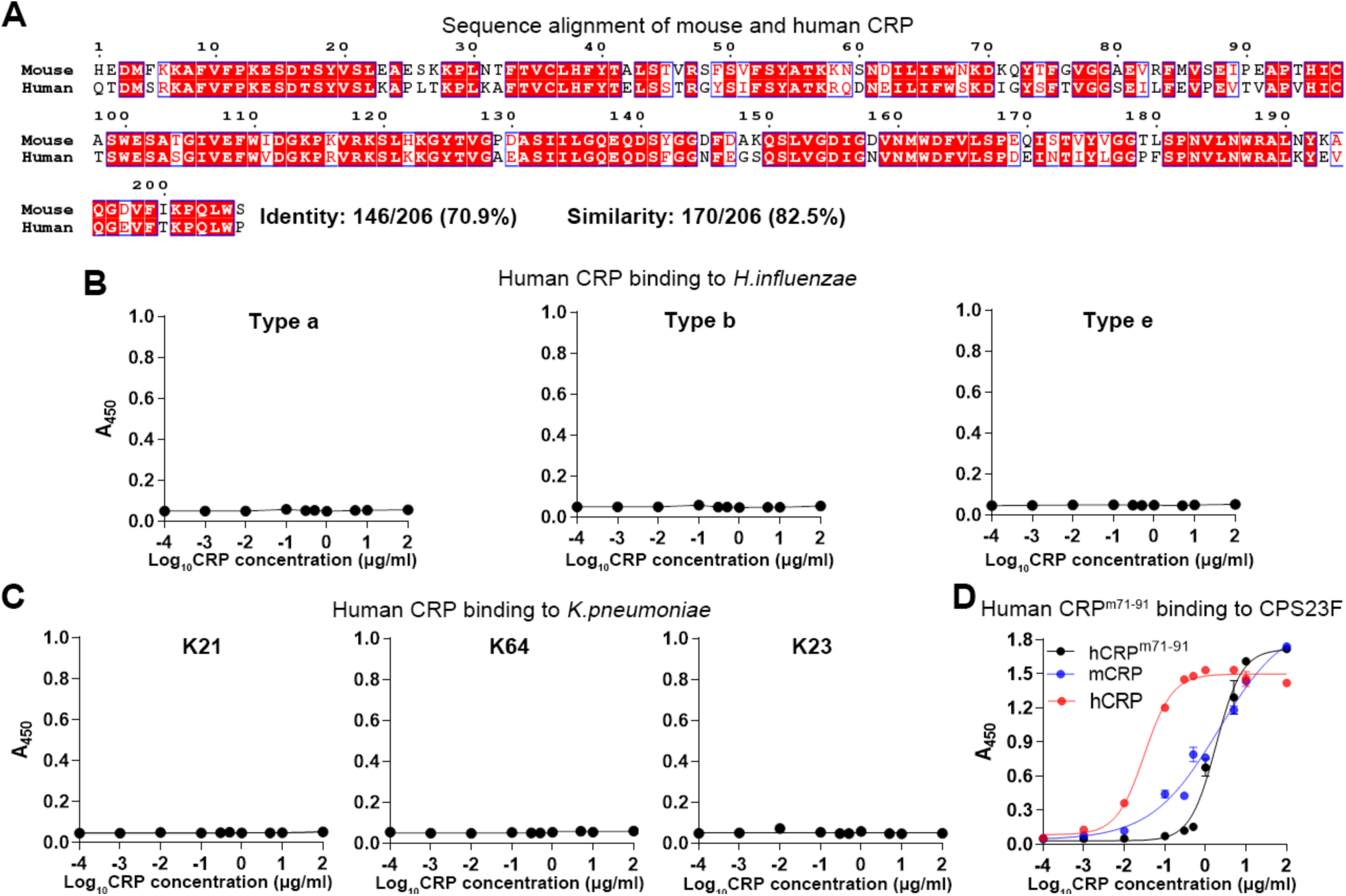
Human CRP mediates broad serotype-specific recognition of capsules. **A.** Sequence comparison between hCRP and mCRP. The amino acid sequences (without signal peptide) from mouse and human CRP were aligned by ClustalW software, and then the alignment results were imported into ENDscript/ESPript website to create alignment figure. The conserved amino acids were indicted with red highlights. **B, C.** r-hCRP binding to *H. influenzae* and *K. pneumoniae*, respectively. CPS-coated wells were incubated with different concentration of hCRP as in Figure 1H. n = 3. **D.** CPS23F binding of a human-mouse CRP hybrid. CPS23F binding of wild type hCRP, mCRP and human-mouse CRP hybrid (m71-91) was determined and presented as in Figure 6I. n = 3.

**Figure S8.**
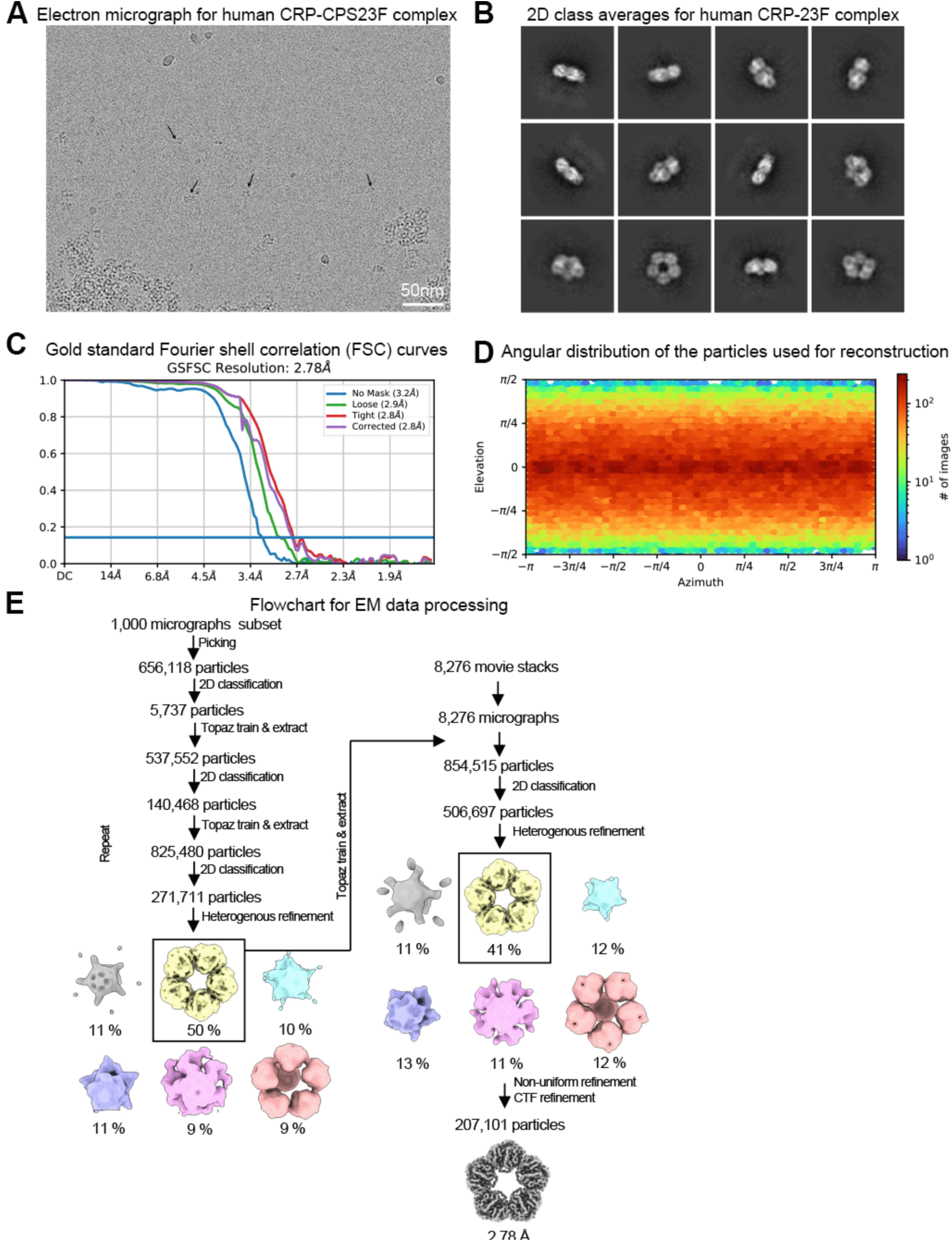
Cryo-EM structural analysis of the human CRP and CPS23F complex. **A.** Representative electron micrograph for the CRP-CPS23F complex. **B.** Representative 2D class averages for the CRP-CPS23F complex. **C.** Gold standard Fourier shell correlation (FSC) curves. **D.** Angular distribution of the particles used for reconstruction. **E.** Flowchart for EM data processing. Details can be found in Materials and Methods.

## SUPPLEMENTAL INFORMATION

**Table S1.** Bacterial information.

**Table S2.** CPS23F-binding mouse protein candidates identified by mass spectrometry*^a^*.

**Table S3.** Information of strains with recombinant protein plasmids.

**Table S4.** Primers used in this study.

**Table S5.** CPS23F-binding human protein candidates identified by mass spectrometry*^a^*.

**Table S6.** Cryo-EM data processing and refinement statistics.

**Movie S1.** IVM visualization of CRP function in KC-mediated *Sp*23F capture.

**Movie S2.** Unaffected KC capture of *Sp*14 in CPR-deficient mice.

**Movie S3.** Visualization of CRP-mediated capture of *Sp*11A by KCs in the liver.

**Movie S4.** Visualization of CRP-mediated capture of *Sp*20B by KCs in the liver.

**Movie S5.** Visualization of CRP-mediated capture of *Sp*23A by KCs in the liver.

**Movie S6.** Visualization of CRP-mediated capture of *Sp*33F by KCs in the liver.

**Movie S7.** Visualization of CRP-mediated capture of *Sp*35B by KCs in the liver.

**Movie S8.** Diminished KC capture of *Hib* in CRP-deficient mice.

**Movie S9.** CRP-mediated capture of *Kp*-K21 by KCs.

**Movie S10.** Comparison of KC-mediated *Sp*23F capture of WT, *C3*^-/-^, and CR3/CRIg^-/-^ mice.

## REFERENCES

1. Tillett, W.S., and Francis, T. (1930). Serological reactions in pneumonia with a non-protein somatic fraction of pneumococcus. J Exp Med 52, 561–571. 10.1084/jem.52.4.561.

2. Mold, C., Edwards, K.M., and Gewurz, H. (1982). Effect of C-reactive protein on the complement-mediated stimulated of human neutrophils by *Streptococcus pneumoniae* serotypes 3 and 6. Infect Immun 37, 987–992. 10.1128/iai.37.3.987-992.1982.

3. Volanakis, J.E., and Kaplan, M.D. (1971). Specificity of C-reactive protein for choline phosphate residues of pneumococcal C-polysaccharide. Proc. Soc. Exp. Biol. Med. 136, 612–614. https://journals.sagepub.com/doi/abs/10.3181/00379727-136-35323.

4. Kolberg, J., Hoiby, E.A., and Jantzen, E. (1997). Detection of the phosphorylcholine epitope in streptococci, *Haemophilus* and pathogenic *Neisseriae* by immunoblotting. Microb Pathog 22, 321–329. 10.1006/mpat.1996.0114.

5. Weiser, J.N., Pan, N., McGowan, K.L., Musher, D., Martin, A., and Richards, J. (1998). Phosphorylcholine on the lipopolysaccharide of *Haemophilus influenzae* contributes to persistence in the respiratory tract and sensitivity to serum killing mediated by C-reactive protein. J Exp Med 187, 631–640. 10.1084/jem.187.4.631.

6. Plebani, M. (2023). Why C-reactive protein is one of the most requested tests in clinical laboratories? Clin Chem Lab Med 61, 1540–1545. 10.1515/cclm-2023-0086.

7. Ji, S.R., Zhang, S.H., Chang, Y., Li, H.Y., Wang, M.Y., Lv, J.M., Zhu, L., Tang, P.M.K., and Wu, Y. (2023). C-reactive protein: the most familiar stranger. J Immunol 210, 699–707. 10.4049/jimmunol.2200831.

8. Eklund, C., Huttunen, R., Syrjanen, J., Laine, J., Vuento, R., and Hurme, M. (2006). Polymorphism of the C-reactive protein gene is associated with mortality in bacteraemia. Scand J Infect Dis 38, 1069–1073. 10.1080/00365540600978922.

9. Mold, C., Nakayama, S., Holzer, T.J., Gewurz, H., and Duclos, T.W. (1981). C-reactive protein is protective against *Streptococcus pneumoniae* infection in mice. Journal of Experimental Medicine 154, 1703–1708. 10.1084/jem.154.5.1703.

10. Ngwa, D.N., Singh, S.K., and Agrawal, A. (2020). C-reactive protein-based strategy to reduce antibiotic dosing for the treatment of pneumococcal infection. Front Immunol 11, 620784. 10.3389/fimmu.2020.620784.

11. Ngwa, D.N., Singh, S.K., Gang, T.B., and Agrawal, A. (2020). Treatment of pneumococcal infection by using engineered human C-reactive protein in a mouse model. Front Immunol 11, 586669. 10.3389/fimmu.2020.586669.

12. Szalai, A.J., Briles, D.E., and Volanakis, J.E. (1995). Human C-reactive protein is protective against fatal *Streptococcus pneumoniae* infection in transgenic mice. J Immunol 155, 2557–2563. 10.4049/jimmunol.155.5.2557s.

13. Yother, J., Volanakis, J.E., and Briles, D.E. (1982). Human C-reactive protein is protective against fatal *Streptococcus pneumoniae* infection in mice. Journal of Immunology 128, 2374–2376. 10.4049/jimmunol.128.5.2374.

14. Holzer, T.J., Edwards, K.M., Gewurz, H., and Mold, C. (1984). Binding of C-reactive protein to the pneumococcal capsule or cell wall results in differential localization of C3 and stimulation of phagocytosis. J Immunol 133, 1424–1430. 10.4049/jimmunol.133.3.1424.

15. Shrive, A.K., Cheetham, G.M., Holden, D., Myles, D.A., Turnell, W.G., Volanakis, J.E., Pepys, M.B., Bloomer, A.C., and Greenhough, T.J. (1996). Three dimensional structure of human C-reactive protein. Nat Struct Biol 3, 346–354. 10.1038/nsb0496-346.

16. Crowell, R.E., Du Clos, T.W., Montoya, G., Heaphy, E., and Mold, C. (1991). C-reactive protein receptors on the human monocytic cell line U-937. Evidence for additional binding to Fc gamma RI. J Immunol 147, 3445–3451. 10.4049/jimmunol.147.10.3445.

17. Bharadwaj, D., Stein, M.P., Volzer, M., Mold, C., and Du Clos, T.W. (1999). The major receptor for C-reactive protein on leukocytes is fcgamma receptor II. J Exp Med 190, 585–590. 10.1084/jem.190.4.585.

18. Marnell, L.L., Mold, C., Volzer, M.A., Burlingame, R.W., and Du Clos, T.W. (1995). C-reactive protein binds to Fc gamma RI in transfected COS cells. J Immunol 155, 2185–2193. 10.4049/jimmunol.155.4.2185.

19. Kaplan, M.H., and Volanakis, J.E. (1974). Interaction of C-reactive protein complexes with the complement system: I. consumption of human complement associated with the reaction of C-reactive protein with pneumococcal C-polysaccharide and with the choline phosphatides, lecithin and sphingomyelin. J Immunol 112, 2135–2147. 10.4049/jimmunol.112.6.2135.

20. Noone, D.P., van der Velden, T.T., and Sharp, T.H. (2021). Cryo-electron microscopy and biochemical analysis offer insights into the effects of acidic pH, such as occur during acidosis, on the complement binding properties of C-reactive protein. Front Immunol 12, 757633. 10.3389/fimmu.2021.757633.

21. Guillon, C., Bigouagou, U.M., Folio, C., Jeannin, P., Delneste, Y., and Gouet, P. (2014). A staggered decameric assembly of human C-reactive protein stabilized by zinc ions revealed by X-ray crystallography. Protein Pept Lett 22, 248–255. 10.2174/0929866522666141231111226.

22. Mortensen, R.F., Osmand, A.P., Lint, T.F., and Gewurz, H. (1976). Interaction of C-reactive protein with lymphocytes and monocytes: complement-dependent adherence and phagocytosis. J Immunol 117, 774–781. 10.4049/jimmunol.117.3.774.

23. Sproston, N.R., and Ashworth, J.J. (2018). Role of C-reactive protein at sites of inflammation and infection. Front Immunol 9, 1–11. 10.3389/fimmu.2018.00754.

24. Mold, C., Rodic-Polic, B., and Du Clos, T.W. (2002). Protection from *Streptococcus pneumoniae* infection by C-reactive protein and natural antibody requires complement but not Fc gamma receptors. J Immunol 168, 6375–6381. 10.4049/jimmunol.168.12.6375.

25. Szalai, A.J., Briles, D.E., and Volanakis, J.E. (1996). Role of complement in C-reactive-protein-mediated protection of mice from *Streptococcus pneumoniae*. Infect Immun 64, 4850–4853. 10.1128/iai.64.11.4850-4853.1996.

26. Naghavi, M., and Collaborators, G.A.R. (2022). Global mortality associated with 33 bacterial pathogens in 2019: a systematic analysis for the global burden of disease study 2019. Lancet 400, 2221–2248. 10.1016/S0140-6736(22)02185-7.

27. An, H., Liu, Y., Qian, C., Huang, X., Wang, L., Whitfield, C., and Zhang, J.-R. (2024). Bacterial capsules. In Molecular Medical Microbiology, Y.-W. Tang, M. Hindiyeh, D. Liu, A. Salis, P. Spearman, and J.-R. Zhang, eds. (Academic Press), pp. 69–96. 10.1016/B978-0-12-818619-0.00150-7.

28. Taylor, C., and Roberts, I.S. (2002). The regulation of capsule expression. In Bacterial Adhesion to Host tissues-Mechanisms and Consequences, M. Wilson, ed. (Cambridge University Press), pp. 115–138.

29. Brown, E.J., and Gresham, H.D. (2012). Phagocytosis. In Fundamental Immunology, W. E.P., ed. (Lippincott-Raven Publishers), pp. 1105–1127.

30. Nahm, M.H., and Katz, J. (2012). Immunity to extracellular bacteria. In Fundamental Immunology, W. E.P., ed. (Lippincott-Raven Publishers), pp. 1001–1015.

31. Whitfield, C., Wear, S.S., and Sande, C. (2020). Assembly of bacterial capsular polysaccharides and exopolysaccharides. Annu Rev Microbiol 74, 521–543. 10.1146/annurev-micro-011420-075607.

32. Manna, S., Werren, J.P., Ortika, B.D., Bellich, B., Pell, C.L., Nikolaou, E., Gjuroski, I., Lo, S., Hinds, J., Tundev, O., et al. (2024). *Streptococcus pneumoniae* serotype 33G: genetic, serological, and structural analysis of a new capsule type. Microbiol Spectr 12, e0357923. 10.1128/spectrum.03579-23.

33. Wyres, K.L., Wick, R.R., Gorrie, C., Jenney, A., Follador, R., Thomson, N.R., and Holt, K.E. (2016). Identification of *Klebsiella* capsule synthesis loci from whole genome data. Microb Genom 2, e000102. 10.1099/mgen.0.000102.

34. White, B. (1938). Pathogenicity of pneumococcus: man. In The Biology of Pneumococcus, (Harvard University Press), pp. 214–237.

35. Retchless, A.C., Topaz, N., Marjuki, H., Marasini, D., Potts, C.C., and Wang, X. (2024). *Haemophilus influenzae*. In Molecular Medical Microbiology, Y.-W. Tang, M. Hindiyeh, D. Liu, A. Salis, P. Spearman, and J.-R. Zhang, eds. (Academic Press), pp. 1399–1421. 10.1016/B978-0-12-818619-0.00129-5.

36. Bilzer, M., Roggel, F., and Gerbes, A.L. (2006). Role of Kupffer cells in host defense and liver disease. Liver Int 26, 1175–1186. 10.1111/j.1478-3231.2006.01342.x.

37. Helmy, K.Y., Katschke, K.J., Jr., Gorgani, N.N., Kljavin, N.M., Elliott, J.M., Diehl, L., Scales, S.J., Ghilardi, N., and van Lookeren Campagne, M. (2006). CRIg: a macrophage complement receptor required for phagocytosis of circulating pathogens. Cell 124, 915–927. 10.1016/j.cell.2005.12.039.

38. Zeng, Z.T., Surewaard, B.G.J., Wong, C.H.Y., Geoghegan, J.A., Jenne, C.N., and Kubes, P. (2016). CRIg functions as a macrophage pattern recognition receptor to directly bind and capture blood-borne gram-positive bacteria. Cell Host Microbe 20, 99–106. 10.1016/j.chom.2016.06.002.

39. Broadley, S.P., Plaumann, A., Coletti, R., Lehmann, C., Wanisch, A., Seidlmeier, A., Esser, K., Luo, S., Ramer, P.C., Massberg, S., et al. (2016). Dual-track clearance of circulating bacteria balances rapid restoration of blood sterility with induction of adaptive immunity. Cell Host Microbe 20, 36–48. 10.1016/j.chom.2016.05.023.

40. Wong, C.H., Jenne, C.N., Petri, B., Chrobok, N.L., and Kubes, P. (2013). Nucleation of platelets with blood-borne pathogens on Kupffer cells precedes other innate immunity and contributes to bacterial clearance. Nat Immunol 14, 785–792. 10.1038/ni.2631.

41. Lee, W.Y., Moriarty, T.J., Wong, C.H., Zhou, H., Strieter, R.M., van Rooijen, N., Chaconas, G., and Kubes, P. (2010). An intravascular immune response to Borrelia burgdorferi involves Kupffer cells and iNKT cells. Nat Immunol 11, 295–302. 10.1038/ni.1855.

42. Surewaard, B.G., Deniset, J.F., Zemp, F.J., Amrein, M., Otto, M., Conly, J., Omri, A., Yates, R.M., and Kubes, P. (2016). Identification and treatment of the *Staphylococcus aureus* reservoir in vivo. J Exp Med 213, 1141–1151. 10.1084/jem.20160334.

43. Kolaczkowska, E., Jenne, C.N., Surewaard, B.G., Thanabalasuriar, A., Lee, W.Y., Sanz, M.J., Mowen, K., Opdenakker, G., and Kubes, P. (2015). Molecular mechanisms of NET formation and degradation revealed by intravital imaging in the liver vasculature. Nat Commun 6, 6673. 10.1038/ncomms7673.

44. Gola, A., Dorrington, M.G., Speranza, E., Sala, C., Shih, R.M., Radtke, A.J., Wong, H.S., Baptista, A.P., Hernandez, J.M., Castellani, G., et al. (2021). Commensal-driven immune zonation of the liver promotes host defence. Nature 589, 131–136. 10.1038/s41586-020-2977-2.

45. An, H., Qian, C., Huang, Y., Li, J., Tian, X., Feng, J., Hu, J., Fang, Y., Jiao, F., Zeng, Y., et al. (2022). Functional vulnerability of liver macrophages to capsules defines virulence of blood-borne bacteria. J Exp Med 219, e20212032. 10.1084/jem.20212032.

46. Huang, X., Li, X., An, H., Wang, J., Ding, M., Wang, L., Li, L., Ji, Q., Qu, F., Wang, H., et al. (2022). Capsule type defines the capability of *Klebsiella pneumoniae* in evading Kupffer cell capture in the liver. PLoS Pathog 18, e1010693. 10.1371/journal.ppat.1010693.

47. Silva-Costa, C., Melo-Cristino, J., and Ramirez, M. (2024). *Streptococcus pneumoniae*. In Molecular Medical Microbiology, Y.-W. Tang, M. Hindiyeh, D. Liu, A. Salis, P. Spearman, and J.-R. Zhang, eds. (Academic Press), pp. 1479–1485. 10.1016/B978-0-12-818619-0.00095-2.

48. Paczosa, M.K., and Mecsas, J. (2016). *Klebsiella pneumoniae*: going on the offense with a strong defense. Microbiology and Molecular Biology Reviews 80, 629–661. 10.1128/Mmbr.00078-15.

49. Haapasalo, K., and Meri, S. (2019). Regulation of the complement system by pentraxins. Front Immunol 10, 1750. 10.3389/fimmu.2019.01750.

50. Densen, P., and Ram, S. (2015). Complement and deficiencies. In Mandell, Douglas, and Bennett’s Principles and Practice of Infectious Diseases, J.E. Bennett, R.D. Dolin, and M.J. Blaser, eds. (Elsevier Churchill Livingstone), pp. 93–115.

51. Walbaum, S., Ambrosy, B., Schutz, P., Bachg, A.C., Horsthemke, M., Leusen, J.H.W., Mocsai, A., and Hanley, P.J. (2021). Complement receptor 3 mediates both sinking phagocytosis and phagocytic cup formation via distinct mechanisms. J Biol Chem 296, 100256. 10.1016/j.jbc.2021.100256.

52. Claus, D.R., Siegel, J., Petras, K., Osmand, A.P., and Gewurz, H. (1977). Interactions of C-reactive protein with the first component of human complement. J Immunol 119, 187–192.

53. Thompson, D., Pepys, M.B., and Wood, S.P. (1999). The physiological structure of human C-reactive protein and its complex with phosphocholine. Structure 7, 169–177. 10.1016/S0969-2126(99)80023-9.

54. Goda, T., and Miyahara, Y. (2017). Calcium-independent binding of human C-reactive protein to lysophosphatidylcholine in supported planar phospholipid monolayers. Acta Biomater 48, 206–214. 10.1016/j.actbio.2016.10.043.

55. Ramadan, M.A., Shrive, A.K., Holden, D., Myles, D.A., Volanakis, J.E., DeLucas, L.J., and Greenhough, T.J. (2002). The three-dimensional structure of calcium-depleted human C-reactive protein from perfectly twinned crystals. Acta Crystallogr D Biol Crystallogr 58, 992–1001. 10.1107/s0907444902005693.

56. Christopeit, T., Gossas, T., and Danielson, U.H. (2009). Characterization of Ca^2+^ and phosphocholine interactions with C-reactive protein using a surface plasmon resonance biosensor. Analytical Biochemistry 391, 39–44. 10.1016/j.ab.2009.04.037.

57. Lee, R.T., Takagahara, I., and Lee, Y.C. (2002). Mapping the binding areas of human C-reactive protein for phosphorylcholine and polycationic compounds: relationship between the two types of binding sites. J Biol Chem 277, 225–232. 10.1074/jbc.M106039200.

58. Bray, C., Bell, L.N., Liang, H., Haykal, R., Kaiksow, F., Mazza, J.J., and Yale, S.H. (2016). Erythrocyte sedimentation rate and C-reactive protein measurements and their relevance in clinical medicine. WMJ 115, 317–321. https://www.ncbi.nlm.nih.gov/pubmed/29094869.

59. Krarup, A., Thiel, S., Hansen, A., Fujita, T., and Jensenius, J.C. (2004). L-ficolin is a pattern recognition molecule specific for acetyl groups. J Biol Chem 279, 47513–47519. 10.1074/jbc.M407161200.

60. Krarup, A., Sorensen, U.B., Matsushita, M., Jensenius, J.C., and Thiel, S. (2005). Effect of capsulation of opportunistic pathogenic bacteria on binding of the pattern recognition molecules mannan-binding lectin, L-ficolin, and H-ficolin. Infect Immun 73, 1052–1060. 10.1128/iai.73.2.1052-1060.2005.

61. Brady, A.M., Calix, J.J., Yu, J., Geno, K.A., Cutter, G.R., and Nahm, M.H. (2014). Low invasiveness of pneumococcal serotype 11A is linked to ficolin-2 recognition of O-acetylated capsule epitopes and lectin complement pathway activation. J Infect Dis 210, 1155–1165. 10.1093/infdis/jiu195.

62. Geno, K.A., Spencer, B.L., Bae, S., and Nahm, M.H. (2018). Ficolin-2 binds to serotype 35B pneumococcus as it does to serotypes 11A and 31, and these serotypes cause more infections in older adults than in children. PLoS One 13, e0209657. 10.1371/journal.pone.0209657.

63. Nahm, M.H., Yu, J., Calix, J.J., and Ganaie, F. (2022). Ficolin-2 lectin complement pathway mediates capsule-specific innate immunity against invasive pneumococcal disease. Front Immunol 13, 841062. 10.3389/fimmu.2022.841062.

64. Neth, O., Jack, D.L., Dodds, A.W., Holzel, H., Klein, N.J., and Turner, M.W. (2000). Mannose-binding lectin binds to a range of clinically relevant microorganisms and promotes complement deposition. Infect Immun 68, 688–693. 10.1128/iai.68.2.688-693.2000.

65. Edwards, K.M., Gewurz, H., Lint, T.F., and Mold, C. (1982). A role for C-reactive protein in the complement-mediated stimulation of human neutrophils by type 27 *Streptococcus pneumoniae*. J Immunol 128, 2493–2496. 10.4049/jimmunol.128.6.2493.

66. Ngwa, D.N., and Agrawal, A. (2019). Structure-function relationships of C-reactive protein in bacterial infection. Front Immunol 10, 166. 10.3389/fimmu.2019.00166.

67. Huang, Y., Li, K., Huang, X., Tian, X., Meng, J., Zhou, H., Wu, J., Dai, Q., Zhang, J.-R., and An, H. (2024). Splenic red pulp macrophages eliminate the liver-resistant *Streptococcus pneumoniae* from the blood circulation. bioRxiv. 10.1101/2024.04.26.591417.

68. Wang, J., An, H., Ding, M., Liu, Y., Wang, S., Jin, Q., Wu, Q., Dong, H., Guo, Q., Tian, X., et al. (2023). Liver macrophages and sinusoidal endothelial cells execute vaccine-elicited capture of invasive bacteria. Sci Transl Med 15, eade0054. 10.1126/scitranslmed.ade0054.

69. Singh, S.K., Ngwa, D.N., and Agrawal, A. (2020). Complement activation by C-reactive protein is critical for protection of mice against pneumococcal infection. Front Immunol 11, 1812. 10.3389/fimmu.2020.01812.

70. Suresh, M.V., Singh, S.K., Ferguson, D.A., Jr., and Agrawal, A. (2006). Role of the property of C-reactive protein to activate the classical pathway of complement in protecting mice from pneumococcal infection. J Immunol 176, 4369–4374. 10.4049/jimmunol.176.7.4369.

71. Chudwin, D.S., Artrip, S.G., Korenblit, A., Schiffman, G., and Rao, S. (1985). Correlation of serum opsonins with in vitro phagocytosis of *Streptococcus pneumoniae*. Infect Immun 50, 213–217. 10.1128/iai.50.1.213-217.1985.

72. Agrawal, A., Shrive, A.K., Greenhough, T.J., and Volanakis, J.E. (2001). Topology and structure of the C1q-binding site on C-reactive protein. J Immunol 166, 3998–4004. 10.4049/jimmunol.166.6.3998.

73. Zurawski, D.V., and McLendon, M.K. (2020). Monoclonal antibodies as an antibacterial approach against bacterial pathogens. Antibiotics (Basel) 9. 10.3390/antibiotics9040155.

74. Simonis, A., Kreer, C., Albus, A., Rox, K., Yuan, B., Holzmann, D., Wilms, J.A., Zuber, S., Kottege, L., Winter, S., et al. (2023). Discovery of highly neutralizing human antibodies targeting *Pseudomonas aeruginosa*. Cell 186, 5098–5113 e5019. 10.1016/j.cell.2023.10.002.

75. Smith, P., DiLillo, D.J., Bournazos, S., Li, F., and Ravetch, J.V. (2012). Mouse model recapitulating human Fcγ receptor structural and functional diversity. Proceedings of the National Academy of Sciences 109, 6181–6186. https://www.pnas.org/doi/epdf/10.1073/pnas.1203954109.

76. Lu, L., Ma, Y., and Zhang, J.R. (2006). *Streptococcus pneumoniae* recruits complement factor H through the amino terminus of CbpA. J Biol Chem 281, 15464–15474. 10.1074/jbc.M602404200.

77. Weiser, J.N., Love, J.M., and Moxon, E.R. (1989). The molecular mechanism of phase variation of *H. influenzae* lipopolysaccharide. Cell 59, 657–665. 10.1016/0092-8674(89)90011-1.

78. An, H., Tian, X., Huang, Y., and Zhang, J.R. (2023). Identification of the mouse Kupffer cell receptors recognizing pneumococcal capsules by affinity screening. Star Protocols 4, 102065. 10.1016/j.xpro.2023.102065.

79. Cao, D., Ma, B., Cao, Z., Zhang, X., and Xiang, Y. (2023). Structure of Semliki Forest virus in complex with its receptor VLDLR. Cell 186, 2208–2218 e2215. 10.1016/j.cell.2023.03.032.

80. Hemsley, A., Arnheim, N., Toney, M.D., Cortopassi, G., and Galas, D.J. (1989). A simple method for site-directed mutagenesis using the polymerase chain reaction. Nucleic Acids Res 17, 6545–6551. 10.1093/nar/17.16.6545.

81. Tian, X., Liu, y., Zhu, K., An, H., Feng, J., Zhang, L., and Zhang, J.R. (2024). Natural antibodies to polysaccharide capsules enable Kupffer cells to capture invading bacteria in the liver sinusoids. bioRxiv. 10.1101/2024.04.26.591254.

82. Zheng, S.Q., Palovcak, E., Armache, J.P., Verba, K.A., Cheng, Y., and Agard, D.A. (2017). MotionCor2: anisotropic correction of beam-induced motion for improved cryo-electron microscopy. Nat Methods 14, 331–332. 10.1038/nmeth.4193.

83. Punjani, A., Rubinstein, J.L., Fleet, D.J., and Brubaker, M.A. (2017). cryoSPARC: algorithms for rapid unsupervised cryo-EM structure determination. Nat Methods 14, 290–296. 10.1038/nmeth.4169.

84. Sanchez-Garcia, R., Gomez-Blanco, J., Cuervo, A., Carazo, J.M., Sorzano, C.O.S., and Vargas, J. (2021). DeepEMhancer: a deep learning solution for cryo-EM volume post-processing. Commun Biol 4, 874. 10.1038/s42003-021-02399-1.

85. Bepler, T., Morin, A., Rapp, M., Brasch, J., Shapiro, L., Noble, A.J., and Berger, B. (2019). Positive-unlabeled convolutional neural networks for particle picking in cryo-electron micrographs. Nat Methods 16, 1153–1160. 10.1038/s41592-019-0575-8.

86. Pettersen, E.F., Goddard, T.D., Huang, C.C., Meng, E.C., Couch, G.S., Croll, T.I., Morris, J.H., and Ferrin, T.E. (2021). UCSF ChimeraX: structure visualization for researchers, educators, and developers. Protein Sci 30, 70–82. 10.1002/pro.3943.

87. Emsley, P., and Cowtan, K. (2004). Coot: model-building tools for molecular graphics. Acta Crystallogr D Biol Crystallogr 60, 2126–2132. 10.1107/S0907444904019158.

88. Adams, P.D., Afonine, P.V., Bunkoczi, G., Chen, V.B., Davis, I.W., Echols, N., Headd, J.J., Hung, L.W., Kapral, G.J., Grosse-Kunstleve, R.W., et al. (2010). PHENIX: a comprehensive Python-based system for macromolecular structure solution. Acta Crystallogr D Biol Crystallogr 66, 213–221. 10.1107/S0907444909052925.

89. Chen, V.B., Arendall, W.B., 3rd, Headd, J.J., Keedy, D.A., Immormino, R.M., Kapral, G.J., Murray, L.W., Richardson, J.S., and Richardson, D.C. (2010). MolProbity: all-atom structure validation for macromolecular crystallography. Acta Crystallogr D Biol Crystallogr 66, 12–21. 10.1107/S0907444909042073.

90. Jumper, J., Evans, R., Pritzel, A., Green, T., Figurnov, M., Ronneberger, O., Tunyasuvunakool, K., Bates, R., Žídek, A., Potapenko, A., et al. (2021). Highly accurate protein structure prediction with AlphaFold. Nature 596, 583–589. 10.1038/s41586-021-03819-2.

