## Supplemental Table S1 for "C-Reactive Protein Drives Potent Clearance of Blood Bacteria in the Liver by Activating the Complement System"

**Table S1. Strains used in this study**

| Strain ID | Genotype | Capsule type | CT50 in WT mouse (min) | CT50 in *Crp^-/-^* mouse (min) | Source | Region |
| --- | --- | --- | --- | --- | --- | --- |
| *S. pneumoniae* |  |  |  |  |  |  |
| TH15943 | TH870Δ*cps*::*cps*8 | 8 | 18.4 | - | - | - |
| TH15984 | Wildtype | 9N | 2.13 | 1.7 | Sputum | China |
| TH13133 | TH870Δ*cps*::*cps*9V | 9V | 1.46 | 2.12 | - | - |
| TH2594 | Wild type | 10A | 1.28 | 1.06 | Blood | China |
| TH12932 | Wild type | 11A | 1.49 | >30 | Sputum | China |
| TH15944 | TH870Δ*cps*::*cps*14 | 14 | 0.57 | 0.47 | - | - |
| TH2941 | Wild type | 15B | 0.97 | >30 | Blood | China |
| TH886 | Wild type | 15C | 0.9 | >30 | CSF | USA |
| TH16784 | Wild type | 16F | 0.79 | >30 | Sputum | China |
| TH2734 | Wild type | 17F | 4.59 | 24.9 | Blood | China |
| TH14190 | TH870Δ*cps*::*cps*19A | 19A | 1.26 | 2.03 | - | - |
| TH14188 | TH870Δ*cps*::*cps*19F | 19F | 1.12 | 1.28 | - | - |
| TH2591 | Wild type | 20B | 1.8 | 28.89 | Blood | China |
| TH16792 | Wild type | 21 | 1.32 | 18.56 | Sputum | China |
| TH16798 | Wild type | 23A | 2.87 | >30 | Blood | China |
| TH901 | Wild type | 23B | 0.87 | 2.06 | Blood | USA |
| TH15945 | TH870Δ*cps*::*cps*23F | 23F | 1.31 | >30 | - | - |
| TH17250 | Wild type | 27 | 5.35 | 16.62 | Blood | China |
| TH874 | Wild type | 33F | 1.08 | >30 | Blood | USA |
| TH16814 | Wild type | 34 | 1.03 | 1.33 | Sputum | China |
| TH16815 | Wild type | 35A | 1.5 | >30 | Sputum | China |
| TH902 | Wild type | 35B | 11.03 | >30 | CSF | USA |
| TH16820 | Wild type | 35C | 1.2 | >30 | Sputum | China |
| TH16824 | Wild type | 37 | 6.71 | >30 | Sputum | China |
| TH16828 | Wild type | 41A | 9.62 | 11.5 | Sputum | China |
| TH16830 | Wild type | 48 | 1.19 | 1.01 | Sputum | China |
| Strain ID | Genotype | Capsule type | CT50 in WT mouse (min) | CT50 in *Crp^-/-^* mouse (min) | Source | Region |
| *H. influenzae* |  |  |  |  |  |  |
| 5558 (TH17228) | Wild type | a | 0.49 | 1.7 | - | - |
| M5216 (TH17189) | Wild type | b | 1.08 | >30 | - | USA |
| TH17229 | Wild type | c | 0.57 | 0.38 | Sputum | USA |
| TH17231 | Wild type | d | 0.38 | 0.31 | Throat | Netherlands |
| NCTC10479 (TH17230) | Wild type | e | >30 |  | - | - |
| TH17232 | Wild type | f | 0.57 | 0.61 | Throat | China |
| *K. pneumoniae* |  |  |  |  |  |  |
| TH12849 | Wild type | K3 | 0.27 | 0.27 | Blood | China |
| TH12880 | Wild type | K7 | 0.43 | 0.44 | Blood | China |
| TH13089 | Wild type | K10 | 0.28 | 0.31 | Bile | China |
| TH12838 | Wild type | K14 | 0.41 | 0.3 | Blood | China |
| TH13012 | Wild type | K21 | 0.24 | 6.55 | Tissue | China |
| TH12852 | Wild type | K23 | 0.28 | 0.29 | Blood | China |
| TH13098 | Wild type | K27 | 0.32 | 0.31 | Blood | China |
| TH13007 | Wild type | K57 | 0.46 | 0.43 | Blood | China |
| TH12855 | Wild type | K60 | 0.9 | 0.63 | Blood | China |
| TH12841 | Wild type | K64 | 0.62 | 1.76 | Blood | China |
| TH12879 | Wild type | K124 | 0.39 | 0.35 | Blood | China |
