## Supplemental Table S2 for "C-Reactive Protein Drives Potent Clearance of Blood Bacteria in the Liver by Activating the Complement System"

**Table S2. CPS23F-binding mouse protein candidates identified by mass spectrometry*^a^***

| Uniprot accession | Protein | Description | Abundance | | CPS23F:CPS8 fold |
| --- | --- | --- | --- | --- | --- |
|  |  |  | CPS23F*^b^* | CPS8 |  |
| P14847 | Crp | C-reactive protein | 6.335 × 10^10^ | 0 | ∞ |
| G3UY98 | Mettl3 | N6-adenosine-methyltransferase subunit | 1.001 × 10^10^ | 0 | ∞ |
| A0A0R4J213 | Slc14a2 | Urea transporter | 2.010 × 10^9^ | 0 | ∞ |
| Q8CFG8 | C1sb | Complement C1s-B subcomponent | 2.000 × 10^9^ | 0 | ∞ |
| Q8CG16 | C1ra | Complement C1r-A subcomponent | 1.790 × 10^9^ | 0 | ∞ |
| A0A077S9N1 | Lyz1 | 1,4-beta-N-acetylmuramidase C | 9.678 × 10^8^ | 0 | ∞ |
| Q497N1 | Rps26 | 40S ribosomal protein S26 | 1.296 × 10^8^ | 0 | ∞ |
| Q6IRU2 | Tpm4 | Tropomyosin alpha-4 chain | 1.261 × 10^8^ | 0 | ∞ |
| A0A1W2P7I2 | Epb41l2 | Band 4.1-like protein 2 | 1.006 × 10^8^ | 0 | ∞ |
| Q8K469 | Oas1g | 2'-5' oligoadenylate synthase | 9.195 × 10^7^ | 0 | ∞ |
| D3YXF5 | C7 | Complement component 7 | 9.184 × 10^7^ | 0 | ∞ |
| P39054-2 | Dnm2 | Isoform 2 of Dynamin-2 | 8.220 × 10^7^ | 0 | ∞ |
| G3X9G4 | Dnm2 | Dynamin GTPase | 8.220 × 10^7^ | 0 | ∞ |
| Q8BPB5 | Efemp1 | EGF-containing fibulin-like extracellular matrix protein 1 | 7.518 × 10^7^ | 0 | ∞ |
| Q8BUT2 | Dnm3 | Dynamin GTPase | 6.234 × 10^7^ | 0 | ∞ |

| Uniprot accession | Protein | Description | Abundance | | CPS23F:CPS8 fold |
| --- | --- | --- | --- | --- | --- |
|  |  |  | CPS23F | CPS8 |  |
| Q9DCU6 | Mrpl4 | 39S ribosomal protein L4, mitochondrial | 5.564 × 10^7^ | 0 | ∞ |
| Q9DBS2 | Tprg1l | Tumor protein p63-regulated gene 1-like protein | 4.830 × 10^7^ | 0 | ∞ |
| O08601 | Mttp | Microsomal triglyceride transfer protein large subunit | 4.129 × 10^7^ | 0 | ∞ |
| A0A1W2P8C0 | Epb41l2 | Band 4.1-like protein 2 (Fragment) | 3.798 × 10^7^ | 0 | ∞ |
| Q6ZPF4 | Fmnl3 | Formin-like protein 3 | 3.786 × 10^7^ | 0 | ∞ |
| Q99JB2 | Stoml2 | Stomatin-like protein 2, mitochondrial | 3.022 × 10^7^ | 0 | ∞ |
| Q3TKM5 | Idh3g | Isocitrate dehydrogenase [NAD] subunit, mitochondrial | 3.015 × 10^7^ | 0 | ∞ |
| Q5SYD0 | Myo1d | Unconventional myosin-Id | 2.776 × 10^7^ | 0 | ∞ |
| Q3UQ44 | Iqgap2 | Ras GTPase-activating-like protein IQGAP2 | 2.561 × 10^7^ | 0 | ∞ |
| Q9DCU9 | Hoga1 | 4-hydroxy-2-oxoglutarate aldolase, mitochondrial | 1.760 × 10^7^ | 0 | ∞ |
| Q3UFJ3 | Pdha1 | Pyruvate dehydrogenase E1 component subunit alpha | 1.496 × 10^7^ | 0 | ∞ |
| P17426-2 | Ap2a1 | Isoform B of AP-2 complex subunit alpha-1 | 1.156 × 10^7^ | 0 | ∞ |
| P17427 | Ap2a2 | AP-2 complex subunit alpha-2 | 1.156 × 10^7^ | 0 | ∞ |
| A0A0U1RPW2 | Tjp1 | Tight junction protein ZO-1 | 9.041 × 10^6^ | 0 | ∞ |
| Q3UGJ5 | Rasa3 | Uncharacterized protein | 8.876 × 10^6^ | 0 | ∞ |
| A0A0A0MQG2 | Sptbn1 | Spectrin beta chain, non-erythrocytic 1 (Fragment) | 6.806 × 10^6^ | 0 | ∞ |
| Q8CC88 | Vwa8 | von Willebrand factor A domain-containing protein 8 | 3.542 × 10^6^ | 0 | ∞ |
| A0A3B0IP04 | C1qa | Complement C1q subcomponent subunit A | 4.576 × 10^9^ | 1.772 × 10^7^ | 258.289 |
| P14106 | C1qb | Complement C1q subcomponent subunit B | 6.649 × 10^9^ | 3.353 × 10^7^ | 198.302 |

| Uniprot accession | Protein | Description | Abundance | | CPS23F:CPS8 fold |
| --- | --- | --- | --- | --- | --- |
|  |  |  | CPS23F | CPS8 |  |
| Q02105 | C1qc | Complement C1q subcomponent subunit C | 6.203 × 10^9^ | 3.497 × 10^7^ | 177.394 |
| E9Q6C2 | C1s1 | Complement component 1, s subcomponent 1 | 3.206 × 10^9^ | 2.660 × 10^7^ | 120.543 |
| G0YP42 | NA | Anti-human Langerin 2G3 lambda chain | 2.004 × 10^9^ | 6.616 × 10^7^ | 30.292 |
| Q9Z2T6 | Krt85 | Keratin, type II cuticular Hb5 | 5.756 × 10^10^ | 1.958 × 10^9^ | 29.404 |
| E9PVG8 | 9530053A07Rik | RIKEN cDNA 9530053A07 gene | 1.336 × 10^11^ | 5.573 × 10^9^ | 23.963 |
| P97290 | Serping1 | Plasma protease C1 inhibitor | 2.084 × 10^9^ | 9.106 × 10^7^ | 22.886 |
| P03995-2 | Gfap | Isoform 2 of Glial fibrillary acidic protein | 5.756 × 10^10^ | 2.749 × 10^9^ | 20.938 |
| Q61171 | Prdx2 | Peroxiredoxin-2 | 1.546 × 10^9^ | 1.028 × 10^8^ | 15.036 |
| Q3UAD6 | Hsp90b1 | HATPase_c domain-containing protein | 6.149 × 10^7^ | 4.119 × 10^6^ | 14.928 |
| Q0VBK2 | Krt80 | Keratin, type II cytoskeletal 80 | 2.116 × 10^8^ | 1.424 × 10^7^ | 14.860 |
| Q8BIG7 | Comtd1 | Catechol O-methyltransferase domain-containing protein 1 | 6.638 × 10^7^ | 5.160 × 10^6^ | 12.866 |
| A2AE15 | Cfp | Complement factor properdin | 1.705 × 10^9^ | 1.355 × 10^8^ | 12.580 |
| Q4L0E7 | Krt77 | Type II cytokeratin Kb39 (Fragment) | 6.887 × 10^10^ | 6.192 × 10^9^ | 11.122 |
| A0A0A0MQ76 | Nop58 | Nucleolar protein 58 | 8.422 × 10^7^ | 7.702 × 10^6^ | 10.934 |
| A0A087WQ46 | Nop58 | Nucleolar protein 58 (Fragment) | 8.422 × 10^7^ | 7.702 × 10^6^ | 10.934 |
| Q91X70 | C6 | Complement component 6 | 2.174 × 10^8^ | 2.044 × 10^7^ | 10.635 |
| Q61782 | NA | Type I epidermal keratin mRNA, 3'end (Fragment) | 7.079 × 10^9^ | 6.987 × 10^8^ | 10.132 |
| E9Q3W4 | Plec | Plectin | 1.679 × 10^8^ | 1.681 × 10^7^ | 9.985 |

| Uniprot accession | Protein | Description | Abundance | | CPS23F:CPS8 fold |
| --- | --- | --- | --- | --- | --- |
|  |  |  | CPS23F | CPS8 |  |
| A0JLV3 | Hist1h2bj | Histone H2B (Fragment) | 1.004 × 10^9^ | 1.033 × 10^8^ | 9.721 |
| P01027-2 | C3 | Isoform Short of Complement C3 | 4.169 × 10^10^ | 4.364 × 10^9^ | 9.554 |
| P42125 | Eci1 | Enoyl-CoA delta isomerase 1, mitochondrial | 2.033 × 10^8^ | 2.212 × 10^7^ | 9.187 |
| O35643 | Ap1b1 | AP-1 complex subunit beta-1 | 1.818 × 10^8^ | 2.000 × 10^7^ | 9.094 |
| Q8BH35 | C8b | Complement component C8 beta chain | 8.207 × 10^7^ | 9.724 × 10^6^ | 8.439 |
| Q3UBI6 | Rpl7 | Uncharacterized protein | 4.551 × 10^8^ | 5.630 × 10^7^ | 8.084 |
| E9Q1Z0 | Krt90 | Keratin 90 | 1.122 × 10^11^ | 1.408 × 10^10^ | 7.971 |
| Q60759 | Gcdh | Glutaryl-CoA dehydrogenase, mitochondrial | 1.316 × 10^8^ | 1.653 × 10^7^ | 7.962 |
| P97350 | Pkp1 | Plakophilin-1 | 5.830 × 10^8^ | 7.355 × 10^7^ | 7.927 |
| Q9CZJ2 | Hspa12b | Heat shock 70 kDa protein 12B | 3.900 × 10^7^ | 5.013 × 10^6^ | 7.779 |
| P04104 | Krt1 | Keratin, type II cytoskeletal 1 | 2.255 × 10^11^ | 2.973 × 10^10^ | 7.585 |
| P01632 | Igkv7-33 | Ig kappa chain V-I region S107A | 6.204 × 10^8^ | 8.380 × 10^7^ | 7.403 |
| P01027 | C3 | Complement C3 | 1.259 × 10^11^ | 1.703 × 10^10^ | 7.395 |
| Q3UAC2 | Rps3a1 | 40S ribosomal protein S3a | 7.314 × 10^7^ | 9.955 × 10^6^ | 7.347 |
| P50446 | Krt6a | Keratin, type II cytoskeletal 6A | 9.620 × 10^10^ | 1.315 × 10^10^ | 7.315 |
| Q3UV11 | Krt6b | Keratin, type II cytoskeletal 6B | 9.620 × 10^10^ | 1.315 × 10^10^ | 7.315 |
| Q8BGZ7 | Krt75 | Keratin, type II cytoskeletal 75 | 9.620 × 10^10^ | 1.315 × 10^10^ | 7.315 |
| Q32P04 | Krt5 | Keratin 5 | 9.620 × 10^10^ | 1.315 × 10^10^ | 7.315 |

| Uniprot accession | Protein | Description | Abundance | | CPS23F:CPS8 fold |
| --- | --- | --- | --- | --- | --- |
|  |  |  | CPS23F | CPS8 |  |
| E9Q557 | Dsp | Desmoplakin | 2.236 × 10^9^ | 3.075 × 10^8^ | 7.273 |
| Q9DCV7 | Krt7 | Keratin, type II cytoskeletal 7 | 3.380 × 10^10^ | 4.701 × 10^9^ | 7.191 |
| A0A2R8VHP3 | Gm5478 | Predicted pseudogene 5478 | 1.043 × 10^11^ | 1.458 × 10^10^ | 7.156 |
| Q6NXH9 | Krt73 | Keratin, type II cytoskeletal 73 | 1.687 × 10^11^ | 2.412 × 10^10^ | 6.993 |
| Q08EK4 | Krt77 | Keratin 77 | 1.954 × 10^11^ | 2.837 × 10^10^ | 6.887 |
| Q5R3T5 | G8(anti-MRBC) | G8(Anti-MRBC hybridoma) light chain (Fragment) | 2.141 × 10^8^ | 3.127 × 10^7^ | 6.846 |
| Q9R0H5 | Krt71 | Keratin, type II cytoskeletal 71 | 1.526 × 10^11^ | 2.269 × 10^10^ | 6.727 |
| Q6IFZ9 | Krt74 | Keratin, type II cytoskeletal 74 | 1.526 × 10^11^ | 2.269 × 10^10^ | 6.727 |
| Q9DCW4 | Etfb | Electron transfer flavoprotein subunit beta | 1.496 × 10^8^ | 2.248 × 10^7^ | 6.654 |
| A0A0B4J1N0 | Ighv1-76 | Immunoglobulin heavy variable 1-76 | 8.139 × 10^8^ | 1.227 × 10^8^ | 6.632 |
| H3BL60 | C3 | Complement C3 (Fragment) | 4.223 × 10^10^ | 6.812 × 10^9^ | 6.199 |
| Q61176 | Arg1 | Arginase-1 | 4.632 × 10^8^ | 7.595 × 10^7^ | 6.099 |
| Q9D9P1 | Chchd3 | MICOS complex subunit Mic19 | 2.521 × 10^7^ | 4.200 × 10^6^ | 6.003 |
| Q99LC5 | Etfa | Electron transfer flavoprotein subunit alpha, mitochondrial | 1.572 × 10^8^ | 2.621 × 10^7^ | 5.999 |
| Q02257 | Jup | Junction plakoglobin | 1.809 × 10^9^ | 3.017 × 10^8^ | 5.996 |
| A1L0X5 | Krt78 | Krt78 protein (Fragment) | 9.494 × 10^10^ | 1.672 × 10^10^ | 5.680 |
| Q8BP54 | Pcx | Uncharacterized protein (Fragment) | 3.810 × 10^7^ | 6.941 × 10^6^ | 5.490 |
| A2A998 | C8a | Complement component C8 alpha chain | 6.800 × 10^7^ | 1.241 × 10^7^ | 5.480 |

| Uniprot accession | Protein | Description | Abundance | | CPS23F:CPS8 fold |
| --- | --- | --- | --- | --- | --- |
|  |  |  | CPS23F | CPS8 |  |
| Q3U026 | Mogs | Uncharacterized protein | 1.214 × 10^8^ | 2.225 × 10^7^ | 5.458 |
| Q5SUA5 | Myo1g | Unconventional myosin-Ig | 8.922 × 10^7^ | 1.760 × 10^7^ | 5.071 |
| P11679 | Krt8 | Keratin, type II cytoskeletal 8 | 1.475 × 10^10^ | 2.946 × 10^9^ | 5.005 |
| Q4VA93 | Prkca | Protein kinase C | 6.742 × 10^7^ | 1.356 × 10^7^ | 4.972 |
| Q99M73 | Krt84 | Keratin, type II cuticular Hb4 | 1.484 × 10^10^ | 3.034 × 10^9^ | 4.891 |
| B9EIU2 | C4a | C4a anaphylatoxin | 7.359 × 10^8^ | 1.506 × 10^8^ | 4.886 |
| Q6IFZ8 | Gm5414 | Predicted gene 5414 | 1.269 × 10^10^ | 2.700 × 10^9^ | 4.700 |
| O70152 | Dpm1 | Dolichol-phosphate mannosyltransferase subunit 1 | 6.392 × 10^7^ | 1.421 × 10^7^ | 4.498 |
| A0A286YDB7 | Ssr1 | Signal sequence receptor subunit alpha (Fragment) | 1.492 × 10^8^ | 3.362 × 10^7^ | 4.437 |
| Q9EQK5 | Mvp | Major vault protein | 8.311 × 10^7^ | 1.886 × 10^7^ | 4.406 |
| Q8VED5 | Krt79 | Keratin, type II cytoskeletal 79 | 3.393 × 10^10^ | 7.781 × 10^9^ | 4.361 |
| P07744 | Krt4 | Keratin, type II cytoskeletal 4 | 1.632 × 10^10^ | 3.771 × 10^9^ | 4.328 |
| P20918 | Plg | Plasminogen | 5.140 × 10^8^ | 1.192 × 10^8^ | 4.313 |
| P06909 | Cfh | Complement factor H | 1.450 × 10^9^ | 3.369 × 10^8^ | 4.305 |
| Q8C196 | Cps1 | Carbamoyl-phosphate synthase [ammonia], mitochondrial | 1.486 × 10^9^ | 3.479 × 10^8^ | 4.270 |
| Q3TQD9 | Sardh | Uncharacterized protein (Fragment) | 8.950 × 10^7^ | 2.111 × 10^7^ | 4.240 |
| E9Q8B5 | Cfhr4 | Complement factor H-related 4 | 5.454 × 10^8^ | 1.303 × 10^8^ | 4.185 |
| A4FUS1 | Rps16 | Rps16 protein | 1.857 × 10^8^ | 4.502 × 10^7^ | 4.126 |

| Uniprot accession | Protein | Description | Abundance | | CPS23F:CPS8 fold |
| --- | --- | --- | --- | --- | --- |
|  |  |  | CPS23F | CPS8 |  |
| A0A3B2WBL1 | Rpl10a | Ribosomal protein | 1.184 × 10^9^ | 2.949 × 10^8^ | 4.015 |
| Q9CR57 | Rpl14 | 60S ribosomal protein L14 | 1.852 × 10^8^ | 4.647 × 10^7^ | 3.985 |
| Q03265 | Atp5f1a | ATP synthase subunit alpha, mitochondrial | 1.111 × 10^9^ | 2.814 × 10^8^ | 3.949 |
| P01029 | C4b | Complement C4-B | 8.334 × 10^8^ | 2.111 × 10^8^ | 3.948 |
| Q14B21 | Mrpl9 | 39S ribosomal protein L9, mitochondrial | 5.759 × 10^7^ | 1.470 × 10^7^ | 3.917 |
| A2AMW0 | Capzb | F-actin-capping protein subunit beta | 1.374 × 10^8^ | 3.527 × 10^7^ | 3.895 |
| Q3TR93 | Decr2 | Uncharacterized protein | 4.438 × 10^7^ | 1.158 × 10^7^ | 3.832 |
| P14115 | Rpl27a | 60S ribosomal protein L27a | 1.710 × 10^8^ | 4.513 × 10^7^ | 3.790 |
| Q61495 | Dsg1a | Desmoglein-1-alpha | 4.736 × 10^8^ | 1.252 × 10^8^ | 3.782 |
| P35700 | Prdx1 | Peroxiredoxin-1 | 3.370 × 10^8^ | 8.911 × 10^7^ | 3.782 |
| Q8R0I8 | Cfhr2 | BC026782 protein | 1.080 × 10^9^ | 2.858 × 10^8^ | 3.777 |
| Q9DBG1 | Cyp27a1 | Sterol 26-hydroxylase, mitochondrial | 2.111 × 10^8^ | 5.618 × 10^7^ | 3.757 |
| Q3U861 | Atp6v1d | V-type proton ATPase subunit D | 2.851 × 10^8^ | 7.666 × 10^7^ | 3.719 |
| P62830 | Rpl23 | 60S ribosomal protein L23 | 1.217 × 10^8^ | 3.293 × 10^7^ | 3.697 |
| F8VPR2 | Fmnl2 | Formin-like protein 2 | 2.802 × 10^7^ | 7.593 × 10^6^ | 3.690 |
| P45952 | Acadm | Medium-chain specific acyl-CoA dehydrogenase, mitochondrial | 9.905 × 10^7^ | 2.709 × 10^7^ | 3.657 |
| Q3TKR5 | Rpl5 | Ribosomal protein L5 | 1.696 × 10^8^ | 4.662 × 10^7^ | 3.638 |
| P68404 | Prkcb | Protein kinase C beta type | 3.678 × 10^7^ | 1.018 × 10^7^ | 3.615 |

| Uniprot accession | Protein | Description | Abundance | | CPS23F:CPS8 fold |
| --- | --- | --- | --- | --- | --- |
|  |  |  | CPS23F | CPS8 |  |
| P33267 | Cyp2f2 | Cytochrome P450 2F2 | 7.924 × 10^7^ | 2.196 × 10^7^ | 3.609 |
| Q00PI9 | Hnrnpul2 | Heterogeneous nuclear ribonucleoprotein U-like protein 2 | 1.743 × 10^8^ | 4.848 × 10^7^ | 3.595 |
| Q5SVF7 | Nipsnap1 | Uncharacterized protein | 1.508 × 10^10^ | 4.250 × 10^9^ | 3.548 |
| Q5EBQ6 | Rpl9 | 60S ribosomal protein L9 | 8.030 × 10^8^ | 2.282 × 10^8^ | 3.519 |
| Q53ZD4 | Mgst1 | Glutathione transferase | 6.168 × 10^8^ | 1.763 × 10^8^ | 3.500 |
| O89054 | Actb | Cytoskeletal beta-actin (Fragment) | 9.044 × 10^8^ | 2.587 × 10^8^ | 3.496 |
| Q8BH80 | Vapb | Vesicle-associated membrane protein, associated protein B and C | 8.400 × 10^7^ | 2.421 × 10^7^ | 3.469 |
| Q9CPQ8 | Atp5mg | ATP synthase subunit g, mitochondrial | 1.978 × 10^9^ | 5.704 × 10^8^ | 3.468 |
| Q99M74 | Krt82 | Keratin, type II cuticular Hb2 | 1.470 × 10^10^ | 4.312 × 10^9^ | 3.409 |
| Q3TD78 | Nipsnap2 | NIPSNAP domain-containing protein | 3.235 × 10^8^ | 9.673 × 10^7^ | 3.344 |
| Q9WV55 | Vapa | Vesicle-associated membrane protein-associated protein A | 1.077 × 10^8^ | 3.244 × 10^7^ | 3.321 |
| P97429 | Anxa4 | Annexin A4 | 2.395 × 10^8^ | 7.443 × 10^7^ | 3.217 |
| Q545A2 | Slc25a5 | ADP/ATP translocase | 1.858 × 10^9^ | 5.871 × 10^8^ | 3.164 |
| Q91XB2 | Mrpl3 | Mitochondrial ribosomal protein L3 | 1.118 × 10^8^ | 3.564 × 10^7^ | 3.136 |
| Q8JZU2 | Slc25a1 | Tricarboxylate transport protein, mitochondrial | 1.205 × 10^8^ | 3.843 × 10^7^ | 3.136 |
| Q99LC3 | Ndufa10 | NADH dehydrogenase [ubiquinone] 1 alpha subcomplex subunit 10, mitochondrial | 4.595 × 10^7^ | 1.466 × 10^7^ | 3.135 |

| Uniprot accession | Protein | Description | Abundance | | CPS23F:CPS8 fold |
| --- | --- | --- | --- | --- | --- |
|  |  |  | CPS23F | CPS8 |  |
| Q8C166 | Cpne1 | Copine-1 | 7.679 × 10^8^ | 2.451 × 10^8^ | 3.133 |
| A0A668KLU9 | Cfhr2 | Complement factor H-related 2 | 9.925 × 10^8^ | 3.177 × 10^8^ | 3.124 |
| P54869 | Hmgcs2 | Hydroxymethylglutaryl-CoA synthase, mitochondrial | 1.129 × 10^8^ | 3.620 × 10^7^ | 3.118 |
| Q8CHT0 | Aldh4a1 | Delta-1-pyrroline-5-carboxylate dehydrogenase, mitochondrial | 3.776 × 10^7^ | 1.215 × 10^7^ | 3.108 |
| Q8BH95 | Echs1 | Enoyl-CoA hydratase, mitochondrial | 4.340 × 10^7^ | 1.397 × 10^7^ | 3.107 |
| Q3U6S1 | Vim | Vimentin | 2.779 × 10^8^ | 8.996 × 10^7^ | 3.089 |
| Q5I0W0 | Atp5pb | ATP synthase F(0) complex subunit B1, mitochondrial | 2.176 × 10^8^ | 7.054 × 10^7^ | 3.085 |
| Q9R092 | Hsd17b6 | 17-beta-hydroxysteroid dehydrogenase type 6 | 5.621 × 10^7^ | 1.851 × 10^7^ | 3.037 |
| O88451 | Rdh7 | Retinol dehydrogenase 7 | 9.450 × 10^7^ | 3.118 × 10^7^ | 3.031 |
| A0A0R3P9C8 | Ndufa9 | NADH dehydrogenase 1 alpha subcomplex subunit 9, mitochondrial | 3.467 × 10^8^ | 1.147 × 10^8^ | 3.024 |
| Q4FJU3 | Crip2 | Crip2 protein | 2.104 × 10^8^ | 6.971 × 10^7^ | 3.019 |
| Q3UGS1 | Nipsnap1 | NIPSNAP domain-containing protein | 7.062 × 10^9^ | 2.360 × 10^9^ | 2.992 |
| Q5J7N1 | Kras | Kras protein | 1.479 × 10^8^ | 4.963 × 10^7^ | 2.979 |
| Q0VDV7 | Kras | Kras protein | 1.479 × 10^8^ | 4.963 × 10^7^ | 2.979 |
| Q3THU8 | Slc25a3 | Uncharacterized protein | 2.713 × 10^8^ | 9.160 × 10^7^ | 2.962 |
| Q80XN0 | Bdh1 | D-beta-hydroxybutyrate dehydrogenase, mitochondrial | 2.267 × 10^9^ | 7.723 × 10^8^ | 2.936 |
| O88844 | Idh1 | Isocitrate dehydrogenase [NADP] cytoplasmic | 5.674 × 10^7^ | 1.936 × 10^7^ | 2.932 |

| Uniprot accession | Protein | Description | Abundance | | CPS23F:CPS8 fold |
| --- | --- | --- | --- | --- | --- |
|  |  |  | CPS23F | CPS8 |  |
| Q9R0X4 | Acot9 | Acyl-coenzyme A thioesterase 9, mitochondrial | 7.331 × 10^7^ | 2.514 × 10^7^ | 2.915 |
| A2A513 | Krt10 | Keratin, type I cytoskeletal 10 | 7.987 × 10^10^ | 2.746 × 10^10^ | 2.909 |
| Q3UA17 | Mtch2 | Uncharacterized protein | 1.390 × 10^8^ | 4.801 × 10^7^ | 2.896 |
| P67778 | Phb | Prohibitin | 3.483 × 10^8^ | 1.203 × 10^8^ | 2.895 |
| P51175 | Ppox | Protoporphyrinogen oxidase | 1.452 × 10^8^ | 5.026 × 10^7^ | 2.890 |
| P21956-2 | Mfge8 | Isoform 2 of Lactadherin | 9.647 × 10^7^ | 3.364 × 10^7^ | 2.868 |
| A0A075B5M7 | Igkv5-39 | Immunoglobulin kappa variable 5-39 | 1.144 × 10^8^ | 4.033 × 10^7^ | 2.837 |
| Q9QZD8 | Slc25a10 | Mitochondrial dicarboxylate carrier | 1.062 × 10^8^ | 3.747 × 10^7^ | 2.833 |
| P99028 | Uqcrh | Cytochrome b-c1 complex subunit 6, mitochondrial | 3.466 × 10^8^ | 1.230 × 10^8^ | 2.817 |
| Q3U9P0 | Rps10 | S10_plectin domain-containing protein | 2.166 × 10^8^ | 7.690 × 10^7^ | 2.816 |
| Q5M9L7 | Rps17 | 40S ribosomal protein S17 | 6.270 × 10^8^ | 2.257 × 10^8^ | 2.778 |
| Q9D0M3-2 | Cyc1 | Isoform 2 of Cytochrome c1, heme protein, mitochondrial | 7.092 × 10^9^ | 2.562 × 10^9^ | 2.768 |
| Q9ESW4 | Agk | Acylglycerol kinase, mitochondrial | 1.811 × 10^8^ | 6.576 × 10^7^ | 2.754 |
| Q3TS44 | Psma1 | Proteasome subunit alpha type | 6.687 × 10^7^ | 2.432 × 10^7^ | 2.750 |
| Q8C2Q8 | Atp5c1 | ATP synthase subunit gamma | 1.649 × 10^10^ | 6.002 × 10^9^ | 2.748 |
| A2AKU9 | Atp5c1 | ATP synthase subunit gamma | 1.649 × 10^10^ | 6.002 × 10^9^ | 2.748 |
| Q76LB8 | Rdh9 | Cis-retinol/androgen dehydrogenase type 3 | 5.394 × 10^7^ | 1.965 × 10^7^ | 2.745 |
| A1E2B8 | NA | Inducible heat shock protein 70 | 2.428 × 10^8^ | 8.915 × 10^7^ | 2.723 |

| Uniprot accession | Protein | Description | Abundance | | CPS23F:CPS8 fold |
| --- | --- | --- | --- | --- | --- |
|  |  |  | CPS23F | CPS8 |  |
| P16054 | Prkce | Protein kinase C epsilon type | 1.291 × 10^8^ | 4.779 × 10^7^ | 2.702 |
| P06684 | C5 | Complement C5 | 2.392 × 10^8^ | 8.867 × 10^7^ | 2.698 |
| Q9DCT2 | Ndufs3 | NADH dehydrogenase iron-sulfur protein 3, mitochondrial | 3.823 × 10^7^ | 1.426 × 10^7^ | 2.681 |
| Q06185 | Atp5me | ATP synthase subunit e, mitochondrial | 5.504 × 10^8^ | 2.055 × 10^8^ | 2.678 |
| P35550 | Fbl | rRNA 2'-O-methyltransferase fibrillarin | 2.567 × 10^7^ | 9.684 × 10^6^ | 2.650 |
| Q8BH04 | Pck2 | Phosphoenolpyruvate carboxykinase [GTP], mitochondrial | 2.699 × 10^7^ | 1.022 × 10^7^ | 2.642 |
| Q3V340 | Adap2 | Uncharacterized protein | 5.671 × 10^7^ | 2.160 × 10^7^ | 2.626 |
| Q545F8 | Rps4x | 40S ribosomal protein S4 | 5.649 × 10^8^ | 2.152 × 10^8^ | 2.625 |
| Z4YKT6 | Dhrs7b | Dehydrogenase/reductase SDR family member 7B | 4.303 × 10^7^ | 1.642 × 10^7^ | 2.620 |
| Q5FWI9 | Ap2m1 | AP-2 complex subunit mu | 1.347 × 10^8^ | 5.164 × 10^7^ | 2.608 |
| Q3TWV4 | Ap2m1 | AP-2 complex subunit mu | 1.347 × 10^8^ | 5.164 × 10^7^ | 2.608 |
| Q9DB77 | Uqcrc2 | Cytochrome b-c1 complex subunit 2, mitochondrial | 2.965 × 10^8^ | 1.142 × 10^8^ | 2.596 |
| Q3TIQ2 | Rpl12 | 60S ribosomal protein L12 | 2.400 × 10^8^ | 9.251 × 10^7^ | 2.595 |
| Q3UJ34 | Ass1 | Argininosuccinate synthase | 8.799 × 10^7^ | 3.394 × 10^7^ | 2.593 |
| G5E8R3 | Pcx | Pyruvate carboxylase | 4.843 × 10^7^ | 1.906 × 10^7^ | 2.541 |
| Q3TCQ3 | Pcx | Pyruvate carboxylase | 4.843 × 10^7^ | 1.906 × 10^7^ | 2.541 |
| Q80X85 | Mrps7 | 28S ribosomal protein S7, mitochondrial | 2.671 × 10^8^ | 1.054 × 10^8^ | 2.534 |

| Uniprot accession | Protein | Description | Abundance | | CPS23F:CPS8 fold |
| --- | --- | --- | --- | --- | --- |
|  |  |  | CPS23F | CPS8 |  |
| B1ARA3 | Rpl26 | 60S ribosomal protein L26 (Fragment) | 1.186 × 10^8^ | 4.697 × 10^7^ | 2.525 |
| A0A0R4IZZ5 | Clec4f | C-type lectin domain family 4 member F | 7.238 × 10^7^ | 2.867 × 10^7^ | 2.524 |
| A8DUP7 | Hbbt1 | Beta-globin | 3.803 × 10^9^ | 1.527 × 10^9^ | 2.491 |
| P48962 | Slc25a4 | ADP/ATP translocase 1 | 1.457 × 10^9^ | 5.871 × 10^8^ | 2.481 |
| P49718 | Mcm5 | DNA replication licensing factor MCM5 | 3.651 × 10^7^ | 1.473 × 10^7^ | 2.479 |
| P20444 | Prkca | Protein kinase C alpha type | 3.349 × 10^7^ | 1.356 × 10^7^ | 2.470 |
| Q542G9 | Anxa2 | Annexin | 3.538 × 10^8^ | 1.441 × 10^8^ | 2.455 |
| Q8VCL2 | Sco2 | Protein SCO2 homolog, mitochondrial | 2.508 × 10^8^ | 1.031 × 10^8^ | 2.433 |
| Q91X77 | Cyp2c50 | Cytochrome P450 2C50 | 2.202 × 10^8^ | 9.057 × 10^7^ | 2.432 |
| P56654 | Cyp2c37 | Cytochrome P450 2C37 | 2.202 × 10^8^ | 9.057 × 10^7^ | 2.432 |
| Q6XVG2 | Cyp2c54 | Cytochrome P450 2C54 | 2.202 × 10^8^ | 9.057 × 10^7^ | 2.432 |
| Q99JR1 | Sfxn1 | Sideroflexin-1 | 5.253 × 10^7^ | 2.162 × 10^7^ | 2.430 |
| Q3U0S6 | Rasip1 | Ras-interacting protein 1 | 5.526 × 10^8^ | 2.278 × 10^8^ | 2.426 |
| P54071 | Idh2 | Isocitrate dehydrogenase [NADP], mitochondrial | 7.282 × 10^8^ | 3.014 × 10^8^ | 2.416 |
| P47963 | Rpl13 | 60S ribosomal protein L13 | 2.375 × 10^8^ | 9.841 × 10^7^ | 2.414 |
| A0A0B4J1J2 | Igkv5-43 | Immunoglobulin kappa chain variable 5-43 (Fragment) | 1.603 × 10^8^ | 6.665 × 10^7^ | 2.405 |
| P01642 | Gm10881 | Ig kappa chain V-V region L7 (Fragment) | 1.603 × 10^8^ | 6.665 × 10^7^ | 2.405 |
| Q5SX53 | Slc25a11 | Uncharacterized protein | 1.890 × 10^8^ | 7.904 × 10^7^ | 2.391 |

| Uniprot accession | Protein | Description | Abundance | | CPS23F:CPS8 fold |
| --- | --- | --- | --- | --- | --- |
|  |  |  | CPS23F | CPS8 |  |
| Q3UKH3 | Acaa2 | Uncharacterized protein | 1.147 × 10^9^ | 4.818 × 10^8^ | 2.380 |
| Q99JY0 | Hadhb | Trifunctional enzyme subunit beta, mitochondrial | 1.414 × 10^9^ | 5.961 × 10^8^ | 2.372 |
| P11928 | Oas1a | 2'-5'-oligoadenylate synthase 1A | 8.029 × 10^7^ | 3.404 × 10^7^ | 2.359 |
| P28843 | Dpp4 | Dipeptidyl peptidase 4 | 2.092 × 10^8^ | 8.911 × 10^7^ | 2.348 |
| P43024 | Cox6a1 | Cytochrome c oxidase subunit 6A1, mitochondrial | 2.011 × 10^9^ | 8.584 × 10^8^ | 2.343 |
| Q9D023 | Mpc2 | Mitochondrial pyruvate carrier 2 | 1.475 × 10^8^ | 6.353 × 10^7^ | 2.322 |
| Q99N15 | Hsd17b10 | 17beta-hydroxysteroid dehydrogenase type 10/short chain L-3-hydroxyacyl-CoA dehydrogenas | 7.508 × 10^7^ | 3.240 × 10^7^ | 2.318 |
| Q54AH9 | Hbb-b2 | Beta-2-globin (Fragment) | 2.972 × 10^9^ | 1.292 × 10^9^ | 2.300 |
| A8DUM2 | Hbbt1 | Beta-globin | 3.803 × 10^9^ | 1.659 × 10^9^ | 2.292 |
| Q9ERI6 | Rdh14 | Retinol dehydrogenase 14 | 6.549 × 10^7^ | 2.892 × 10^7^ | 2.264 |
| Q9DBM2 | Ehhadh | Peroxisomal bifunctional enzyme | 3.280 × 10^8^ | 1.458 × 10^8^ | 2.249 |
| Q8VEH7 | Oas1g | 2'-5' oligoadenylate synthase | 7.654 × 10^7^ | 3.404 × 10^7^ | 2.249 |
| B1Q450 | HBB1 | Hemoglobin beta chain subunit | 4.079 × 10^9^ | 1.819 × 10^9^ | 2.242 |
| J3QNY6 | Abcb11 | Bile salt export pump | 5.077 × 10^7^ | 2.265 × 10^7^ | 2.241 |
| Q9DB20 | Atp5po | ATP synthase subunit O, mitochondrial | 1.995 × 10^9^ | 8.969 × 10^8^ | 2.225 |
| D3YUT3 | Rps19 | 40S ribosomal protein S19 (Fragment) | 4.018 × 10^8^ | 1.807 × 10^8^ | 2.224 |
| Q549A5 | Clu | Clusterin | 3.740 × 10^9^ | 1.685 × 10^9^ | 2.219 |
| Q3TVJ8 | Ssr4 | Signal sequence receptor subunit delta | 9.117 × 10^7^ | 4.145 × 10^7^ | 2.200 |

| Uniprot accession | Protein | Description | Abundance | | CPS23F:CPS8 fold |
| --- | --- | --- | --- | --- | --- |
|  |  |  | CPS23F | CPS8 |  |
| Q2XU92 | Acsbg2 | Long-chain-fatty-acid--CoA ligase ACSBG2 | 2.468 × 10^9^ | 1.123 × 10^9^ | 2.198 |
| P56480 | Atp5f1b | ATP synthase subunit beta, mitochondrial | 1.943 × 10^8^ | 8.902 × 10^7^ | 2.183 |
| Q8VDD5 | Myh9 | Myosin-9 | 1.011 × 10^8^ | 4.639 × 10^7^ | 2.179 |
| Q9Z0X1 | Aifm1 | Apoptosis-inducing factor 1, mitochondrial | 3.229 × 10^7^ | 1.489 × 10^7^ | 2.169 |
| Q3V1K9 | Des | IF rod domain-containing protein | 1.083 × 10^8^ | 5.020 × 10^7^ | 2.157 |
| Q3TTN3 | Vdac3 | Voltage-dependent anion-selective channel protein 3 | 2.530 × 10^8^ | 1.180 × 10^8^ | 2.145 |
| Q9QUK9 | Try5 | TESP4 | 7.629 × 10^10^ | 3.594 × 10^10^ | 2.123 |
| E9Q986 | Ctnnd1 | Catenin delta-1 | 1.669 × 10^8^ | 7.952 × 10^7^ | 2.099 |
| Q8CB17 | Fetub | Uncharacterized protein | 1.365 × 10^8^ | 6.507 × 10^7^ | 2.098 |
| A8DUK2 | Hbbt1 | Beta-globin | 5.656 × 10^9^ | 2.700 × 10^9^ | 2.095 |
| D3YUP5 | Exoc3l2 | Exocyst complex component 3-like 2 | 1.210 × 10^8^ | 5.804 × 10^7^ | 2.084 |
| Q3UEK1 | Mbl2 | Mannan-binding protein | 1.947 × 10^9^ | 9.353 × 10^8^ | 2.082 |
| A0A068BGR9 | Ndufa7 | Complex I-B14.5a | 1.158 × 10^8^ | 5.571 × 10^7^ | 2.078 |
| Q8BMD8 | Slc25a24 | Calcium-binding mitochondrial carrier protein SCaMC-1 | 2.364 × 10^8^ | 1.139 × 10^8^ | 2.075 |
| A0A1L1SQA8 | Rps25 | 40S ribosomal protein S25 | 4.448 × 10^8^ | 2.144 × 10^8^ | 2.074 |
| Q8BU88 | Mrpl22 | 39S ribosomal protein L22, mitochondrial | 1.531 × 10^8^ | 7.389 × 10^7^ | 2.073 |
| Q921R2 | Rps13 | 40S ribosomal protein S13 | 1.092 × 10^9^ | 5.272 × 10^8^ | 2.071 |
| Q9R0Z4 | Ehhadh | L-specific multifunctional beta-oxdiation protein | 1.791 × 10^8^ | 8.659 × 10^7^ | 2.069 |

| Uniprot accession | Protein | Description | Abundance | | CPS23F:CPS8 fold |
| --- | --- | --- | --- | --- | --- |
|  |  |  | CPS23F | CPS8 |  |
| P21614 | Gc | Vitamin D-binding protein | 3.846 × 10^8^ | 1.865 × 10^8^ | 2.062 |
| Q9D8L4 | NA | Uncharacterized protein | 3.695 × 10^8^ | 1.793 × 10^8^ | 2.061 |
| A0A075B5N3 | Igkv8-28 | Immunoglobulin kappa variable 8-28 | 1.757 × 10^9^ | 8.629 × 10^8^ | 2.036 |
| Q9WTI7 | Myo1c | Unconventional myosin-Ic | 1.605 × 10^9^ | 7.945 × 10^8^ | 2.020 |
| A0A4V6JA63 | Ighg2b | IgG2b (Fragment) | 4.208 × 10^8^ | 2.083 × 10^8^ | 2.020 |
| Q7M754 | Gm5409 | Try10-like trypsinogen | 1.085 × 10^9^ | 5.378 × 10^8^ | 2.018 |
| Q91VT4 | Cbr4 | Carbonyl reductase family member 4 | 7.887 × 10^8^ | 3.913 × 10^8^ | 2.016 |
| P62843 | Rps15 | 40S ribosomal protein S15 | 1.353 × 10^8^ | 6.757 × 10^7^ | 2.003 |
| P54310 | Lipe | Hormone-sensitive lipase | 7.734 × 10^7^ | 3.867 × 10^7^ | 2.000 |

*^a^*Proteins were considered as CPS23F receptor candidates once meeting following criteria:

1. Annotated as receptor or binding proteins;

2. CPS23F:CPS8 enrichment fold ≥ 2.000.

*^b^*Proteins were ranked according to the protein abundance in CPS23F group.

NA, not accessible.
