## Supplemental Table S3 for "C-Reactive Protein Drives Potent Clearance of Blood Bacteria in the Liver by Activating the Complement System"

**Table S3. Information of strains with recombinant protein plasmids**

| Strain ID | Plasmid backbone | Insertion sequence/segment | Application |
| --- | --- | --- | --- |
| TH17157 | pCMV-chikv-strepII | cDNA of mouse CRP | Overexpression of candidates in  HEK293F cells |
| TH17158 | pCMV-chikv-strepII | cDNA of human CRP |  |
| TH17198 | pCMV-chikv-strepII | cDNA of E81A mutant mouse CRP |  |
| TH17199 | pCMV-chikv-strepII | cDNA of F66A mutant mouse CRP |  |
| TH17200 | pCMV-chikv-strepII | cDNA of E81A mutant human CRP |  |
| TH17201 | pCMV-chikv-strepII | cDNA of F66A mutant human CRP |  |
| TH17202 | pCMV-chikv-strepII | cDNA of T76Y mutant human CRP |  |
| TH17202 | pCMV-chikv-strepII | cDNA of G76Y mutant mouse CRP |  |
| TH17388 | pCMV-chikv-strepII | cDNA of 71-91 mutant human CRP |  |
