## Supplemental Table S4 for "C-Reactive Protein Drives Potent Clearance of Blood Bacteria in the Liver by Activating the Complement System"

**Table S4.** **Primers used in this study**

| Primer ID | Sequence (5’-3’)*^a^* | Application |
| --- | --- | --- |
| Pr19504 | GTCGACATGGAGAAGCTACTCTGGTGCCTTC | Amplification of mouse CRP fragment in TH17157 |
| Pr19505 | GGATCCTCATTTTTCGAACTGCGGGTGGCTCCAACCTCCCGATCCACCTCCGGACCACAGCTGCGGCTTAATAAAC |  |
| Pr19512 | AAAACTGCAGGCCACCATGGAGAAGCTGT | Amplification of human CRP fragment in TH17158 |
| Pr19513 | CGCGGATCCTCATTTTTCGAACTG |  |
| Pr19562 | GAAT**CGT**ACTGCAGCACCACCCACTCCAAAAGTATACTGTTTATCCTTATTCCAAAATATGAGAAT | Construction of E81A mutant mouse CRP with  Pr19504/Pr19505 from TH17157 |
| Pr19563 | ATTCTCATATTTTGGAATAAGGATAAACAGTATACTTTTGGAGTGGGTGGTGCT**GCA**GTACGATTC |  |
| Pr19564 | GAATCGTACTTCAGCACCACCCACTCCAAAAGTATACTGTTTATCCTTATTCCA**AGC**TATGAGAAT | Construction of F66A mutant mouse CRP with Pr19504/Pr19505 from TH17157 |
| Pr19565 | ATTCTCATA**GCT**TGGAATAAGGATAAACAGTATACTTTTGGAGTGGGTGGTGCTGAAGTACGATTC |  |
| Pr19566 | GAATAATAT**TGC**AGACCCACCCACTGTAAAACTGTATCCTATATCCTTAGACCAAAATATGAGAAT | Construction of E81A mutant human CRP with Pr19512/Pr19513 from TH17158 |
| Pr19567 | ATTCTCATATTTTGGTCTAAGGATATAGGATACAGTTTTACAGTGGGTGGGTCT**GCA**ATATTATTC |  |
| Pr19568 | GAATAATATTTCAGACCCACCCACTGTAAAACTGTATCCTATATCCTTAGACCAA**GCT**ATGAGAAT | Construction of F66A mutant human CRP with Pr19512/Pr19513 from TH17158 |
| Pr19569 | ATTCTCAT**AGC**TTGGTCTAAGGATATAGGATACAGTTTTACAGTGGGTGGGTCTGAAATATTATTC |  |
| Pr19570 | GAATAATATTTCAGACCCACCCACGTAAAAACT**GTA**TCCTATATCCTTAGACCAAAATATGAGAAT | Construction of T76Y mutant human CRP with Pr19512/Pr19513 from TH17158 |
| Pr19571 | ATTCTCATATTTTGGTCTAAGGATATAGGA**TAC**AGTTTTTACGTGGGTGGGTCTGAAATATTATTC |  |
| Pr19572 | GAATCGTACTTCAGCACCACCCACGTAAAAA**GTA**TACTGTTTATCCTTATTCCAAAATATGAGAAT | Construction of G76Y mutant mouse CRP with  Pr19504/Pr19505 from TH17157 |
| Pr19573 | ATTCTCATATTTTGGAATAAGGATAAACAGTA**TAC**TTTTTACGTGGGTGGTGCTGAAGTACGATTC |  |
| Pr19933 | **GGTGCTGAAGTACGATTCATGGTTTCAGAGATTCCTGAG**GCTCCAGTACACATTTGTAC | Amplification of human CRP^m71-91^ in TH17158 |
| Pr19934 | **TCGTACTTCAGCACCACCCACTCCAAAAGTATACTGTTT**ATCCTTAGACCAAAATATGA |  |

*^a^* Underlined nucleotides indicate restriction enzyme sites, and bold nucleotides suggest mutated sites.
