## Supplemental Table S5 for "C-Reactive Protein Drives Potent Clearance of Blood Bacteria in the Liver by Activating the Complement System"

**Table S5. CPS23F-binding human protein candidates identified by mass spectrometry*^a^***

| Uniprot accession | Protein | Description | Abundance | | CPS23F:CPS8 fold |
| --- | --- | --- | --- | --- | --- |
|  |  |  | CPS23F*^b^* | CPS8 |  |
| P02776 | PF4 | Platelet factor 4 | 1.211 × 10^9^ | 0 | ∞ |
| P02741 | Crp | C-reactive protein | 8.833 × 10^8^ | 0 | ∞ |
| P12883 | **MYH7** | Myosin-7 | 7.135 × 10^8^ | 0 | ∞ |
| P13533 | **MYH6** | Myosin-6 | 6.685 × 10^8^ | 0 | ∞ |
| A0A6E1W127 | **MYH7B** | Myosin-7B | 4.116 × 10^8^ | 0 | ∞ |
| A0A590UJU8 | **MYL2** | Myosin regulatory light chain 2, ventricular/cardiac muscle isoform | 3.322 × 10^8^ | 0 | ∞ |
| I3L1K6 | **MYL4** | Myosin light chain 4 | 3.192 × 10^8^ | 0 | ∞ |
| P12524 | **MYCL** | Protein L-Myc | 2.374 × 10^8^ | 0 | ∞ |
| Q9UIW2 | **PLXNA1** | Plexin-A1 | 2.022 × 10^8^ | 0 | ∞ |
| K7EQL4 | **TNNT1** | Troponin T, slow skeletal muscle | 1.343 × 10^8^ | 0 | ∞ |
| G3V489 | **TNNI1** | Troponin I, slow skeletal muscle | 1.308 × 10^8^ | 0 | ∞ |
| P05976 | **MYL1** | Myosin light chain 1/3, skeletal muscle isoform | 1.240 × 10^8^ | 0 | ∞ |
| P35609 | **ACTN2** | Alpha-actinin-2 | 1.109 × 10^8^ | 0 | ∞ |
| A0A075B7D8 | **IGHV3OR15-7** | Immunoglobulin heavy variable 3/OR15-7 | 9.126× 10^7^ | 0 | ∞ |
| E9PJL7 | **CRYAB** | Alpha-crystallin B chain | 8.506 × 10^7^ | 0 | ∞ |
| H3BML9 | **MYL11** | Myosin regulatory light chain 2, skeletal muscle isoform | 7.248 × 10^7^ | 0 | ∞ |

| Uniprot accession | Protein | Description | Abundance | | CPS23F:CPS8 fold |
| --- | --- | --- | --- | --- | --- |
|  |  |  | CPS23F*^b^* | CPS8 |  |
| P06732 | **CKM** | Creatine kinase M-type | 6.791 × 10^7^ | 0 | ∞ |
| P05109 | **S100A8** | Protein S100-A8 | 6.667 × 10^7^ | 0 | ∞ |
| P68363 | **TUBA1B** | Tubulin alpha-1B chain | 5.891 × 10^7^ | 0 | ∞ |
| P63316 | **TNNC1** | Troponin C, slow skeletal and cardiac muscles | 5.037 × 10^7^ | 0 | ∞ |
| P68366 | **TUBA4A** | Tubulin alpha-4A chain | 5.014 × 10^7^ | 0 | ∞ |
| P02745 | **C1QA** | Complement C1q subcomponent subunit | 4.815 × 10^7^ | 0 | ∞ |
| A0A2R2Y2Q3 | **TPM3** | Tropomyosin 3 nu | 4.514 × 10^7^ | 0 | ∞ |
| G3V1V7 | **MYBPC1** | Myosin binding protein C, slow type, isoform CRA_e | 4.461 × 10^7^ | 0 | ∞ |
| P20851 | **C4BPB** | C4b-binding protein beta chain | 4.287 × 10^7^ | 0 | ∞ |
| Q13103 | **SPP2** | Secreted phosphoprotein 24 | 4.226 × 10^7^ | 0 | ∞ |
| H7C5W9 | **ATP2A2** | Endoplasmic reticulum class 1/2 Ca (2+) ATPase | 4.151 × 10^7^ | 0 | ∞ |
| P04075 | **ALDOA** | Fructose-bisphosphate aldolase A | 4.123 × 10^7^ | 0 | ∞ |
| A0A6Q8PFK8 | **HSPB1** | Heat shock protein beta-1 | 3.264 × 10^7^ | 0 | ∞ |
| P27105 | **STOM** | Stomatin OS=Homo sapiens | 2.986 × 10^7^ | 0 | ∞ |
| A0A0A0MS76 | ZNF99 | Zinc finger protein 99 | 2.774 × 10^7^ | 0 | ∞ |
| P12273 | **PIP** | Prolactin-inducible protein | 2.631 × 10^7^ | 0 | ∞ |

| Uniprot accession | Protein | Description | Abundance | | CPS23F:CPS8 fold |
| --- | --- | --- | --- | --- | --- |
|  |  |  | CPS23F*^b^* | CPS8 |  |
| H0YKX5 | **TPM1** | Tropomyosin alpha-1 chain | 2.542 × 10^7^ | 0 | ∞ |
| A0A1X7SC65 | **HSPB6** | Heat shock protein beta-6 | 2.386 × 10^7^ | 0 | ∞ |
| A0A0C4DH36 | **IGHV3-38** | Probable non-functional immunoglobulin heavy variable 3-38 | 2.348 × 10^7^ | 0 | ∞ |
| P11217 | **PYGM** | Glycogen phosphorylase, muscle form | 2.330 × 10^7^ | 0 | ∞ |
| P06727 | **APOA4** | Apolipoprotein A-IV | 2.289 × 10^7^ | 0 | ∞ |
| E5RGZ4 | ENO3 | 2-phospho-D-glycerate hydro-lyase | 2.133 × 10^7^ | 0 | ∞ |
| Q14315 | **FLNC** | Filamin-C OS=Homo sapiens | 2.071 × 10^7^ | 0 | ∞ |
| P06733 | **ENO1** | Alpha-enolase OS=Homo sapiens | 1.910 × 10^7^ | 0 | ∞ |
| A0A0G2JL69 | **C2** | C3/C5 convertase | 1.679 × 10^7^ | 0 | ∞ |
| Q8WZ42 | **TTN** | Titin OS=Homo sapiens | 1.506 × 10^7^ | 0 | ∞ |
| A0A3B3ISR2 | **C1R** | Complement subcomponent C1r | 1.319 × 10^7^ | 0 | ∞ |
| P25705 | **ATP5F1A** | ATP synthase subunit alpha, mitochondrial | 4.854 × 10^8^ | 3.279 × 10^6^ | 148.017 |
| Q6KB66 | **KRT80** | Keratin, type II cytoskeletal 80 | 4.573 × 10^8^ | 1.081 × 10^7^ | 42.301 |
| H0YF91 | **HYDIN** | Hydrocephalus-inducing protein homolog | 1.584 × 10^9^ | 4.293 × 10^7^ | 36.884 |
| A0A0A0MSG9 | SUN3 | SUN domain-containing protein 3 | 1.334 × 10^8^ | 7.284 × 10^6^ | 18.308 |
| P02748 | **C9** | Complement component C9 | 1.206 × 10^10^ | 8.664 × 10^8^ | 13.918 |

| Uniprot accession | Protein | Description | Abundance | | CPS23F:CPS8 fold |
| --- | --- | --- | --- | --- | --- |
|  |  |  | CPS23F*^b^* | CPS8 |  |
| P07360 | **C8G** | Complement component C8 gamma chain | 2.327 × 10^9^ | 1.720 × 10^8^ | 13.527 |
| Q9H2G4 | **TSPYL2** | Testis-specific Y-encoded-like protein 2 | 3.146 × 10^9^ | 2.627 × 10^7^ | 11.974 |
| P07358 | **C8B** | Complement component C8 beta chain | 2.066 × 10^9^ | 1.771 × 10^8^ | 11.662 |
| P07357 | **C8A** | Complement component C8 alpha chain | 3.728 × 10^9^ | 3.352 × 10^8^ | 11.122 |
| E9PJV1 | **GLYATL1** | Glycine N-acyltransferase-like protein 1 | 1.299 × 10^9^ | 1.302 × 10^8^ | 9.977 |
| P01031 | **C5** | Complement C5 | 3.468 × 10^9^ | 3.852 × 10^8^ | 9.01 |
| P13671 | **C6** | Complement component C6 | 2.071 × 10^9^ | 2.402 × 10^8^ | 8.623 |
| P10643 | **C7** | Complement component C7 | 2.001 × 10^9^ | 2.373 × 10^8^ | 8.43 |
| C9JGI8 | **PALS2** | MAGUK p55 subfamily member 6 | 2.038 × 10^8^ | 2.560 × 10^7^ | 7.961 |
| A0A140TA32 | **C4A** | C4a anaphylatoxin | 4.151 × 10^9^ | 5.320 × 10^8^ | 7.802 |
| A0A140TA29 | **C4B** | C4a anaphylatoxin | 4.151 × 10^9^ | 5.320 × 10^8^ | 7.802 |
| P36980 | **CFHR2** | Complement factor H-related protein 2 | 1.512 × 10^9^ | 1.985 × 10^8^ | 7.617 |
| P06702 | **S100A9** | Protein S100-A9 | 1.044 × 10^8^ | 1.384 × 10^7^ | 7.541 |
| Q03591 | **CFHR1** | Complement factor H-related protein 1 | 2.790 × 10^9^ | 3.726 × 10^8^ | 7.488 |
| P01024 | **C3** | Complement C3 | 2.253 × 10^11^ | 3.149 × 10^10^ | 7.154 |
| A0A0G2JPR0 | **C4A** | C4a anaphylatoxin | 3.709 × 10^9^ | 5.320 × 10^8^ | 6.971 |

| Uniprot accession | Protein | Description | Abundance | | CPS23F:CPS8 fold |
| --- | --- | --- | --- | --- | --- |
|  |  |  | CPS23F*^b^* | CPS8 |  |
| P12814 | **ACTN1** | Alpha-actinin-1 | 5.768 × 10^7^ | 8.496E6 | 6.789 |
| P68133 | **ACTA1** | Actin, alpha skeletal muscle | 2.460 × 10^9^ | 4.121× 10^8^ | 5.969 |
| Q9BXR6 | **CFHR5** | Complement factor H-related protein 5 | 4.829 × 10^9^ | 8.661× 10^8^ | 5.576 |
| P02671 | **FGA** | Fibrinogen alpha chain | 6.017× 10^8^ | 1.094× 10^8^ | 5.502 |
| P01599 | **IGKV1-17** | Immunoglobulin kappa variable 1-17 | 1.058× 10^8^ | 1.978 × 10^7^ | 5.350 |
| Q92496 | **CFHR4** | Complement factor H-related protein 4 | 5.265 × 10^7^ | 9.962× 10^6^ | 5.285 |
| O14791 | **APOL1** | Apolipoprotein L1 | 1.524× 10^8^ | 3.320 × 10^7^ | 4.590 |
| P04003 | **C4BPA** | C4b-binding protein alpha chain | 1.950 × 10^9^ | 4.296× 10^8^ | 4.540 |
| G5E9F8 | PROS1 | Vitamin K-dependent protein S | 9.908 × 10^7^ | 2.214 × 10^7^ | 4.475 |
| P07477 | **PRSS1** | Trypsin-1 | 1.365× 10^8^ | 3.058 × 10^7^ | 4.464 |
| P02743 | **APCS** | Serum amyloid P-component | 8.399 × 10^9^ | 1.900 × 10^9^ | 4.421 |
| P27169 | **PON1** | Serum paraoxonase/arylesterase 1 | 1.090× 10^8^ | 2.519 × 10^7^ | 4.329 |
| P0CG39 | **POTEJ** | POTE ankyrin domain family member J | 8.361× 10^8^ | 1.963× 10^8^ | 4.259 |
| P27918 | **CFP** | Properdin | 1.878× 10^10^ | 4.566 × 10^9^ | 4.112 |
| A0A0A0MS15 | **IGHV3-49** | Immunoglobulin heavy variable 3-49 | 1.796× 10^8^ | 4.401 × 10^7^ | 4.082 |
| P06312 | **IGKV4-1** | Immunoglobulin kappa variable 4-1 | 1.781 × 10^9^ | 4.664× 10^8^ | 3.819 |

| Uniprot accession | Protein | Description | Abundance | | CPS23F:CPS8 fold |
| --- | --- | --- | --- | --- | --- |
|  |  |  | CPS23F*^b^* | CPS8 |  |
| P60709 | **ACTB** | Actin, cytoplasmic 1 | 1.753 × 10^9^ | 4.630 × 10^8^ | 3.787 |
| P02747 | **C1QC** | Complement C1q subcomponent subunit C | 1.414 × 10^8^ | 3.740 × 10^7^ | 3.781 |
| Q562R1 | **ACTBL2** | Beta-actin-like protein 2 | 1.153 × 10^9^ | 3.107 × 10^8^ | 3.711 |
| P04406 | **GAPDH** | Glyceraldehyde-3-phosphate dehydrogenase | 7.976 × 10^7^ | 2.168 × 10^7^ | 3.679 |
| A0A0A0MRZ8 | **IGKV3D-11** | Immunoglobulin kappa variable 3D-11 | 5.353 × 10^9^ | 1.489 × 10^9^ | 3.595 |
| P05089 | **ARG1** | Arginase-1 OS=Homo sapiens | 1.918 × 10^7^ | 5.736 × 10^6^ | 3.343 |
| A0A0J9YX35 | **IGHV3-64D** | Immunoglobulin heavy variable 3-64D | 3.812 × 10^8^ | 1.251 × 10^8^ | 3.046 |
| Q96IY4 | **CPB2** | Carboxypeptidase B2 | 5.950 × 10^7^ | 1.991 × 10^7^ | 2.988 |
| P59665 | **DEFA1; DEFA1B** | Neutrophil defensin 1 | 7.638 × 10^7^ | 2.613 × 10^7^ | 2.924 |
| P55056 | **APOC4** | Apolipoprotein C-IV | 6.771 × 10^9^ | 2.365 × 10^9^ | 2.863 |
| P02647 | **APOA1** | Apolipoprotein A-I | 5.019 × 10^8^ | 1.788 × 10^8^ | 2.807 |
| P02675 | **FGB** | Fibrinogen beta chain | 5.329 × 10^8^ | 1.931 × 10^8^ | 2.759 |
| A0A075B6I9 | **IGLV7-46** | Immunoglobulin lambda variable 7-46 | 1.094 × 10^8^ | 4.061 × 10^7^ | 2.694 |
| Q9H4B7 | **TUBB1** | Tubulin beta-1 chain | 3.513 × 10^7^ | 1.327 × 10^7^ | 2.648 |
| P01009 | **SERPINA1** | Alpha-1-antitrypsin | 9.661 × 10^7^ | 3.678 × 10^7^ | 2.627 |
| P02765 | **AHSG** | Alpha-2-HS-glycoprotein | 3.442 × 10^7^ | 1.366 × 10^7^ | 2.520 |
| Q02413 | **DSG1** | Desmoglein-1 | 8.443 × 10^7^ | 3.388 × 10^7^ | 2.492 |
| Q13156 | **RPA4** | Replication protein A 30 kDa subunit | 4.989 × 10^8^ | 2.050 × 10^8^ | 2.434 |

| Uniprot accession | Protein | Description | Abundance | | CPS23F:CPS8 fold |
| --- | --- | --- | --- | --- | --- |
|  |  |  | CPS23F*^b^* | CPS8 |  |
| P49913 | **CAMP** | Cathelicidin antimicrobial peptide | 2.239 × 10^9^ | 9.260 × 10^8^ | 2.418 |
| P08603 | **CFH** | Complement factor H | 8.914 × 10^8^ | 3.707 × 10^8^ | 2.405 |
| P01764 | **IGHV3-23** | Immunoglobulin heavy variable 3-23 | 2.257 × 10^9^ | 9.898 × 10^8^ | 2.280 |
| A0A0J9YY99 | NA | Ig-like domain-containing protein | 2.257 × 10^9^ | 9.898 × 10^8^ | 2.280 |
| P04114 | **APOB** | Apolipoprotein B-100 | 3.842 × 10^7^ | 1.701 × 10^7^ | 2.259 |
| A0A0C4DH35 | **IGHV3-35** | Probable non-functional immunoglobulin heavy variable 3-35 | 1.567 × 10^9^ | 7.011 × 10^8^ | 2.235 |
| P01040 | **CSTA** | Cystatin-A | 6.798 × 10^7^ | 3.178 × 10^7^ | 2.139 |
| P10909 | **CLU** | Clusterin | 5.618 × 10^8^ | 2.672 × 10^8^ | 2.102 |
| P02751 | **FN1** | Fibronectin | 1.272 × 10^8^ | 6.112 × 10^7^ | 2.081 |
| P01042 | **KNG1** | Kininogen-1 | 5.250 × 10^8^ | 2.559 × 10^8^ | 2.052 |
| F5H265 | **UBC** | Polyubiquitin-C | 9.249 × 10^7^ | 4.563 × 10^7^ | 2.027 |

*^a^*Proteins were considered as CPS23F receptor candidates once meeting following criteria:

1. Annotated as receptor or binding proteins;

2. CPS23F:CPS8 enrichment fold ≥ 2.000.

*^b^*Proteins were ranked according to the protein abundance in CPS23F group.

NA, not accessible.
