## Supplemental Table S6 for "C-Reactive Protein Drives Potent Clearance of Blood Bacteria in the Liver by Activating the Complement System"

**Table S6. Cryo-EM data processing and refinement statistics**

| **Data collection and processing** | **CRP/23F complex** |
| --- | --- |
| Voltage (kV) | 300 |
| Electron exposure (e^-^/Å^2^) | 50 |
| Defocus range (μm) | -1.0~-2.0 |
| Pixel size (Å) | 0.85 |
| Number of frames collected | 32 |
| Micrographs Collected (no.) | 8.276 |
| Symmetry imposed | C5 |
| Final particles (no.) | 207,101 |
| Map resolution (Å) | 2.78 |
| FSC threshold | 0.143 |
| **Refinement** |  |
| Initial model used (PDB code) | 1B09 |
| Map sharpening methods | DeepEMhancer |
| Model composition |  |
| Non-hydrogen atoms | 8430 |
| Protein residues | 1030 |
| Ligands | 15 |
| R.m.s. deviations |  |
| Bond lengths (Å) | 0.010 |
| Bond angles (°) | 1.115 |
| Validation |  |
| MolProbity Score | 2.04 |
| Clash Score | 6.10 |
| Poor rotamers (%) | 3.00 |
| Ramachandran plot |  |
| Favored | 95.10 |
| Allowed | 4.90 |
| Disallowed | 0.00 |
